# Identification of autosomal cis expression quantitative trait methylation (cis eQTMs) in children’s blood

**DOI:** 10.1101/2020.11.05.368076

**Authors:** Carlos Ruiz-Arenas, Carles Hernandez-Ferrer, Marta Vives-Usano, Sergi Marí, Inés Quintela, Dan Mason, Solène Cadiou, Maribel Casas, Sandra Andrusaityte, Kristine Bjerve Gutzkow, Marina Vafeiadi, John Wright, Johanna Lepeule, Regina Grazuleviciene, Leda Chatzi, Ángel Carracedo, Xavier Estivill, Eulàlia Martí, Geòrgia Escaramís, Martine Vrijheid, Juan R González, Mariona Bustamante

**Author notes:** Corresponding authors: Carlos Ruiz-Arenas, Mariona Bustamante.

## Abstract

**Background:** The identification of expression quantitative trait methylation (eQTMs), defined as associations between DNA methylation levels and gene expression, might help the biological interpretation of epigenome-wide association studies (EWAS). We aimed to identify autosomal cis eQTMs in children’s blood, using data from 832 children of the Human Early Life Exposome (HELIX) project.

**Methods:** Blood DNA methylation and gene expression were measured with the Illumina 450K and the Affymetrix HTA v2 arrays, respectively. The relationship between methylation levels and expression of nearby genes (1 Mb window centered at the transcription start site, TSS) was assessed by fitting 13.6 M linear regressions adjusting for sex, age, cohort, and blood cell composition.

**Results:** We identified 39,749 blood autosomal cis eQTMs, representing 21,966 unique CpGs (eCpGs, 5.7% of total CpGs) and 8,886 unique transcript clusters (eGenes, 15.3% of total transcript clusters, equivalent to genes). In 87.9% of these cis eQTMs, the eCpG was located at <250 kb from eGene’s TSS; and 58.8% of all eQTMs showed an inverse relationship between the methylation and expression levels. Only around half of the autosomal cis-eQTMs eGenes could be captured through annotation of the eCpG to the closest gene. eCpGs had less measurement error and were enriched for active blood regulatory regions and for CpGs reported to be associated with environmental exposures or phenotypic traits. 40.4% of eQTMs had at least one genetic variant associated with methylation and expression levels. The overlap of autosomal cis eQTMs in children’s blood with those described in adults was small (13.8%), and age-shared cis eQTMs tended to be proximal to the TSS and enriched for genetic variants.

**Conclusions:** This catalogue of autosomal cis eQTMs in children’s blood can help the biological interpretation of EWAS findings and is publicly available at https://helixomics.isglobal.org/.

**Funding:** The study has received funding from the European Community’s Seventh Framework Programme (FP7/2007-206) under grant agreement no 308333 (HELIX project); the H2020-EU.3.1.2. - Preventing Disease Programme under grant agreement no 874583 (ATHLETE project); from the European Union’s Horizon 2020 research and innovation programme under grant agreement no 733206 (LIFECYCLE project), and from the European Joint Programming Initiative “A Healthy Diet for a Healthy Life” (JPI HDHL and Instituto de Salud Carlos III) under the grant agreement no AC18/00006 (NutriPROGRAM project). The genotyping was supported by the project PI17/01225, funded by the Instituto de Salud Carlos III and co-funded by European Union (ERDF, “A way to make Europe”) and the Centro Nacional de Genotipado-CEGEN (PRB2-ISCIII).

## Introduction

Cells from the same individual, although sharing the same genome sequence, differentiate into diverse lineages that finally give place to specific cell types with unique functions. This is orchestrated by the epigenome, which regulates gene expression in a cell/tissue- and time- specific manner (Cavalli and Heard, 2019; Feinberg, 2018; Lappalainen and Greally, 2017). Besides its central role in regulating embryonic and fetal development, X-chromosome inactivation, genomic imprinting, and silencing of repetitive DNA elements, the epigenome is also responsible for the plasticity and cellular memory in response to environmental perturbations (Cavalli and Heard, 2019; Feinberg, 2018; Lappalainen and Greally, 2017).

Massive epigenetic alterations, caused by somatic mutations, age, injury, or environmental exposures, were initially described in cancer (Feinberg, 2018). The paradigm of environmental factors modifying the epigenome and leading to increased disease risk was then extrapolated from cancer to a wide range of common diseases. Consequently, in recent years, a high number of epigenome-wide association studies (EWAS) have been performed, investigating the relation of prenatal and postnatal exposure to environmental factors with DNA methylation, and of DNA methylation with disease (Feinberg, 2018; Lappalainen and Greally, 2017). EWAS findings have been inventoried in two catalogues: the EWAS catalog (Battram et al., 2021) and the EWAS Atlas (Li et al., 2019). The latter includes 0.5 M associations for 498 traits from 1,216 studies, including 155 different cells/tissues.

Despite the success of EWAS in identifying altered methylation patterns, various challenging issues still must be solved: the role of genetic variation; the access to the target tissue/cell; confounding reverse causation; and biological interpretation (Feinberg, 2018; Lappalainen and Greally, 2017). Regarding the latter, most studies do not have transcriptional data to test the effect of DNA methylation on gene expression. When these data are not available, a common approach is to assume that CpG DNA methylation affects the expression of the closest gene (Sharp et al., 2017). Although this approach is easy to implement, it is limited. Indeed, CpG DNA methylation might regulate distant genes or might not regulate any gene at all (Bonder et al., 2017; Lappalainen and Greally, 2017). Another approach to elucidate the effect of DNA methylation on gene expression when transcriptional data are not available is to perform expression quantitative trait methylation (eQTM) studies. These are genome-wide studies investigating the associations between the levels of DNA methylation and gene expression (Gondalia et al., 2019; Küpers et al., 2019). Several eQTM studies have been performed in diverse cell types/tissues: whole blood (Bonder et al., 2017; Kennedy et al., 2018), monocytes (Husquin et al., 2018; Kennedy et al., 2018; Liu et al., 2013), lymphoblastoid cell lines, T-cells and fibroblasts derived from umbilical cords (Gutierrez-Arcelus et al., 2015, 2013), fibroblasts (Wagner et al., 2014), liver (Bonder et al., 2014), skeletal muscle (Leland Taylor et al., 2019), nasal airway epithelium (Kim et al., 2020), and placenta (Delahaye et al., 2018). As most of the EWAS are conducted in whole blood (Felix et al., 2018; Li et al., 2019), there is a need for comprehensive eQTM studies in this tissue. To date, available eQTM studies in whole blood only cover samples from adults (Bonder et al., 2017; Kennedy et al., 2018) and their validity in children has not been assessed.

In this study, we analyzed DNA methylation and gene expression data from the Human Early-Life Exposome (HELIX) project to an autosomal cis eQTM catalogue in children’s blood (https://helixomics.isglobal.org/). We analyzed the proportion of cis eQTMs captured through annotation to the closest gene, characterized them at the functional level, assessed the influence of genetic variation and compared them with eQTMs identified in adults. An overview of all the analyses can be found in Figure 1. This public resource will help the functional interpretation of EWAS findings in children.

**Figure 1.**
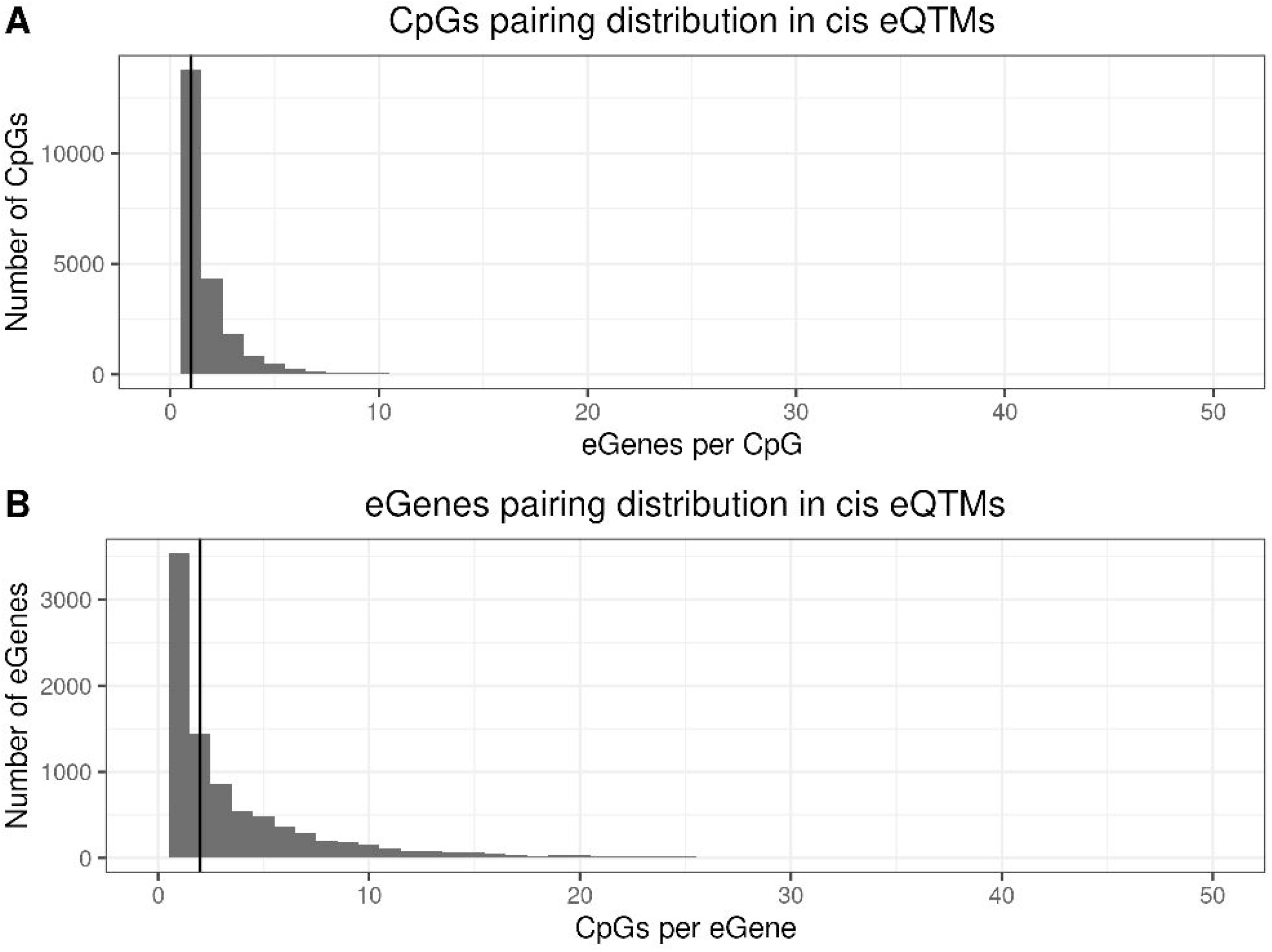
Analysis workflow. The figure summarizes the analyses conducted in this study. The first step was (1) the identification of blood autosomal cis eQTMs (1 Mb window centered at the transcription start site, TSS, of the gene) in 823 European ancestry children from the HELIX project, by a linear model adjusted for age, sex, cohort, and blood cell type proportions. All the associations are reported in the web catalogue (www.helixomics.isglobal.org). Then, (2) we explored the distance from the eCpG (CpG involved in an eQTM) to eGene’s TSS (gene involved in an eQTM), the effect size of the association, and classified eCpGs in different types. Next, (3) we evaluated the proportion of eGenes potentially inferred through annotation of eCpGs to the closest gene. Finally, (4) we functionally characterized eCpGs and eGenes; (5) assessed the contribution of genetic variants; and (6) evaluated the influence of age.

## Methods

### Sample of the study

The Human Early Life Exposome (HELIX) study is a collaborative project across 6 established and on-going longitudinal population-based birth cohort studies in Europe (Maitre et al., 2018): the Born in Bradford (BiB) study in the UK (Wright et al., 2013), the Étude des Déterminants pré et postnatals du développement et de la santé de l’Enfant (EDEN) study in France (Heude et al., 2016), the INfancia y Medio Ambiente (INMA) cohort in Spain (Guxens et al., 2012), the Kaunus cohort (KANC) in Lithuania (Grazuleviciene et al., 2009), the Norwegian Mother, Father and Child Cohort Study (MoBa)(Magnus et al., 2016) and the RHEA Mother Child Cohort study in Crete, Greece (Chatzi et al., 2017). All participants in the study signed an ethical consent and the study was approved by the ethical committees of each study area (Maitre et al., 2018).

In the present study, we selected a total of 832 children of European ancestry that had both DNA methylation and gene expression data. Ancestry was determined with cohort-specific self-reported questionnaires.

### Biological samples

DNA was obtained from buffy coats collected in EDTA tubes at mean age 8.1 years old. Briefly, DNA was extracted using the Chemagen kit (Perkin Elmer), in batches by cohort. DNA concentration was determined in a NanoDrop 1000 UV-Vis Spectrophotometer (Thermo Fisher Scientific) and with Quant-iT™ PicoGreen® dsDNA Assay Kit (Life Technologies).

RNA was extracted from whole blood samples collected in Tempus tubes (Applied Biosystems) using the MagMAX for Stabilized Blood Tubes RNA Isolation Kit (Thermo Fisher Scientific), in batches by cohort. The quality of RNA was evaluated with a 2100 Bioanalyzer (Agilent) and the concentration with a NanoDrop 1000 UV-Vis Spectrophotometer (Thermo Fisher Scientific). Samples classified as good RNA quality had an RNA Integrity Number (RIN) > 5, a similar RNA integrity pattern at visual inspection, and a concentration >10 ng/ul. Mean values for the RIN, concentration (ng/ul) and Nanodrop 260/230 ratio were: 7.05, 109.07 and 2.15, respectively.

## DNA methylation assessment

DNA methylation was assessed with the Infinium HumanMethylation450K BeadChip (Illumina), following manufacturer’s protocol at the National Spanish Genotyping Centre (CEGEN), Spain. Briefly, 700 ng of DNA were bisulfite-converted using the EZ 96-DNA methylation kit following the manufacturer’s standard protocol, and DNA methylation measured using the Infinium protocol. A HapMap sample was included in each plate. In addition, 24 HELIX inter-plate duplicates were included. Samples were randomized considering cohort, sex, and panel. Paired samples from the panel study (samples from the same subject collected at different time points) were processed in the same array. Two samples were repeated due to their overall low quality.

DNA methylation data was pre-processed using *minfi* R package (RRID:SCR_012830) (Aryee et al., 2014). We increased the stringency of the detection p-value threshold to <1e- 16, and probes not reaching a 98% call rate were excluded (Lehne et al., 2015). Two samples were filtered due to overall quality: one had a call rate <98% and the other did not pass quality control parameters of the *MethylAid* R package (RRID:SCR_002659) (van Iterson et al., 2014). Then, data was normalized with the functional normalization method with Noob background subtraction and dye-bias correction (Fortin et al., 2014b). Then, we checked sex consistency using the *shinyMethyl* R package (Fortin et al., 2014a), genetic consistency of technical duplicates, biological duplicates (panel study), and other samples making use of the genotype probes included in the Infinium HumanMethylation450K

BeadChip and the genome-wide genotyping data, when available. In total four samples were excluded, two with discordant sex and two with discordant genotypes. Batch effect (slide) was corrected using the *ComBat* R package (RRID:SCR_010974) (Johnson et al., 2007). Duplicated samples, one of the samples from the panel study and HapMap samples were removed as well as control probes, probes in sexual chromosomes, probes designed to detect Single Nucleotide Polymorphisms (SNPs) and probes to measure methylation levels at non-CpG sites, giving a final number of 386,518 probes.

CpG annotation was conducted with the *IlluminaHumanMethylation450kanno.ilmn-12.hg19* R package (Hansen, n.d.). Briefly, this package annotates CpGs to proximal promoter (200 bp upstream the TSS - TSS200), distant promoter (from 200 to 1,500 bp upstream the TSS - TSS1500), 5’UTR, first exon, gene body, and 3’UTR regions. CpGs farther than 1,500 bp from the TSS were not annotated to any gene. Relative position to CpG islands (island, shelve, shore and open sea) was also provided by the same R package.

Annotation of CpGs to 15 chromatin states was retrieved from the Roadmap Epigenomics Project web portal (RRID:SCR_008924) (https://egg2.wustl.edu/roadmap/web_portal/). Each CpG in the array was annotated to one or several chromatin states by taking a state as present in that locus if it was described in at least 1 of the 27 blood-related cell types.

### Gene expression assessment

Gene expression, including coding and non-coding transcripts, was assessed with the Human Transcriptome Array 2.0 ST arrays (HTA 2.0) (Affymetrix) at the University of Santiago de Compostela (USC), Spain. Amplified and biotinylated sense-strand DNA targets were generated from total RNA. Affymetrix HTA 2.0 arrays were hybridized according to Affymetrix recommendations using the Manual Target preparation for GeneChip Whole Transcript (WT) expression arrays and the labeling and hybridization kits. In each round, several batches of 24-48 samples were processed. Samples were randomized within each batch considering sex and cohort. Paired samples from the panel study were processed in the same batch. Two different types of control RNA samples (HeLa or FirstChoice® Human Brain Reference RNA) were included in each batch, but they were hybridized only in the first batches. Raw data were extracted with the AGCC software (Affymetrix) and stored into CEL files. Ten samples failed during the laboratory process (7 did not have enough cRNA or ss- cDNA, 2 had low fluorescence, and 1 presented an artifact in the CEL file).

Data was normalized with the GCCN (SST-RMA) algorithm at the gene level. Annotation of transcript clusters (TCs) was done with the ExpressionConsole software using the HTA-2.0 Transcript Cluster Annotations Release na36 annotation file from Affymetrix. After normalization, several quality control checks were performed and four samples with discordant sex and two with low call rates were excluded (Buckberry et al., 2014). One of the samples from the panel study was also eliminated for this analysis. Control probes and probes in sexual chromosomes or probes without chromosome information were excluded. Probes with a DABG (Detected Above Background) p-value <0.05 were considered to have an expression level different from the background, and they were defined as detected. Probes with a call rate <1% were excluded from the analysis. The final dataset consisted of 58,254 TCs.

Gene expression values were log_2_ transformed and batch effect controlled by residualizing the effect of surrogate variables calculated with the sva method (RRID:SCR_012836) (Leek et al., 2007) while protecting for main variables in the study (cohort, age, sex, and blood cellular composition).

### Blood cellular composition

Main blood cell type proportions (CD4+ and CD8+ T-cells, natural killer cells, monocytes, eosinophils, neutrophils, and B-cells) were estimated using the Houseman algorithm (Houseman et al., 2012) and the Reinius reference panel (Reinius et al., 2012) from raw methylation data.

### Genome-wide genotyping

Genome-wide genotyping was performed using the Infinium Global Screening Array (GSA) MD version 1 (Illumina), which contains 692,367 variants, at the Human Genomics Facility (HuGe-F), Erasmus MC, The Netherlands. Genotype calling was done using the GenTrain2.0 algorithm based on a custom cluster file implemented in the GenomeStudio software (RRID:SCR_010973). Annotation was done with the GSAMD-24v1- 0_20011747_A4 manifest. Samples were genotyped in two rounds, and 10 duplicates were included which confirmed high inter-round consistency.

Quality control was performed with the PLINK program (RRID:SCR_001757) following standard recommendations (Chang et al., 2015; Purcell et al., 2007). We applied the following sample quality controls: sample call rate <97% (N filtered=43), sex concordance (N=8), heterozygosity based on >4 SD (N=0), relatedness with PI_HAT >0.185 (N=10, including potential DNA contamination), duplicates (N=19). Then, we used the *peddy* tool (RRID:SCR_017287) to predict ancestry from GWAS data (Pedersen and Quinlan, 2017). We contrasted ancestry predicted from GWAS with ancestry recorded in the questionnaires. Twelve samples were excluded due to discordances between the two variables. Overall, 93 (6.7%) samples, including the duplicates, were filtered out. The variant quality control included the following steps: variant call rate <95% (N filtered=4,046), non-canonical PAR (N=47), minor allele frequency (MAF) <1% (N=178,017), Hardy-Weinberg equilibrium (HWE) p-value <1e-06 (N=913). Some other SNPs were filtered out during the matching between data and reference panel before imputation (N=14,436).

Imputation of the GWAS data was performed with the Imputation Michigan server (RRID:SCR_017579) (Das et al., 2016) using the Haplotype Reference Consortium (HRC) cosmopolitan panel, Version r1.1 2016 (McCarthy et al., 2016). Before imputation, PLINK GWAS data was converted into VCF format and variants were aligned with the reference genome. The phasing of the haplotypes was done with Eagle v2.4 (RRID:SCR_017262) (Loh et al., 2016) and the imputation with minimac4 (RRID:SCR_009292) (Fuchsberger et al., 2015), both implemented in the code of the Imputation Michigan server. In total, we retrieved 40,405,505 variants after imputation. Then, we applied the following QC criteria to the imputed dataset: imputation accuracy (R2) >0.9, MAF >1%, HWE p-value >1e-06; and genotype probabilities were converted to genotypes using the best guest approach. The final post-imputation quality-controlled dataset consisted of 1,304 samples and 6,143,757 variants (PLINK format, Genome build: GRCh37/hg19, + strand).

### Identification of autosomal cis eQTMs in children’s blood

To test associations between DNA methylation levels and gene expression levels in cis (cis eQTMs), we paired each Gene to CpGs closer than 500 kb from its TSS, either upstream or downstream. For each Gene, the TSS was defined based on HTA-2.0 annotation, using the start position for transcripts in the + strand, and the end position for transcripts in the - strand. CpGs position was obtained from Illumina 450K array annotation. Only CpGs in autosomal chromosomes (from chromosome 1 to 22) were tested. In the main analysis, we fitted for each CpG-Gene pair a linear regression model between gene expression and methylation levels adjusted for age, sex, cohort, and blood cell type composition. A second model was run without adjusting for blood cellular composition and it is only reported on the online web catalog, but not discussed in this manuscript. Although some of the unique associations of the unadjusted model might be real, others might be confounded by the large methylation and expression changes among blood cell types.

To ensure that CpGs paired to a higher number of Genes do not have higher chances of being part of an eQTM, multiple-testing was controlled at the CpG level, following a procedure previously applied in the Genotype-Tissue Expression (GTEx) project (Gamazon et al., 2018). Briefly, our statistic used to test the hypothesis that a pair CpG-Gene is significantly associated is based on considering the lowest p-value observed for a given CpG and all its paired Gene (e.g., those in the 1 Mb window centered at the TSS). As we do not know the distribution of this statistic under the null, we used a permutation test. We generated 100 permuted gene expression datasets and ran our previous linear regression models obtaining 100 permuted p-values for each CpG-Gene pair. Then, for each CpG, we selected among all CpG-Gene pairs the minimum p-value in each permutation and fitted a beta distribution that is the distribution we obtain when dealing with extreme values (e.g. minimum) (Dudbridge and Gusnanto, 2008). Next, for each CpG, we took the minimum p-value observed in the real data and used the beta distribution to compute the probability of observing a lower p-value. We defined this probability as the empirical p-value of the CpG. Then, we considered as significant those CpGs with empirical p-values to be significant at 5% false discovery rate using Benjamini-Hochberg method. Finally, we applied a last step to identify all significant CpG-Gene pairs for all eCpGs. To do so, we defined a genome-wide empirical p-value threshold as the empirical p-value of the eCpG closest to the 5% false discovery rate threshold. We used this empirical p-value to calculate a nominal p-value threshold for each eCpG, based on the beta distribution obtained from the minimum permuted p-values. This nominal p-value threshold was defined as the value for which the inverse cumulative distribution of the beta distribution was equal to the empirical p-value. Then, for each eCpG, we considered as significant all eCpG-Gene variants with a p-value smaller than nominal p-value.

### Characterization of the child blood autosomal cis eQTM catalogue

Wilcoxon tests were run to compare continuous variables (e.g., methylation range, CpG probe reliability, etc.) vs. categorical variables (e.g., low, medium, and high categories of methylation levels, eCpGs vs non eCpGs, etc.). We run a linear model to test the association between the effect size and the distance between the CpG and the Gene’s TSS. For this test, we compared the absolute value of the effect size vs log_10_ of absolute value of the distance, *Enrichment of eCpGs for regulatory elements:* Enrichment of eQTMs for regulatory elements were tested using Chi-square tests with non eQTMs as reference, unless otherwise stated. Results with a p-value <0.05 were considered statistically significant. Annotation of eQTMs to regulatory elements (gene relative positions, CpG island relative positions, and blood ROADMAP chromatin states) is described in the section “DNA methylation assessment”. Enrichment for CpGs classified in 3 groups based on their median methylation levels (low: 0.0-0.3; medium: >0.3-0.7; and high: >0.7-1.0) was tested similarly.

*Enrichment of eCpGs for CpGs associated with phenotypic traits and exposures:* We also explored the enrichment of eQTMs for phenotypic traits and/or environmental exposures reported in the EWAS catalog (Battram et al., 2021) and the EWAS Atlas (Li et al., 2019). We used version 03-07-2019 of the EWAS catalog and selected those studies conducted in whole or peripheral blood of European ancestry individuals. We downloaded EWAS Atlas data on 27-11-2019 and selected those studies performed in whole blood or peripheral blood of European ancestry individuals or with unreported ancestry. Enrichment was tested as indicated above.

*Enrichment of eCpGs for age-variable CpGs:* We used results from the MeDALL and the Epidelta projects to test whether eQTMs were enriched for CpGs variable from birth to childhood and adolescence. For MeDALL we downloaded data from supplementary material of the following manuscript that assesses changes from 0 to 4y and from 4y to 8y (Xu et al., 2017). For Epidelta, we downloaded the full catalogue (version 2020-07-17) from their website (http://epidelta.mrcieu.ac.uk/). In Epidelta, we considered a CpG as age-variable if its p-value from model 1 that assesses linear changes from 0 to 17 years (variable M1.change.p) was <1e-7 (Bonferroni threshold as suggested in the study). Variable CpGs were classified as increased methylation if their change estimate (variable M1.change.estimate) was >0, and as decreased methylation, otherwise. Enrichment was tested as indicated above.

*Enrichment of eGenes for Gene Ontology - Biological Processes (GO-BP):* We also tested whether eGenes were enriched for specific GO-BP terms using the *topGO* R package (RRID:SCR_014798) (J, 2010) and using the genes annotated by Affymetrix in our dataset as background (58,254 Genes annotated to 23,054 Gene Symbols). We applied the *weight01* algorithm, which considers GO-BP terms hierarchy for p-values computation. GO- BP terms with q-value <0.001 were considered statistically significant.

### Comparison of genes associated with eQTMs versus annotation of eQTMs to the closest gene

We evaluated whether genes associated with eQTMs could be captured through the Illumina annotation, which links CpGs to the closest gene in a maximum distance of 1500 bp. For this, CpGs were annotated to Gene Symbols using the *IlluminaHumanMethylation450kanno.ilmn-12.hg19* R package (Hansen, n.d.), while Genes were annotated to Gene Symbols using the HTA-2.0 Transcript Cluster Annotations Release na36 annotation file from Affymetrix. Given that CpGs and Genes could be annotated to several genes, we considered that a CpG-Gene pair was annotated to the same gene if at least one of the genes annotated to the CpG was present among the genes in the HTA-2.0 array. In total, we identified 327,931 CpG-Gene pairs annotated to the same gene, and thus that could be compared. Then, a Chi-square test was applied to compute whether eQTMs were enriched for these 327,931 comparable CpG-Gene pairs, using as background all 13M CpG-Gene pairs.

Next, we evaluated whether the relative position of the CpG in the genic region was related to the expression of the paired Gene. To do so, the comparable 327,931 CpG-Gene pairs were expanded to 383,672 entries. Each entry represented a CpG-Gene pair annotated to a unique gene relative position. Thus, for instance, a CpG-Gene pair with the CpG annotated to two relative gene positions of the same gene was included as two entries, each time annotated to a different gene relative position. In this expanded CpG-Gene pair set, Chi-square tests were run to test the enrichment of eQTMs for gene relative positions, using the 383,672 entries as background.

### Evaluation of the genetic contribution on child blood autosomal cis eQTMs

We used two approaches to evaluate the influence of genetic effects in child blood autosomal cis eQTMs. First, we analyzed heritability estimates of CpGs computed by Van Dongen and colleagues (van Dongen et al., 2016). Total additive and SNP-heritabilities were compared between eCpGs and non eCpGs, using a Wilcoxon test. We also run linear regressions between heritability measures (outcome) and eCpGs classified according to the number of eGenes they were associated with.

Second, we tested whether eCpGs were more likely regulated by SNPs than non eCpGs (i.e., whether they were enriched for meQTL). In order to define meQTLs in HELIX, we selected 9.9 M cis and trans meQTLs with a p-value <1e-7 in the ARIES dataset consisting of data from children of 7 years old (Gaunt et al., 2016). Then, we tested whether this subset of 9.9 M SNPs were also meQTLs in HELIX by running meQTL analyses using *MatrixEQTL* R package (Shabalin, 2012), adjusting for cohort, sex, age, blood cellular composition and the first 20 principal components (PCs) calculated from genome-wide genetic data of the GWAS variability. We confirmed 2.8 M meQTLs in HELIX (p-value <1e-7). Trans meQTLs represented <10% of the 2.8 M meQTLs. Enrichment of eCpGs for meQTLs was computed using a Chi-square test, using non eCpGs as background.

Finally, we tested whether meQTLs were also eQTLs for the eGenes linked to the eCpGs. To this end, we run eQTL analyses (gene expression being the outcome and 2.8 M SNPs the predictors) with *MatrixEQTL* adjusting for cohort, sex, age, blood cellular composition and the first 20 GWAS PCs in HELIX. We considered as significant eQTLs the SNP-Gene pairs with p-value <1e-7 and with the direction of the effect consistent with the direction of the meQTL and the eQTM.

### Comparison with adult blood eQTM catalogues: GTP and MESA

We compared our list of child blood autosomal cis eQTMs obtained in HELIX with the cis and trans eQTMs described in blood of two adult cohorts: GTP and MESA (Kennedy et al., 2018). DNA methylation was assessed with the Infinium HumanMethylation450K BeadChip (Illumina) in the 3 cohorts. In HELIX, gene expression was assessed with the Human Transcriptome Array 2.0 ST arrays (HTA 2.0) (Affymetrix), and in GTP and MESA with the HumanHT-12 v3.0 and v4.0 Expression BeadChip (Illumina).

For the comparison of eQTMs between adults and children, eGenes in the two studies were annotated to a common gene nomenclature, by using the Gene Symbol annotation provided by the authors form GTP and MESA, and the Gene Symbol provided by the Affymetrix annotation in HELIX. Some eQTMs involved Genes (HELIX) or gene probes (GTP and MESA) annotated to more than one gene (Gene Symbol); and also different Genes (HELIX) or gene probes (GTP and MESA) were annotated to the same Gene Symbol. To handle this issue, we split our comparison in two analyses.

First, we checked whether CpG-gene pairs reported in GTP and MESA were eQTMs (significant CpG-gene pairs) in HELIX. By doing this, the comparison was restricted to cis effects (as HELIX only considered cis effects). When a CpG-gene pair in GTP or MESA mapped to multiple CpG-gene pairs in HELIX, we only considered the CpG-gene pair with the smallest p-value in HELIX. Next, Pearson’s correlations between the effect sizes of the different studies were computed.

Second, we explored whether HELIX eQTMs were also present in GTP and/or MESA. When a CpG-gene pair in HELIX mapped to multiple CpG-gene pairs in GTP and/or MESA, we only considered the CpG-gene pair with the smallest p-value in these cohorts. As a result, HELIX eQTMs were classified in age-shared (if present in adults at p-value <1e-05, in GTP and/or MESA) and children-specific (absent in adult cohorts). For these two subsets of eQTMs, enrichment for ROADMAP chromatin states, methylation measurement error, and distance from the eCpG to the eGene’s TSS, was tested as explained above.

### Data and software availability

The raw data used to generate the eQTM catalogue are not publicly available due to privacy restrictions but are available from the corresponding author on request. Catalogue of eQTMs described in this manuscript is publicly available at https://helixomics.isglobal.org/. Scripts to reproduce the analysis can be found in a public GitHub repository (https://github.com/yocra3/methExprsHELIX/) and as a supplementary file.

## Results

### Study population and molecular data

The study includes 823 children of European ancestry from the HELIX project with available blood DNA methylation and gene expression data. These children, enrolled in 6 cohorts, were aged between 6 and 11 years and the number of males and females was balanced (Table 1).

**Table 1.** Descriptive of the study population. BiB: Born in Bradford study (UK). EDEN: Étude des Déterminants pré et postnatals du développement et de la santé de l’Enfant (France). KANC: Kaunus cohort (Lithuania). MoBa: Norwegian Mother, Father and Child Cohort Study (Norway). RHEA: Mother Child Cohort study (Greece). INMA: INfancia y Medio Ambiente cohort (Spain).

| Variable | BiB | EDEN | KANC | MoBa | RHEA | INMA | All |
| --- | --- | --- | --- | --- | --- | --- | --- |
| N (%) | 80 (9.7%) | 80 (9.7%) | 143 (17.4%) | 188 (22.9%) | 154 (18.7%) | 178 (21.6%) | 823 (100%) |
| Female (%) | 36 (45%) | 35 (43.8%) | 64 (44.8%) | 88 (46.8%) | 69 (44.8%) | 80 (44.9%) | 372 (45.2%) |
| Male (%) | 44 (55%) | 45 (56.2%) | 79 (55.2%) | 100 (53.2%) | 85 (55.2%) | 98 (55.1%) | 451 (54.8%) |
| Age, in years (IQR) | 6.65 (6.44-6.84) | 10.76 (10.37-11.22) | 6.40 (6.12-6.88) | 8.53 (8.17-8.83) | 6.45 (6.36-6.62) | 8.84 (8.44-9.21) | 8.06 (6.49-8.86) |
| Natural Killer cells (IQR) | 0.01 (0.00-0.03) | 0.02 (0.00-0.04) | 0.04 (0.01-0.07) | 0.02 (0.00-0.07) | 0.01 (0.00-0.03) | 0.03 (0.01-0.05) | 0.02 (0.00-0.05) |
| B-cell (IQR) | 0.12 (0.11-0.15) | 0.09 (0.07-0.11) | 0.11 (0.09-0.13) | 0.11 (0.09-0.14) | 0.14 (0.11-0.16) | 0.10 (0.08-0.13) | 0.11 (0.09-0.14) |
| CD4+ T-cell (IQR) | 0.21 (0.18-0.25) | 0.16 (0.14-0.20) | 0.17 (0.14-0.21) | 0.21 (0.17-0.25) | 0.20 (0.16-0.26) | 0.17 (0.14-0.21) | 0.19 (0.15-0.23) |
| CD8+ T-cell (IQR) | 0.13 (0.11-0.17) | 0.11 (0.08-0.13) | 0.13 (0.10-0.16) | 0.14 (0.11-0.17) | 0.14 (0.11-0.16) | 0.12 (0.09-0.14) | 0.13 (0.10-0.16) |
| Monocytes (IQR) | 0.09 (0.07-0.10) | 0.09 (0.07-0.11) | 0.08 (0.06-0.09) | 0.08 (0.07-0.10) | 0.09 (0.07-0.10) | 0.09 (0.07-0.11) | 0.08 (0.07-0.10) |
| Granulocytes (IQR) | 0.41 (0.35-0.47) | 0.52 (0.47-0.56) | 0.46 (0.40-0.53) | 0.41 (0.32-0.48) | 0.41 (0.34-0.48) | 0.48 (0.42-0.55) | 0.44 (0.37-0.52) |

Continuous variables are expressed as mean and interquartile range (IQR).

After quality control, our dataset consists of 386,518 CpGs and 58,254 transcript clusters (TCs) in autosomal chromosomes (from 1 to 22). TCs are defined as groups of one or more probes covering a region of the genome, reflecting all the exonic transcription evidence known for the region, and corresponding to a known or putative gene. Thus, we will refer TCs to Genes indistinctively. According to Affymetrix annotation, 23,054 of the Genes encoded a protein. To detect cis effects, we paired each Gene to all CpGs closer than 0.5 Mb from its transcription start site (TSS), either upstream or downstream (1 Mb window centered at the TSS). In total, we obtained 13.6 M CpG-Gene pairs, where each CpG was paired to a median of 30 Genes; and each Gene was paired to a median of 162 CpGs (Figure 1 – figure supplement 1).

### Identification of autosomal cis eQTMs in children’s blood

We tested the association between DNA methylation and gene expression levels in the 13.6 M autosomal CpG-Gene pairs through linear regressions adjusting for sex, age, cohort, and cellular composition. After correcting for multiple testing (see Material and Methods), we identified 39,749 statistically significant autosomal cis eQTMs in children’s blood (0.29% of total CpG-Gene pairs). These eQTMs comprised 21,966 unique CpGs (5.7% of total CpGs) and 8,886 unique Genes (15.3% of total Genes), of which 6,288 were annotated as coding genes. For simplicity, we will refer to them as eQTMs (statistically significant associations of CpG-Gene pairs), eCpGs (CpGs involved in eQTMs), and eGenes (Genes involved in eQTMs). 23,355 eQTMs (58.8% of total) showed inverse associations, meaning that higher DNA methylation was associated with lower gene expression. In eQTMs, each eGene was associated with a median of 2 eCpGs, while each eCpG was associated with a median of 1 eGene (Figure 1 – figure supplement 2). eCpGs presented higher methylation variability in the population (Figure 1 – figure supplement 3), and were measured with lower technical error (Sugden et al., 2020) (Figure 1 – figure supplement 4). Indeed, 13,278 eCpGs (60.4% of total) were measured with probes which had an intraclass correlation coefficient (ICC) >0.4, which is indicative of reliable measurements. Moreover, eGenes had higher call rates (Figure 1 – figure supplement 5).

The complete catalogue of eQTMs can be downloaded from https://helixomics.isglobal.org/.

### Overview of autosomal cis eQTMs in children’s blood

#### Distance from the eCpG to the eGene’s TSS and effect size

eCpGs tended to be close to the TSS of the targeted eGenes, being this distance <250Kb for 87.9% of all eQTMs (Figure 2A). Globally, the median distance between an eCpG and the TSS of its associated eGene was 1.1 kb (IQR = -33 kb; 65 kb), being eCpGs closer to the TSS in inverse eQTMs than in positive. The observed downstream shift could be explained because we chose the most upstream TSS for each Gene according to the Affymetrix HTAv2 annotation. A similar shift was observed for expression quantitative trait loci, eQTLs, (i.e., single nucleotide polymorphisms, SNPs, associated with gene expression) in the Genotype-Tissue Expression (GTEx) project (Gamazon et al., 2018).

**Figure 2.**
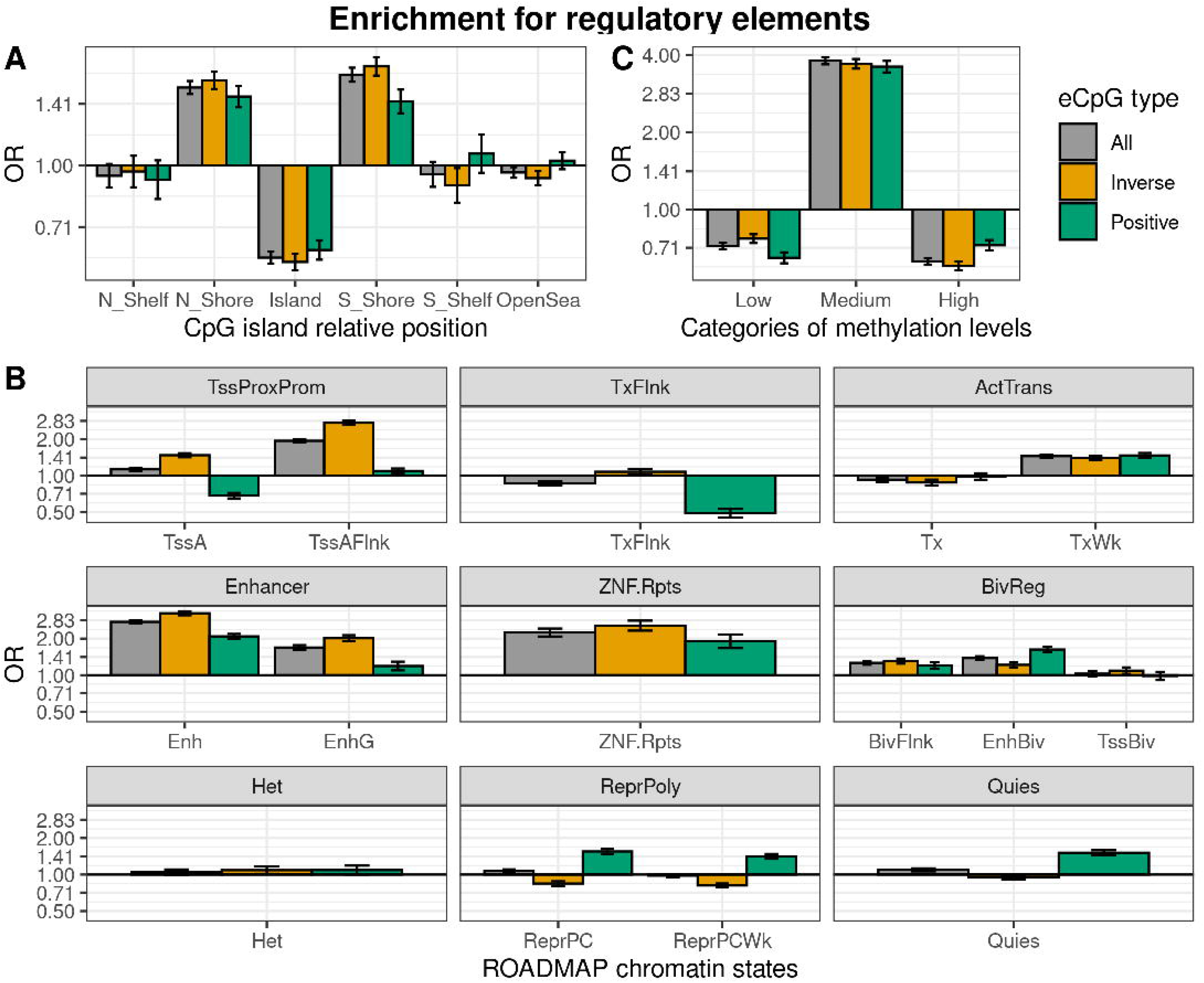
Distance between CpG and Gene’s TSS and effect size in child blood autosomal cis eQTMs. A) Distribution of the distance between CpG and Gene’s TSS by eQTM type. CpG-Gene pairs were classified in non eQTMs (black); inverse eQTMs (yellow); and positive eQTMs (green). The x-axis represents the distance between the CpG and the Gene’s TSS (kb). Non eQTMs median distance: -0.013 kb (interquartile range - IQR = -237; 236). Positive eQTMs median distance: -4.9 kb (IQR = -38; 79). Inverse eQTMs median distance: -0.7 kb (IQR = -29; 54). B) Effect size versus eCpG- Gene’s TSS distance in eQTMs. The x-axis represents the distance between the eCpG and the eGene’s TSS (kb). The y-axis represents the effect size as the log2 fold change in gene expression produced by a 0.1 increase in DNA methylation (or 10 percentile increase). To improve visualization, a 99% winsorization has been applied to log2 fold change values: values more extreme than 99% percentile (in absolute value) have been changed for the 99% quantile value (in absolute value). eQTMs are classified in inverse (yellow) and positive (green). Each eQTM is represented by one dot. The darker the color, the more dots overlapping, and so the higher the number of eQTMs with the same effect size and eCpG-eGene’s TSS distance.

We report the effect size of eQTMs as the log_2_ fold change (FC) of gene expression per 0.1 points increase in methylation (or 10 percentile increase). In absolute terms, the median effect size was 0.12, being the minimum 0.002 and the maximum 16.0, with 96.3% of the eQTMs with an effect size <0.5. A median effect size of 0.12 means that a change of 0.1 points in methylation levels was associated with around a 9% increase/decrease of gene expression. We observed an inverse linear association between the eCpG-eGene’s TSS distance and the effect size (p-value = 7.75e-9, Figure 2B); while we did not observe significant differences in effect size due to the relative orientation of the eCpG (upstream or downstream) with respect to the eGene’s TSS (p-value = 0.68).

#### Classification of eCpGs

As shown in Table 2, we classified eCpGs into 5 types, by following 2 criteria: (1) the number of eGenes affected, distinguishing between mono eCpGs (associated with a unique eGene), and multi eCpGs (associated with >= 2 eGenes); and (2) the direction of the effect, distinguishing between inverse, positive and bivalent eCpGs (with inverse effects on some eGenes and positive effects on others). Mono inverse eCpGs were the most abundant type (36.8%) (Table 2). CpGs not associated with the expression of any Gene were named as non eCpGs. We used these categories in the subsequent analyses.

**Table 2.** Classification of eCpGs by type. Percentages refer to the total number of eCpGs.

|  | Inverse (N, %) | Positive (N, %) | Bivalent (N, %) | Total (N, %) |
| --- | --- | --- | --- | --- |
| Mono | 8,084 (36.8%) | 5,681 (25.9%) | 0, by definition | 13,765 (62.7%) |
| Multi | 3,738 (17.0%) | 2,400 (10.9%) | 2,063 (9.4%) | 8,201 (37.3%) |
| Total | 11,822 (53.8%) | 8,081 (36.8%) | 2,063 (9.4%) | 21,966 (100%) |

### Comparison of eGenes with the closest annotated gene

A standard approach to interpret EWAS findings is to assume that a CpG regulates the expression of proximal genes. These genes are usually identified through the Illumina 450K annotation (Hansen, n.d.), which annotates a CpG to a gene when the CpG maps into the gene body, untranslated, or promoter region defined as <1,500 bp upstream the TSS. We evaluated to which extent the Illumina 450K annotation captured the eQTMs identified in our catalogue.

First, we observed that CpG-Gene pairs where CpG and Gene were annotated to the same Gene Symbol were more likely eQTMs than CpG-Gene pairs annotated to different Gene Symbols or without gene annotation (OR = 11.90, p-value <2e-16). Next, we assessed whether the gene annotated to the eCpG with the Illumina 450K annotation was coincident with the eGene found in our analysis. To answer this, we selected 14,797 eCpGs (67.4% of total eCpGs) annotated to Gene Symbols also present in the Affymetrix array, and thus comparable. In 7,808 out of these 14,797 eCpGs, the eCpG was associated with the expression of an eGene coincident with at least one of the Gene Symbols in Illumina’s annotation (52.8% of eCpGs with comparable gene annotation, 35.5% of all eCpGs).

Finally, we explored whether the relative gene position of a CpG determines its association with gene expression. We selected the 327,931 CpG-Gene pairs with the CpG and Gene annotated to the same Gene Symbol. Within this subset, eCpGs were enriched for CpGs in 5’UTRs and gene body positions, while depleted for CpGs in proximal promoters and 3’UTRs (Figure 2 – figure supplement 1). Interestingly, we observed that inverse and positive eCpGs were enriched for CpGs located in different gene regions: inverse for CpGs in distal promoters (TSS1500) and 5’UTRs; positive for CpGs in gene bodies.

Overall, only around half of the eGenes targeted by the eQTMs could be identified by the Illumina 450K annotation. We also found that while eCpGs were enriched for TSS1500, 5’UTRs, and gene body positions.

### Functional characterization of autosomal cis eQTMs in children’s blood

#### Enrichment of eCpGs for genomic regulatory elements

We characterized eCpGs by evaluating their enrichment for diverse regulatory elements, including CpG island relative positions and 15 chromatin states retrieved from 27 blood cell types from the ROADMAP Epigenomics project (Roadmap Epigenomics Consortium et al., 2015). First, we found that eCpGs were depleted for CpG islands, while mostly enriched for CpG island shores, but also for shelves and open sea (Figure 3A). We did not observe relevant differences between inverse and positive eCpGs.

**Figure 3.**
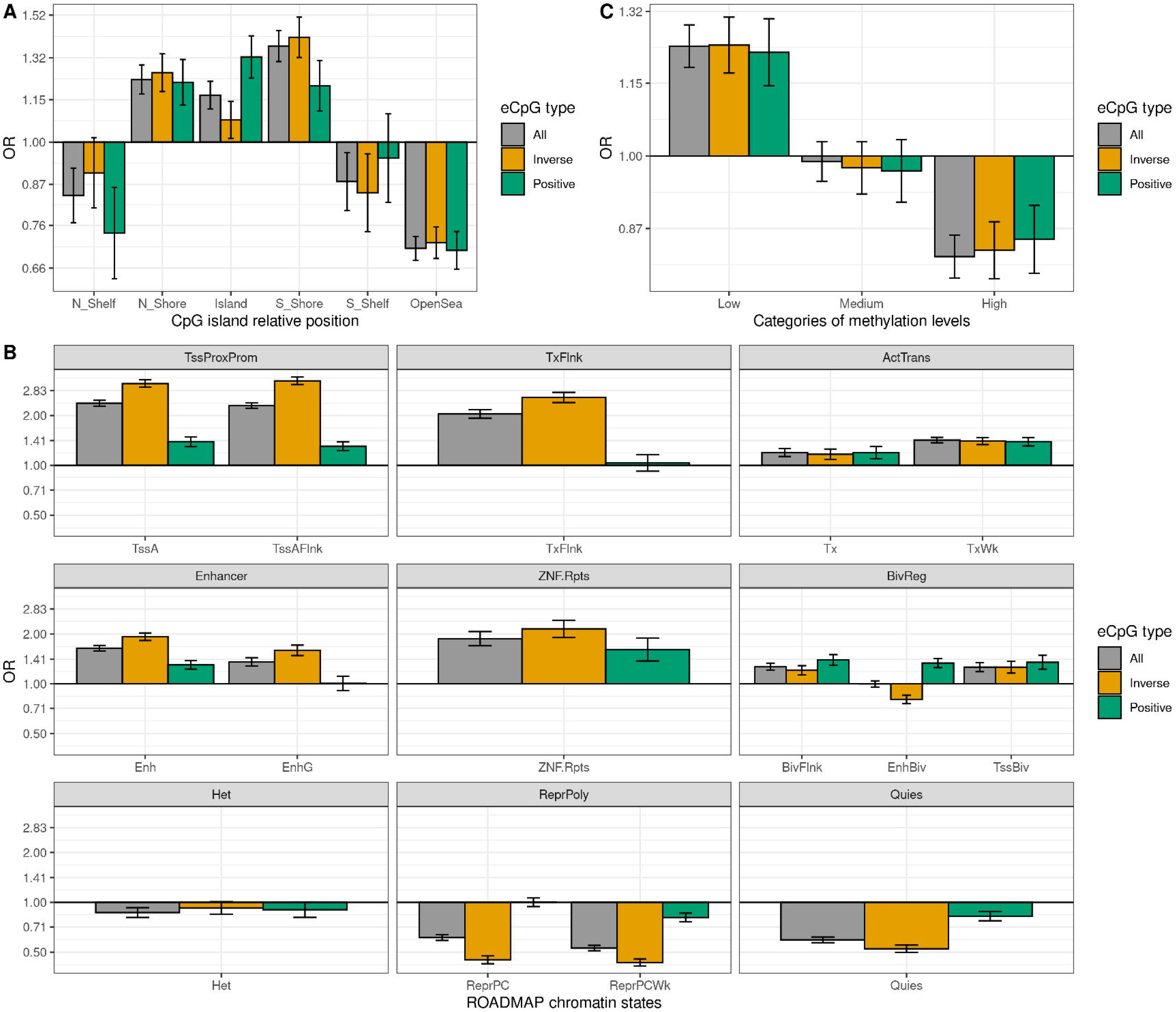
Enrichment of cis autosomal eCpGs in children’s blood for different regulatory elements. eCpGs were classified in all (grey), inverse (yellow), and positive (green). The y-axis represents the odds ratio (OR) of the enrichment. In all cases, the enrichment was computed against non eCpGs. A) Enrichment for CpG island relative positions: CpG island, N- and S-shore, N- and S- shelf, and open sea. B) Enrichment for ROADMAP blood chromatin states (Roadmap Epigenomics Consortium et al., 2015): active TSS (TssA); flanking active TSS (TssAFlnk); transcription at 5’ and 3’ (TxFlnk); transcription region (Tx); weak transcription region (TxWk); enhancer (Enh); genic enhancer (EnhG); zinc finger genes and repeats (ZNF.Rpts); flanking bivalent region (BivFlnx); bivalent enhancer (EnhBiv); bivalent TSS (TssBiv); heterochromatin (Het); repressed Polycomb (ReprPC); weak repressed Polycomb (ReprPCWk); and quiescent region (Quies). Chromatin states can be grouped in active transcription start site proximal promoter states (TssProxProm), active transcribed states (ActTrans), enhancers (Enhancers), bivalent regulatory states (BivReg), and repressed Polycomb states (ReprPoly). C) Enrichment for categories of CpGs with different median methylation levels: low (0-0.3), medium (0.3-0.7), and high (0.7-1) (Huse et al., 2015).

Second, we assessed whether eCpGs were enriched for ROADMAP blood chromatin states (Roadmap Epigenomics Consortium et al., 2015) (Figure 3B). eCpGs were enriched for several active states, such as enhancers or active transcription regions. Nonetheless, we observed some discrepancies between eCpGs subtypes: only inverse eCpGs were enriched for proximal promoter states while only positive eCpGs were depleted for transcription at 5’ and 3’ (TxFlnk). In inactive chromatin states, both positive and inverse eCpGs were enriched for bivalent regulatory states (BivReg), while only positive eCpGs were enriched for repressed and weak repressed Polycomb regions (ReprPC, ReprPCWk) and quiescent regions (Quies).

Third, we also analyzed whether eCpGs had different methylation levels. We found that eCpGs were enriched for CpGs with medium (>0.3-0.7) methylation levels and depleted for CpGs with low (0-0.3) or high (>0.7-1) methylation levels (Figure 3C).

Finally, we wondered whether these enrichments could be affected by the bias introduced by methylation measurement error; thus, we repeated all the enrichment analyses only considering 75,836 CpGs measured with reliable probes (ICC >0.4) (Sugden et al., 2020) (Figure 3 – figure supplement 1). After this filtering, the enrichments for CpG island relative positions and for categories of CpGs according to their methylation levels changed substantially: eCpGs passed from being depleted to being enriched for CpG island positions (Figure 3 – figure supplement 1A), and from being enriched for CpGs with medium methylation levels to being enriched for CpGs with low methylation levels (Figure 3 – figure supplement 1C). On the contrary, the magnitudes of enrichments for most of the active chromatin states were increased (Figure 3 – figure supplement 1B); while enrichments of positive eCpGs for inactive states (ReprPoly and Quies) were reverted. Overall, selecting reliable CpG probes reduced the differences between inverse and positive eCpGs and resulted in enrichments for active chromatin states and depletions for inactive states.

#### Gene-set enrichment analysis

To identify which biological functions were regulated by our list of eQTMs, we ran gene-set enrichment analyses using the list of eGenes. 5,503 out of the 8,886 unique Gene Symbols annotated to eGenes were present in Gene Ontology - Biological Processes (GO-BP), leading to 52 enriched terms (q-value <0.001) (Table S1). As expected from the tissue analyzed, 50% of the terms were related to immune responses (N = 26), followed by terms associated with cellular (N = 16) and metabolic (N = 10) processes. Among immune terms, 9 of them were part of innate immunity, 9 of adaptive response, and 8 were related to general/other immune pathways. Most enriched GO-BP terms were also found when running the enrichment with the list of eGenes derived from eQTMs measured with reliable CpG probes (ICC >0.4) (Table S1).

#### Enrichment for CpGs reported in the EWAS catalogues

We assessed whether eCpGs were enriched for CpGs previously related to phenotypic traits and/or environmental exposures. To this end, we retrieved CpGs from EWAS performed in blood of European ancestry subjects: 143,384 CpGs from the EWAS catalog (Battram et al., 2021), and 54,599 CpGs from the EWAS Atlas (Li et al., 2019). We found that eCpGs were enriched for CpGs in these EWAS databases in comparison to non eCpGs. Although we observed larger odds ratios (ORs) for CpGs listed in the EWAS Atlas than for CpGs in the EWAS Catalog (Figure 3 – figure supplement 2A), this difference disappeared after removing CpGs with less reliable measurements (ICC <0.4) (Figure 3 – figure supplement 2B).

### Genetic contribution to autosomal cis eQTMs in children’s blood

#### Additive and SNP heritability of eQTMs

We hypothesized that genetic variation might regulate DNA methylation and gene expression in some of the autosomal cis eQTMs in children’s blood. To test this, we used two measures of genetic influence: (1) heritability of blood DNA methylation levels for each CpG, calculated from twin designs (total additive heritability) and from genetic relationship matrices (SNP heritability), as reported by Van Dongen and colleagues (van Dongen et al., 2016); and (2) methylation quantitative trait loci (meQTLs, SNPs associated with DNA methylation levels) identified in the ARIES dataset (Gaunt et al., 2016).

First, we found that eCpGs had higher total additive and SNP heritabilities than non eCpGs (median difference of 0.31 and 0.11, respectively, p-value <2e-16 for both). Moreover, total additive and SNP heritabilities were higher for eCpGs associated with a larger number of eGenes (increase of 0.025 and 0.026 points per eGene, respectively, with a p-value <2e-16 for both) (Figure 4A and 4B). After removing CpG probes with unreliable measurements (ICC <0.4), differences in median total additive heritability between eCpGs and non eCpGs were still present, but smaller (0.15, p-value <2e-16); whereas differences in SNP heritabilities were maintained (0.11, p-value <2e-16) (Figure 4 – figure supplement 1).

**Figure 4.**
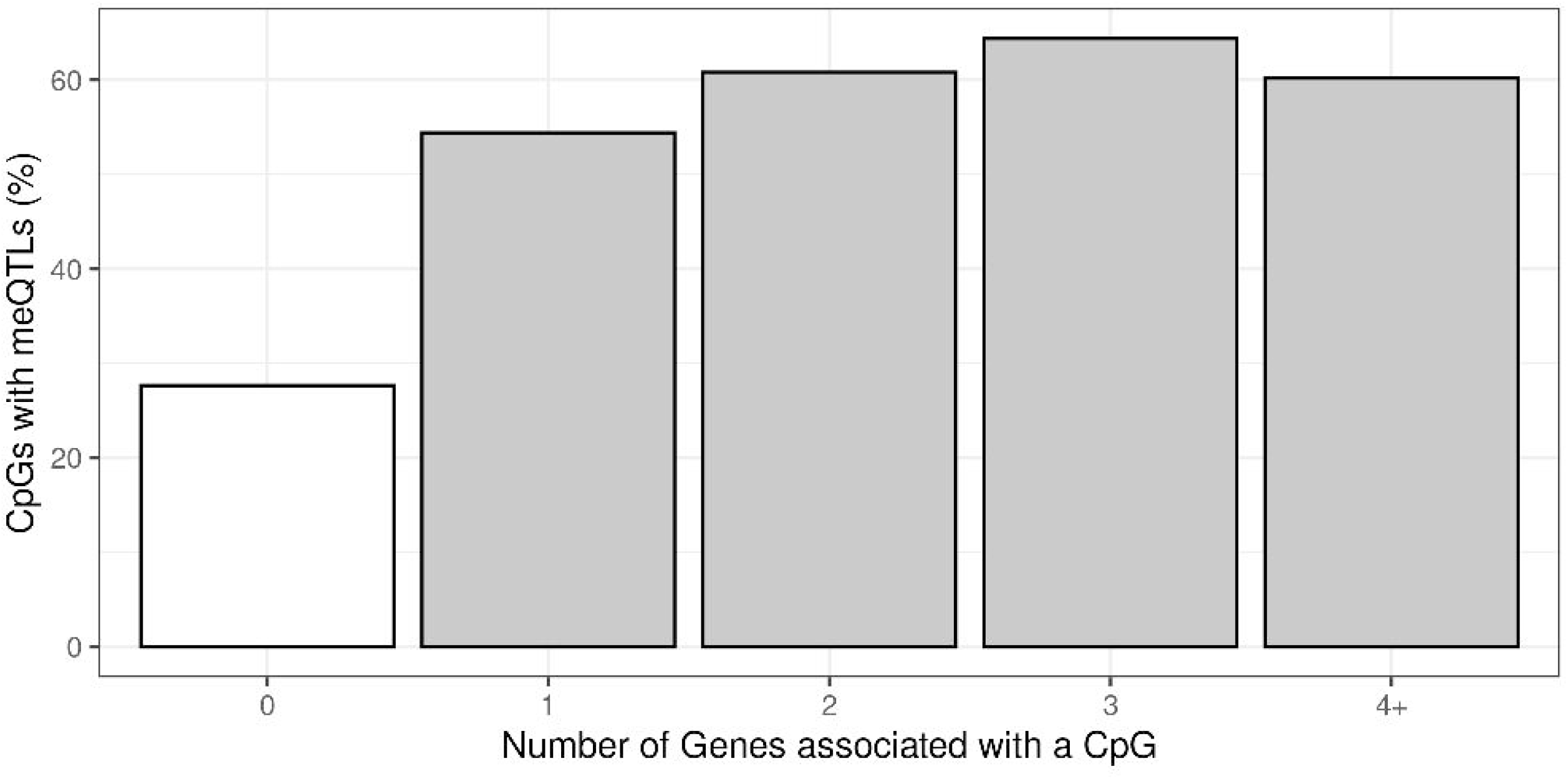
Genetic contribution to autosomal cis eQTMs in children’s blood. CpGs were grouped by the number of Genes they were associated with, where 0 means that a CpG was not associated with any Gene (non eCpG). A) Total additive heritability and B) SNP heritability as inferred by Van Dongen and colleagues (van Dongen et al., 2016). The y-axis represents heritability and the x-axis each group of CpGs associated with a given number of Genes. C) Proportion of CpGs having a meQTL (methylation quantitative trait locus), by each group of CpGs associated with a given number of Genes.

#### Overlap with methylation and expression quantitative trait loci (meQTLs and eQTLs)

Second, we studied whether eCpGs were enriched for meQTLs, either in cis or trans. We analyzed 1,078,466 meQTLs identified in blood samples of 7-year-old children in the ARIES dataset and replicated in HELIX (see Material and Methods). These meQTLs affected the methylation of 36,671 CpGs through a total of 2,820,145 SNP-CpG pairs. 10,187 eCpGs (27.8% total eCpGs) presented at least 1 meQTL, being eCpGs enriched in CpGs associated with genetic variants (OR: 11.06, p-value <2e-6). In addition, among CpGs with meQTLs, eCpGs were associated with a higher number of meQTLs (median: 74, IQR: 27; 162) than non eCpGs (median: 32, IQR = 10; 77). Finally, eCpGs associated with a higher number of eGenes are more likely to be associated with at least one meQTL (Figure 4C). After removing CpG probes with unreliable measurements (ICC <0.4), we observed the same trends, although the enrichment of eCpGs for CpGs with at least one meQTL was reduced (OR = 3.5, p-value <2.2e-16) (Figure 4 – figure supplement 2). Finally, we observed that eCpGs with at least one meQTL were measured with higher reliability (higher ICC) than eCpGs without any meQTL (Figure 4 – figure supplement 3).

We, then, determined whether meQTLs were also eQTLs for the eGenes. After multiple-testing correction, we identified 1,368,613 SNP-CpG-Gene trios with consistent direction of effect, and 12,799 with inconsistent direction. These formers comprised 16,055 unique eQTMs (40.4% of significant eQTMs); 8,503 unique eCpGs (38.7% of total eCpGs); and 4,098 unique eGenes (46.1% of total eGenes), of which 3,154 were coding (50.2% of total coding eGenes). In these trios, eGenes were associated with a median of 2 eCpGs (IQR = 1; 5) and 67 SNPs (IQR = 21; 149); whereas eCpGs were associated with a median of 1 eGene (IQR = 1; 2) and 53 SNPs (IQR = 17; 124). One example of such a SNP-CpG-Gene trio is formed by rs11585123-cg15580684-TC01000080.hg.1 (*AJAP1*), in chromosome 10 (Figure 4 – figure supplement 4).

Next, we run gene-set enrichment analyses with the 2,746 eGenes involved in these trios. We identified 35 significant GO-BP terms (q-value <0.001). Of these, 14 were related to immunity (6 innate, 4 adaptive immunity, and 4 general/other); 11 to cellular processes; and 10 to metabolic processes (Table S1). In comparison to all eGenes, eGenes under genetic control had a reduction in the number of GO-BP terms involving immune and cellular functions (Table S2).

Overall, we found that a substantial part of the eQTMs seems to be under genetic control, and the SNPs associated with DNA methylation levels were also associated with gene expression levels.

### Influence of age on autosomal cis eQTMs in children’s blood

#### Enrichment for age-variable eQTMs

To understand the association between changes in methylation and gene expression throughout life, first we evaluated whether eCpGs were enriched for CpGs with variable blood methylation levels from birth to childhood/adolescence. To this end, we retrieved the CpGs that vary with age from two databases: 14,150 CpGs with variable methylation levels in children between 0 and 8 years (9,647 with increased and 4,503 with decreased methylation) from the MeDALL project (Xu et al., 2017); and 244,283 CpGs with variable methylation levels in children and adolescents between 0 and 17 years (168,314 with increased and 75,969 with decreased methylation) from the Epidelta project (RH et al., 2021). Of note, 90% of the CpGs identified in the MeDALL project were also reported in the Epidelta. We found that eCpGs were enriched for age-variable CpGs in both MeDALL and Epidelta databases, but more markedly for CpGs reported in MeDALL (Figure 5A). In both databases, positive and inverse eCpGs showed stronger ORs for CpGs with increased and decreased methylation levels over age, respectively. After excluding CpG probes with unreliable measurements (ICC <0.4), MeDALL enrichments were reduced to the magnitude of Epidelta enrichments, while the differences between positive and inverse eCpGs were more evident (Figure 5 – figure supplement 1).

**Figure 5.**
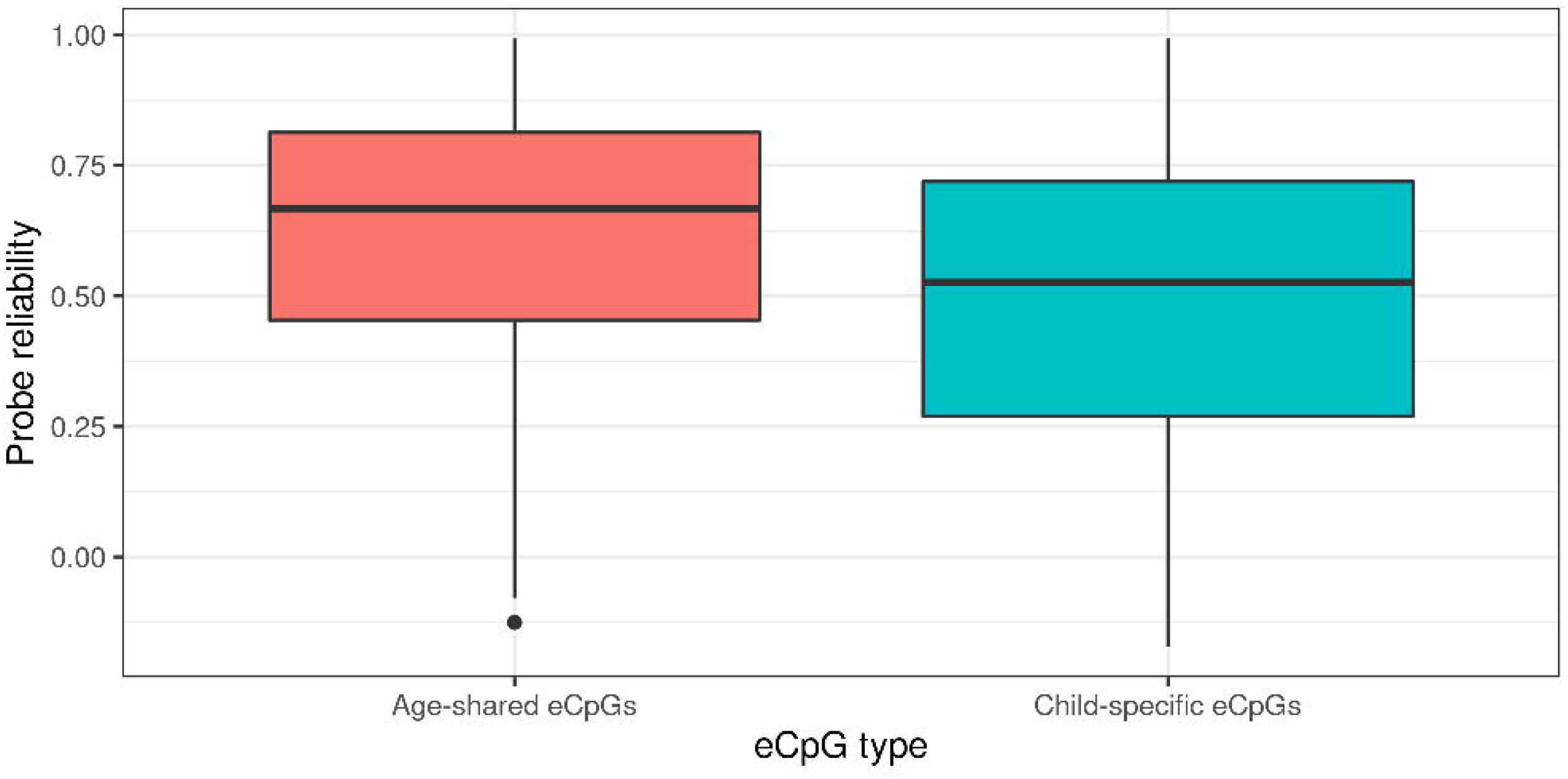
Influence of age on autosomal cis eQTMs in children’s blood. A) Enrichment of eCpGs for CpGs with age-variable methylation levels, in comparison to non eCpGs. eCpGs were classified in all (grey); inverse (yellow); and positive (green). Age-variable CpGs were retrieved from the MeDALL project (from birth to childhood (Xu et al., 2017)) and from the Epidelta project (from birth to adolescence (RH et al., 2021)). They were classified in variable (CpGs with methylation levels that change with age); decreased (CpGs with methylation levels that decrease with age); and increased (CpGs with methylation levels that increase with age). The y-axis represents the odds ratio (OR) of the enrichment. B) Overlap between autosomal cis/trans eQTMs identified in adults (GTP: whole blood; MESA: monocytes) (Kennedy et al., 2018) with cis eQTMs identified in children (HELIX: whole blood). All CpG-gene pairs reported at p-value <1e-5 in GTP or MESA that could be compared with pairs in HELIX are shown. C) Overlap between blood autosomal cis eQTMs identified in HELIX children with cis/trans eQTMs identified in adults (GTP: whole blood; MESA: monocytes) (Kennedy et al., 2018). All CpG-gene pairs in HELIX that could be compared with pairs in GTP or MESA are shown. Note: The comparison has been split into two plots because one eGene in HELIX can be mapped to different expression probes in GTP and MESA, and vice-versa. Only comparable CpG-Gene pairs are shown (see Material and Methods).

#### Overlap with autosomal eQTMs in adult blood

We evaluated whether autosomal cis eQTMs in children’s blood were consistent in adult populations. For this, we used data from the study of autosomal cis and trans eQTMs in adults’ blood based on two cohorts: (1) GTP, whole blood and 333 samples; and (2) MESA, monocytes and 1,202 samples, by Kennedy and colleagues (Kennedy et al., 2018). The catalogue contains the summary statistics of all autosomal cis (<50 kb from the TSS) and trans (otherwise) CpG-gene pairs at p-value <1e-5, although only CpG-gene associations at p-value <1e-11 were considered significant eQTMs in their study. To compare their findings with ours, we mapped Genes and gene probes to Gene Symbols and compared CpG-gene pairs (see Material and Methods, Table S3).

We observed that 57.9% and 35.3% of eQTMs with p-value <1e-5 in GTP and MESA were also eQTMs in HELIX, thus age-shared eQTMs (Figure 6B). More than 90% of age-shared eQTMs have the same direction in GTP/MESA than in HELIX (Table S4). In addition, effect sizes in GTP/MESA were correlated with effects sizes in HELIX (Table S4).

Only 5,471 (13.8%) of the eQTMs identified in HELIX children were reported in adult GTP or MESA catalogues at p-value <1e-5 (Figure 6C). We explored whether eQTMs identified both in HELIX children and in adults (age-shared eQTMs) had different characteristics compared to eQTMs only found in children (child-specific eQTMs). Age-shared eQTMs involved 4,364 eCpGs and 1,689 eGenes, whereas children-specific eQTMs involved 19,584 eCpGs and 8,429 eGenes. Age-shared eCpGs had higher reliability (higher ICC) (Figure 5 – figure supplement 2) and tended to be closer to the TSS than child-specific eCpGs (Figure 5 – figure supplement 3). The enrichment for ROADMAP blood chromatin states (Roadmap Epigenomics Consortium et al., 2015) of age-shared and child-specific eCpGs in comparison to non eCpGs was quite similar (Figure 5 – figure supplement 4). Nonetheless, age-shared eCpGs showed higher ORs of enrichment for proximal promoters. Both types of eCpGs were enriched for meQTLs compared to non eCpGs, with the OR being stronger for age-shared eCpGs (OR = 20.7) than for child-specific eCpGs (OR = 10.3).

Overall, we found that eQTMs were enriched for CpGs whose methylation levels changed from birth to adolescence. The overlap between child and adult eQTMs was small: only 13.8% of HELIX eQTMs had also been described in adults. Age-shared eCpGs tended to be proximal to the TSS, enriched for promoter chromatin states, and with stronger signals of genetic regulation.

## Discussion

In this work, we present a blood autosomal cis eQTM catalogue in children. We identified 39,749 eQTMs, representing 21,966 unique eCpGs and 8,886 unique eGenes (6,288 of which were coding). 23,355 eQTMs (58.8% of all eQTMs) showed inverse associations. A substantial fraction was influenced by genetic variation, and the overlap with eQTMs reported in adults was small.

The characteristics of the autosomal cis eQTMs in children’s blood were highly consistent with patterns previously described in other studies. Most of the eCpGs tended to be proximal to the eGene’s TSS (Kennedy et al., 2018; Leland Taylor et al., 2019). The magnitude of the effect seemed to be proportional to the distance between the eCpG and the eGene’s TSS, but this association was weak. Although higher DNA methylation is assumed to lead to lower expression, we found that around 40% of eQTMs were positively associated with gene expression. This percentage is in line with previous results from different tissues (Gutierrez-Arcelus et al., 2015, 2013; Küpers et al., 2019). Inverse and positive eCpGs tended to be localized in enhancers and other active regulatory regions and not in CpG islands, a pattern that was also previously reported (Gutierrez-Arcelus et al., 2015; Küpers et al., 2019). Despite these common locations, inverse eCpGs were specifically found around active TSSs (including the distal promoter and the 5’UTR), while positive eCpGs were localized in gene body regions. These results highlight the importance of the genomic context to infer the direction of the association between DNA methylation and gene expression (Kennedy et al., 2018). We want to point out that the causal relationship between DNA methylation and gene expression cannot be definitely inferred from our study. Indeed, there is some evidence suggesting that DNA methylation could be a consequence of gene expression, as opposed to the often assumed concept that regulation of gene expression is mediated by DNA methylation (Gutierrez-Arcelus et al., 2013; Jones, 2012).

eQTMs can be influenced by genetic variation (Lu et al., 2019). In HELIX, eCpGs linked to the expression of several eGenes had higher heritabilities and were associated with a higher number of meQTLs than non eCpGs. This could suggest that eCpGs that regulate the expression of several genes, the so-called master regulators, are more prone to be themselves regulated by genetic variation. We, then, searched for SNPs simultaneously associated with DNA methylation (meQTLs) and gene expression (eQTLs) in our data. We identified 1.3 M SNP-CpG-Gene trios with consistent direction of the effect. Interestingly, the number of GO-BP terms related to immune and cellular functions was reduced for eGenes under genetic control, in comparison to all eGenes; on the contrary, the number of GO-BP terms involving metabolic processes was maintained. This may suggest that the influence of environmental factors is more relevant for immune pathways, while genetic factors might be more determinant in regulating metabolic processes in blood cells. Given the non-negligible effect of genetics in eQTMs, we would advise studying the effect of genetic variants on the association between environmental factors or phenotypic traits and DNA methylation.

In order to know how eQTMs behave along life-course, we compared blood autosomal cis eQTMs identified in HELIX children with cis and trans eQTMs reported by Kennedy and colleagues in whole blood and monocytes from adult populations (Kennedy et al., 2018). We found that only 13.8% of the autosomal eQTMs in children’s blood were also reported in adults. Similarly, a modest proportion of adult blood eQTMs was present in children (58% from GTP and 35% from MESA). This small overlap between adult and child eQTMs has different explanations: methodological issues, such as gene expression platforms with low overlap; statistical methods and statistical power; cohort-specific environmental exposures; and cellular composition. Unsurprisingly, HELIX and MESA presented the highest divergence, as HELIX assessed eQTMs in whole blood and MESA in monocytes. Despite the effect of these methodological and confounding factors, it is known that DNA methylation and gene expression change with age (Melé et al., 2015; RH et al., 2021; Xu et al., 2017); consequently, we could expect only partial overlap between adult and child eQTMs. The short list of age-shared eCpGs tended to encompass CpGs located in promoters and regulated by genetic variants. Moreover, the overall location of eQTMs in regulatory elements was similar between adults and children (Gutierrez-Arcelus et al., 2015; Küpers et al., 2019). This could represent a specific characteristic of eQTMs that are persistent over time. An alternative explanation is that this kind of eQTMs (genetically regulated and close to the TSS) are easier to be detected and shared among any two studies because they show stronger effects. Finally, we observed that HELIX eQTMs usually involved CpGs whose methylation varied between birth and childhood/adolescence, and they tended to activate rather than inactivate transcription over this period. Also, they were enriched for CpGs found to be related to environmental factors and phenotypic traits in the EWAS Atlas and EWAS Catalog.

As previously described (Sugden et al., 2020), CpG probes have different measurement error and thus different reliability and reproducibility. Consequently, CpGs measured with less error have more chances of being found associated with traits and thus reported in EWAS catalogues. In HELIX, we found that CpG probe ICC was higher for these different cases: for eCpGs, in comparison to non eCpGs; for age-shared eCpGs, in comparison to children-specific eCpGs; and for eCpGs with meQTLs, in comparison to eCpGs without meQTLs. In this line, enrichments of eCpGs for CpGs listed in the EWAS Atlas or in the MeDALL project were markedly attenuated when only considering CpGs measured with good reliability. Moreover, CpG probe reliability is dependent on DNA methylation level and variance (highly unmethylated or highly methylated CpGs, which tend to have low variances, are measured with more error); and genomic regulatory elements are characterized by particular methylation levels. Therefore, this biased the enrichments for regulatory elements. For instance, after considering only reliable probes, the distribution of eQTMs in CpG island relative positions changed completely (Figure 3 – figure supplement 1). Moreover, the enrichments for active chromatin states were amplified and differences between inverse and positive eCpGs attenuated.

Our study of autosomal cis eQTMs in children’s blood has several strengths compared to previous eQTM studies. First, we report all CpG-gene pairs we tested in our analysis, as opposed to existing blood eQTM catalogues which only reported pairs passing a given p-value threshold (Bonder et al., 2017; Kennedy et al., 2018). Reporting all pairs tested allows replication and meta-analyses, reducing publication bias. Second, we report which eQTMs are influenced by genetic variation, and researchers can take this into account when exploring the relationship between methylation and expression in their data. Finally, as others (Wu et al., 2018), we describe that only around half of the CpG-Gene relationships are captured through annotation to the closest gene. Therefore, our eQTM catalogue becomes an essential and powerful tool to help researchers interpret their EWAS, with a particular focus on childhood.

The catalogue also has some limitations. First, it only covers a fraction of all CpG-Gene pairs, as both the methylation and gene expression arrays have limited resolution. Nonetheless, the catalogue will be useful for most researchers as the methylation array is widely used, and the gene expression array covers almost all the coding genes. Second, the catalogue does not include sex chromosomes which require more complex analyses to address X-inactivation and sex-specific effects that will be addressed in future studies. Third, due to statistical power limitations, only cis effects were tested. Despite that, we observed that eCpGs tended to be close to the gene they regulate, so the catalogue is expected to cover most of the CpG-Gene associations. Fourth, effect sizes should be considered with caution as the association between DNA methylation and gene expression might be non-linear, and the effect of outlier values was not systematically explored (Johnson et al., 2017). Fifth, models were adjusted for blood cell type composition and, while this has allowed us to control for major differences in methylation and gene expression among blood cell types, it might also have resulted in over-adjustment in some CpG-Gene pairs. Moreover, the analysis of bulk data might have limited the identification of eQTMs specific to a subset of blood cell types, the identification of which would need more sophisticated statistical and/or experimental methods. Finally, we acknowledge that the catalogue will be useful for biological interpretation of EWAS if it is true that DNA methylation is not a mere mark of cell memory to past exposures (without transcriptional consequences or with time-limited ones) (Tsai et al., 2018).

In summary, besides characterizing child blood autosomal cis eQTMs and reporting how they are affected by genetics and age, we provide a unique public resource: a catalogue with 13.6 M CpG-gene pairs and of 1.3 M SNP-CpG-gene trios (https://helixomics.isglobal.org/).

This information will improve the biological interpretation of EWAS findings.

## Figure legends

**Figure 1 – figure supplement 1.**
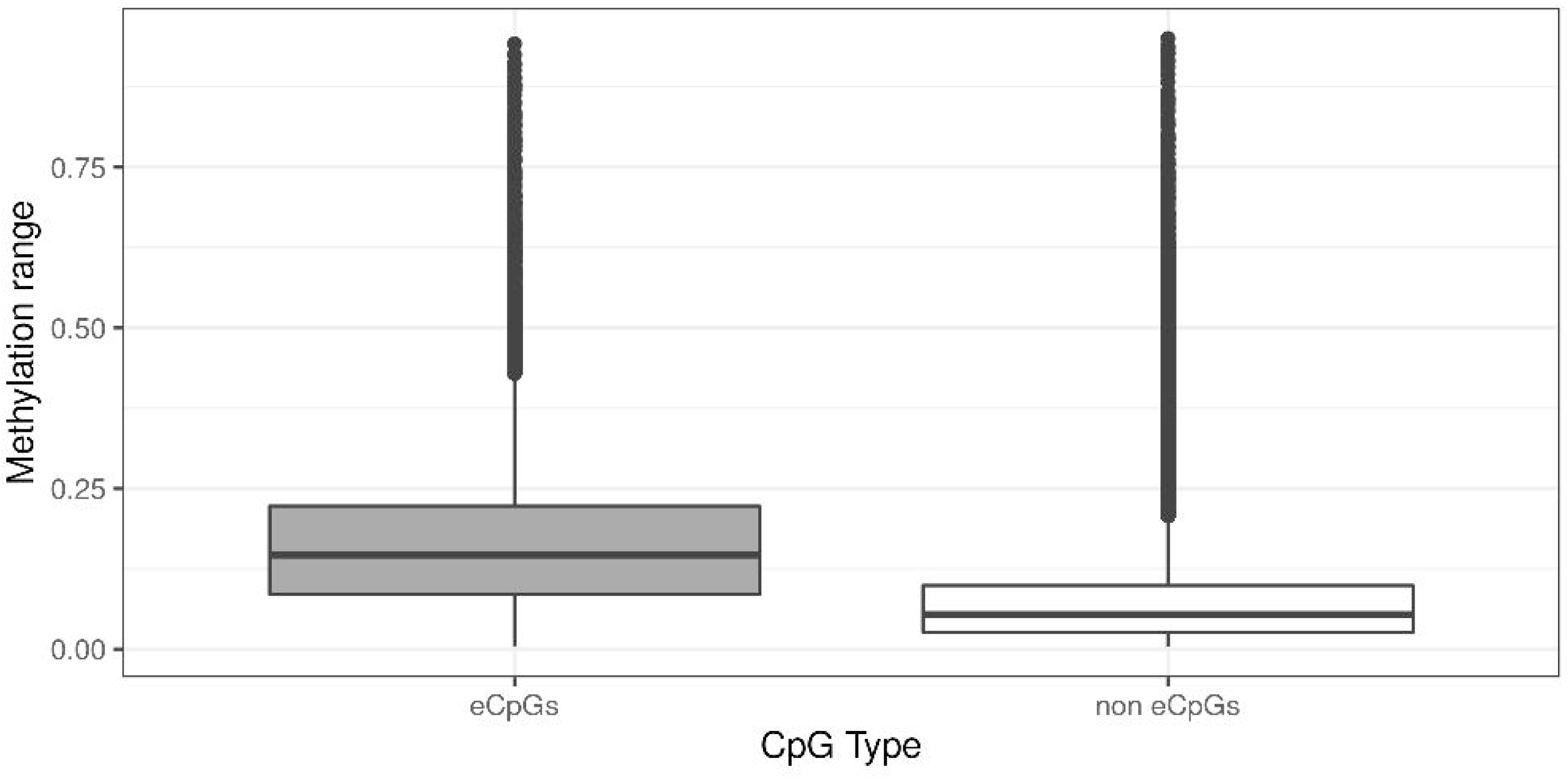
Distribution of Genes and CpGs in all CpG-Gene pairs. A) Distribution of the number of Genes paired with each CpG. The y-axis represents the number of CpGs that are paired with a given number of Genes, indicated in the x-axis. The vertical line marks the median of the distribution. Each CpG was paired to a median of 30 Genes (IQR: 20; 46). B) Distribution of the number of CpGs paired with each Gene. The y-axis represents the number of Genes that are paired with a given number of CpGs, indicated in the x-axis. The vertical line marks the median of the distribution. Each Gene was paired to a median of 162 CpGs (IQR: 93; 297).

**Figure 1 – figure supplement 2.**
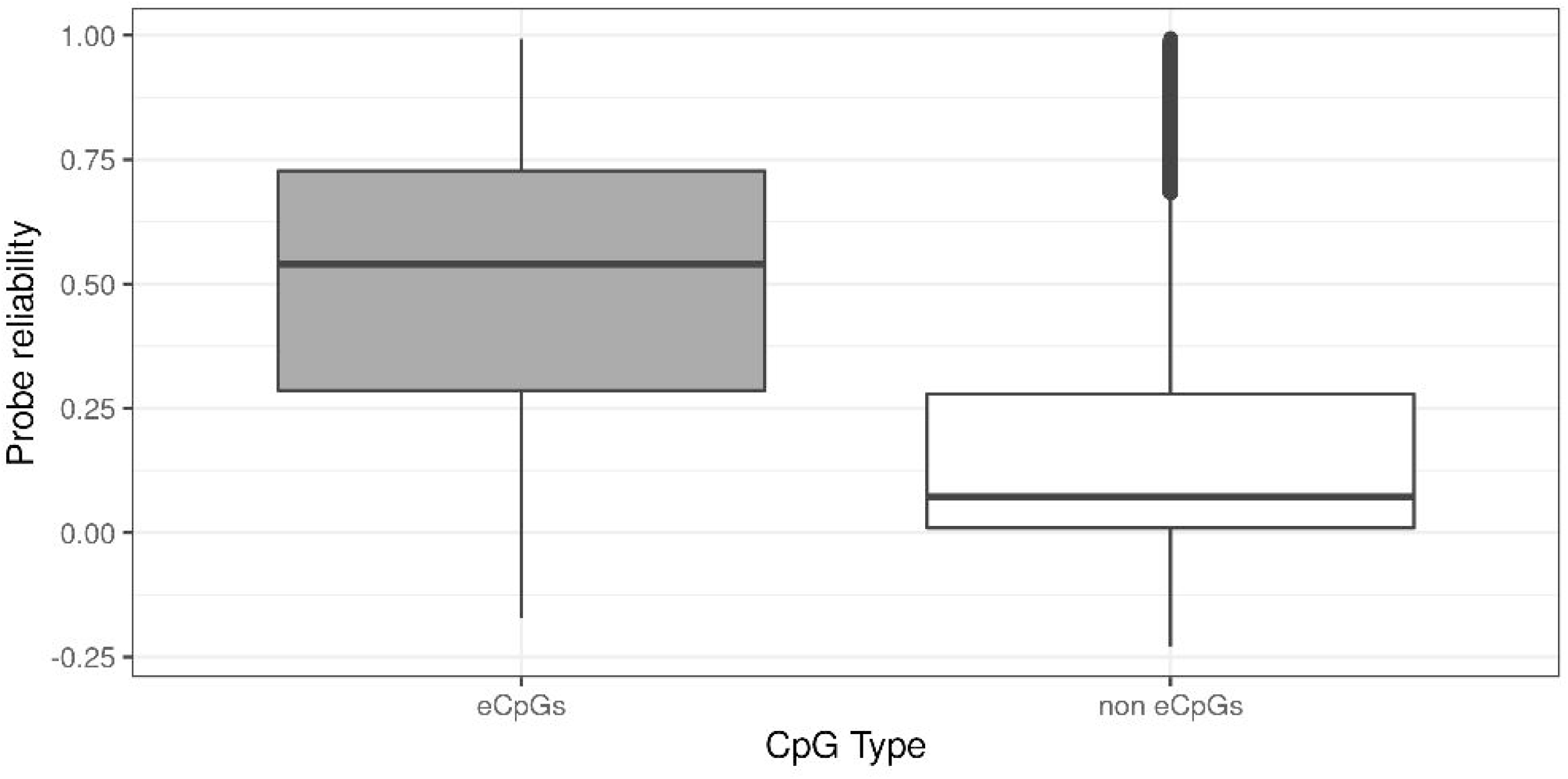
Distribution of eGenes and eCpGs in autosomal cis eQTMs. A) Distribution of the number of eGenes paired with each eCpG in eQTMs. The y-axis represents the number of eCpGs that are paired with a given number of eGenes, indicated in the x-axis. Each eCpG was associated with a median of 1 eGene (IQR = 1; 2). B) Distribution of the number of eCpGs paired with each eGenes in eQTMs. The y-axis represents the number of eGenes that are paired with a given number of eCpGs, indicated in the x-axis. Each eGene was associated with a median of 2 eCpGs (IQR = 1; 5).

**Figure 1 – figure supplement 3.**
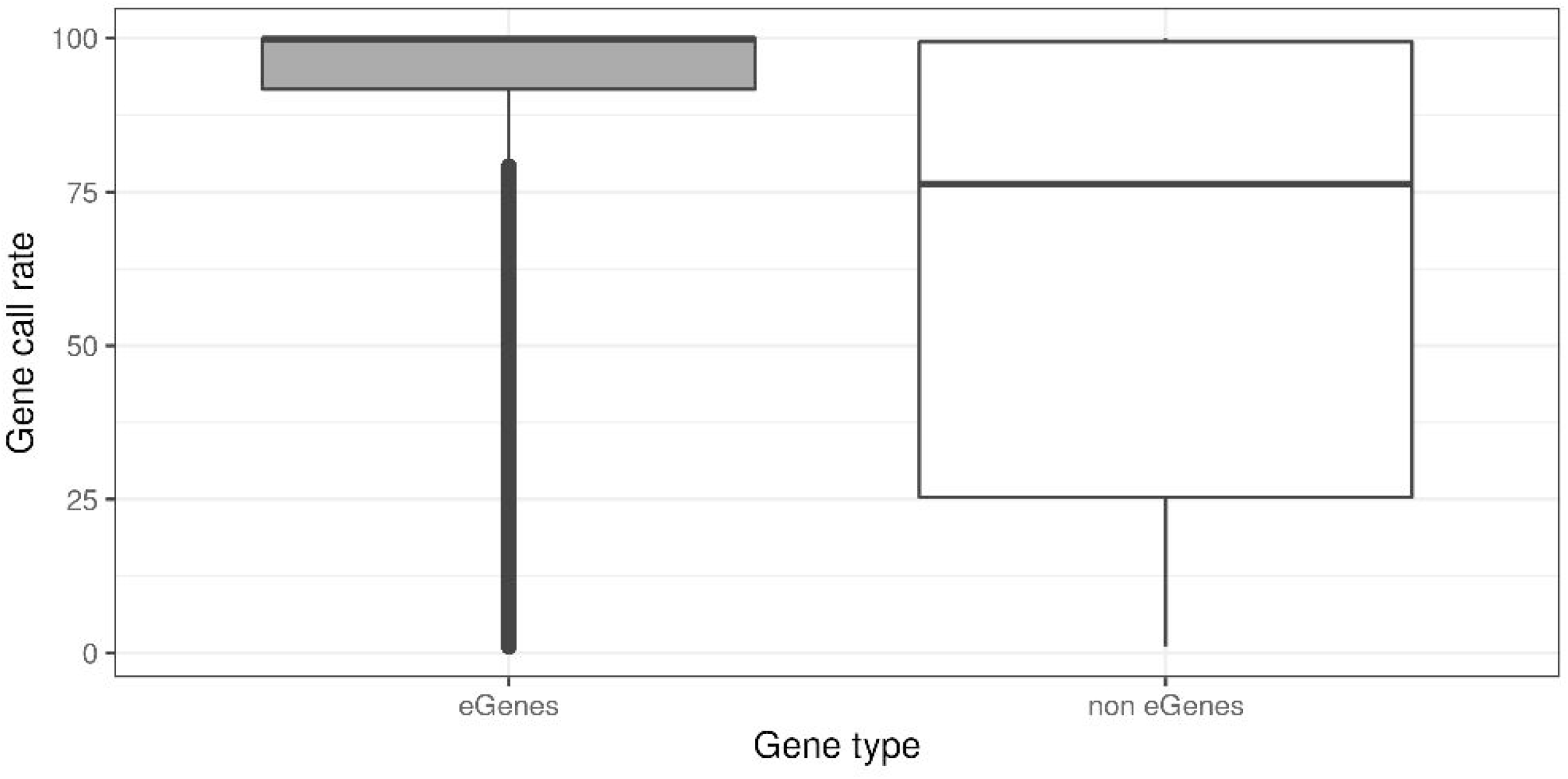
DNA methylation range by CpG type. CpGs were classified in: eCpGs (CpGs associated with gene expression, N=21,966, in grey) and non eCpGs (N=364,452, in white). Methylation range was computed as the difference between the methylation values in percentile 1 and percentile 99 (Lin et al., 2017).

**Figure 1 – figure supplement 4.**
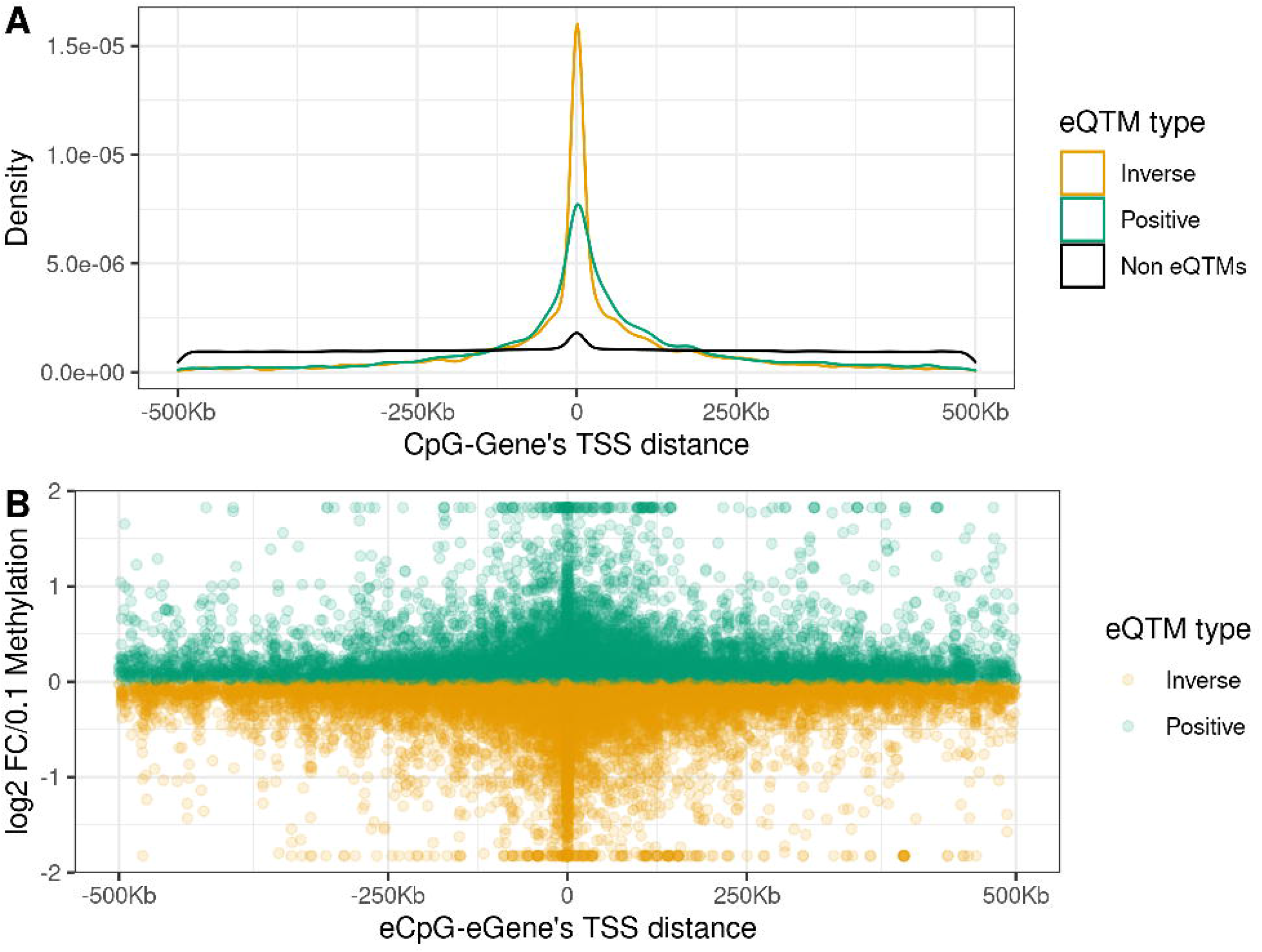
Probe reliability by CpG type. CpGs were classified in: eCpGs (CpGs associated with gene expression, N=21,966, in grey) and non eCpGs (N=364,452, in white). Probe reliability was based on intraclass correlation coefficients (ICC) obtained from (Sugden et al., 2020).

**Figure 1 – figure supplement 5.**
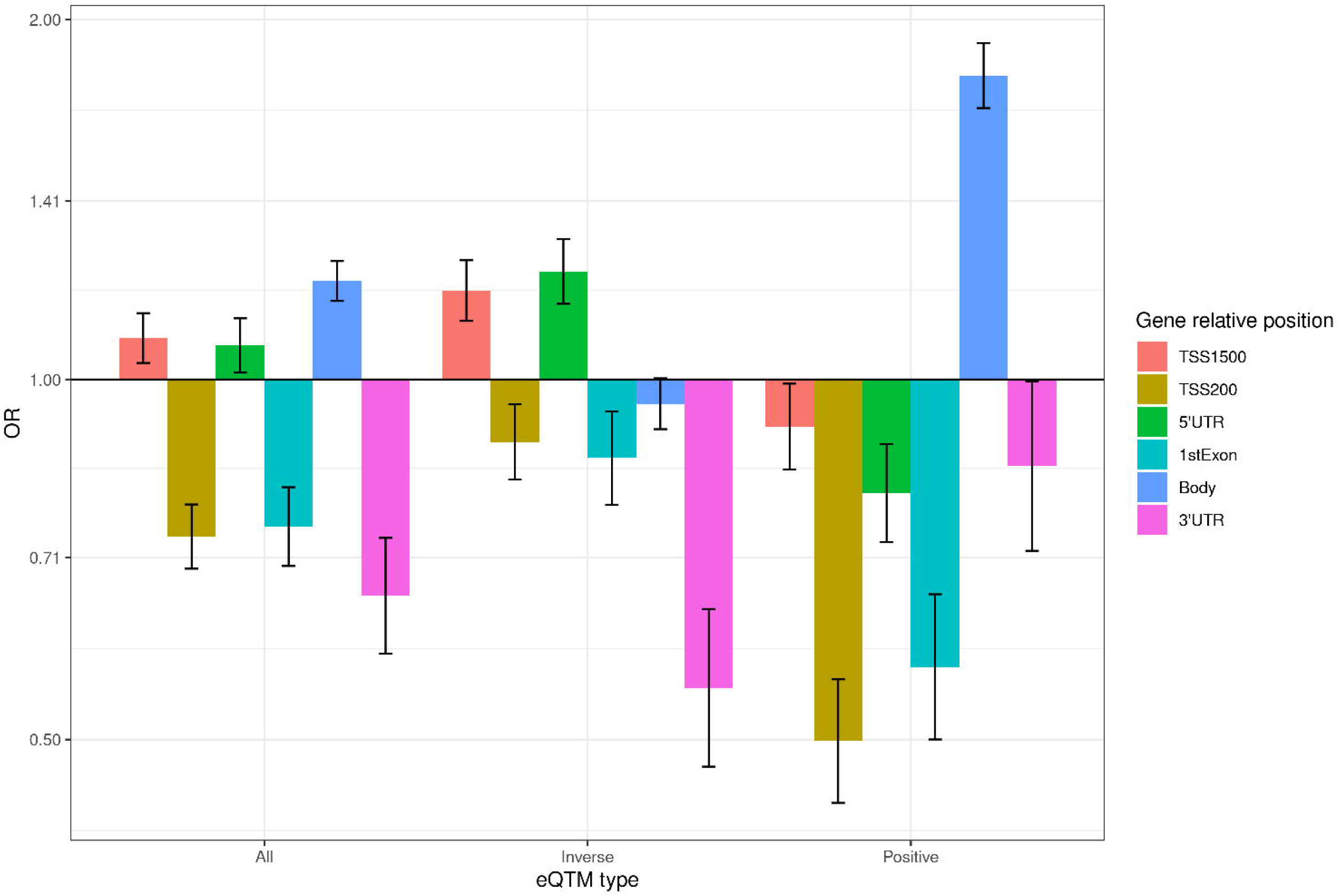
Genes call rate distribution by Gene type. Genes were classified in: eGenes (Genes associated DNA methylation, N=8,886, in grey) and non eGenes (N=51,806, in white). For a given Gene, call rate is the proportion of children with gene expression levels over the background noise.

**Figure 2 – figure supplement 1.**
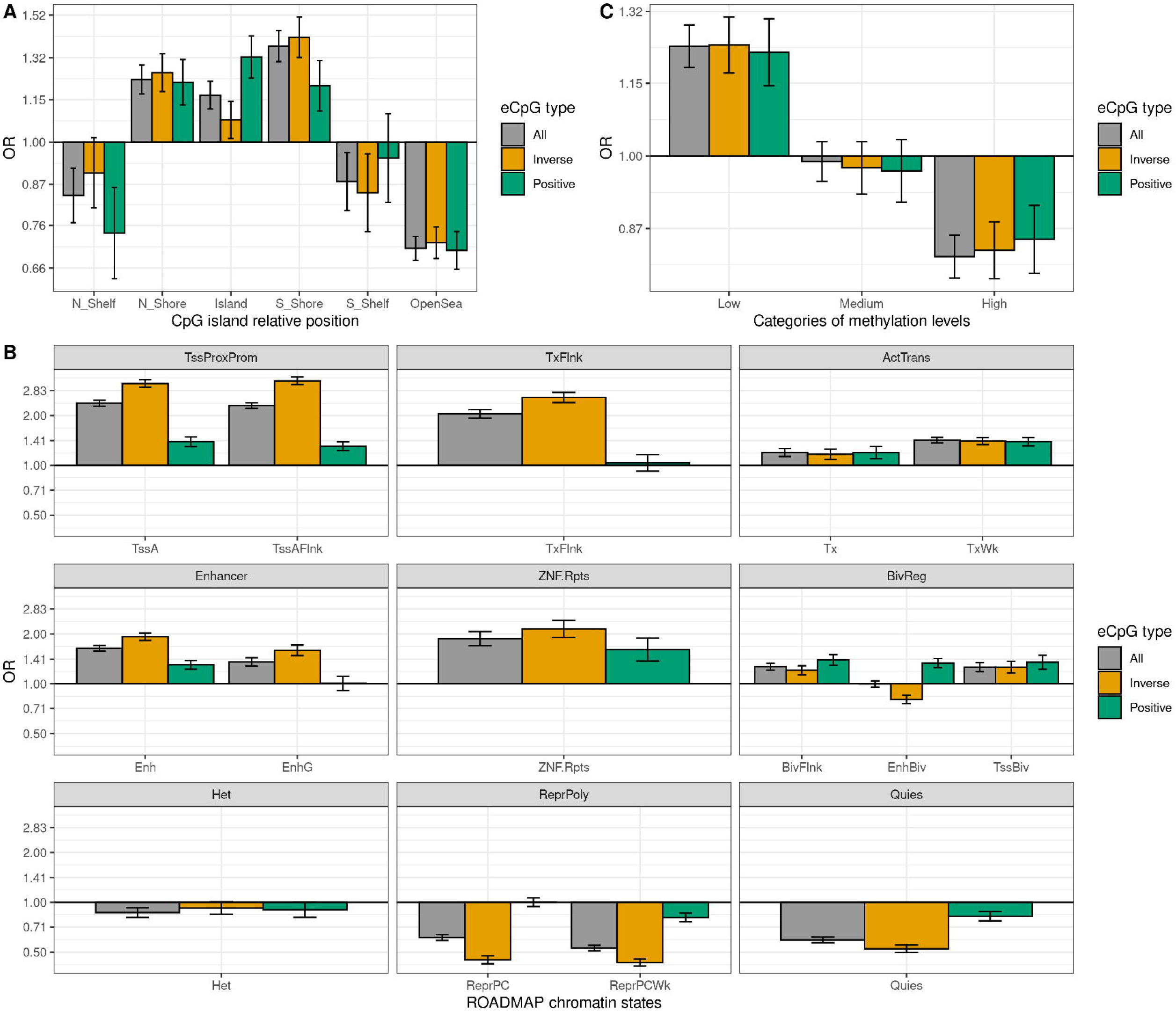
Enrichment of eCpGs for gene relative positions. We selected the subset of 327,931 CpG-Gene pairs where the CpG and the Gene were annotated to the same gene. Enrichment was computed for all eCpGs in this subset, and for inverse and positive eCpGs. Genic regions are classified in distal promoter from 200 to 1,500 bp (TSS1500); proximal promoter up to 200 bp (TSS200), 5’ untranslated region (5’UTR); 1st exon; gene body; and 3’ untranslated region (3’UTR). The y-axis represents the odds ratio (OR) of the enrichment. For all gene relative positions, the enrichment was computed against CpG-Gene pairs with CpG and Gene annotated to the same gene that were not eQTMs.

**Figure 3 – figure supplement 1.**
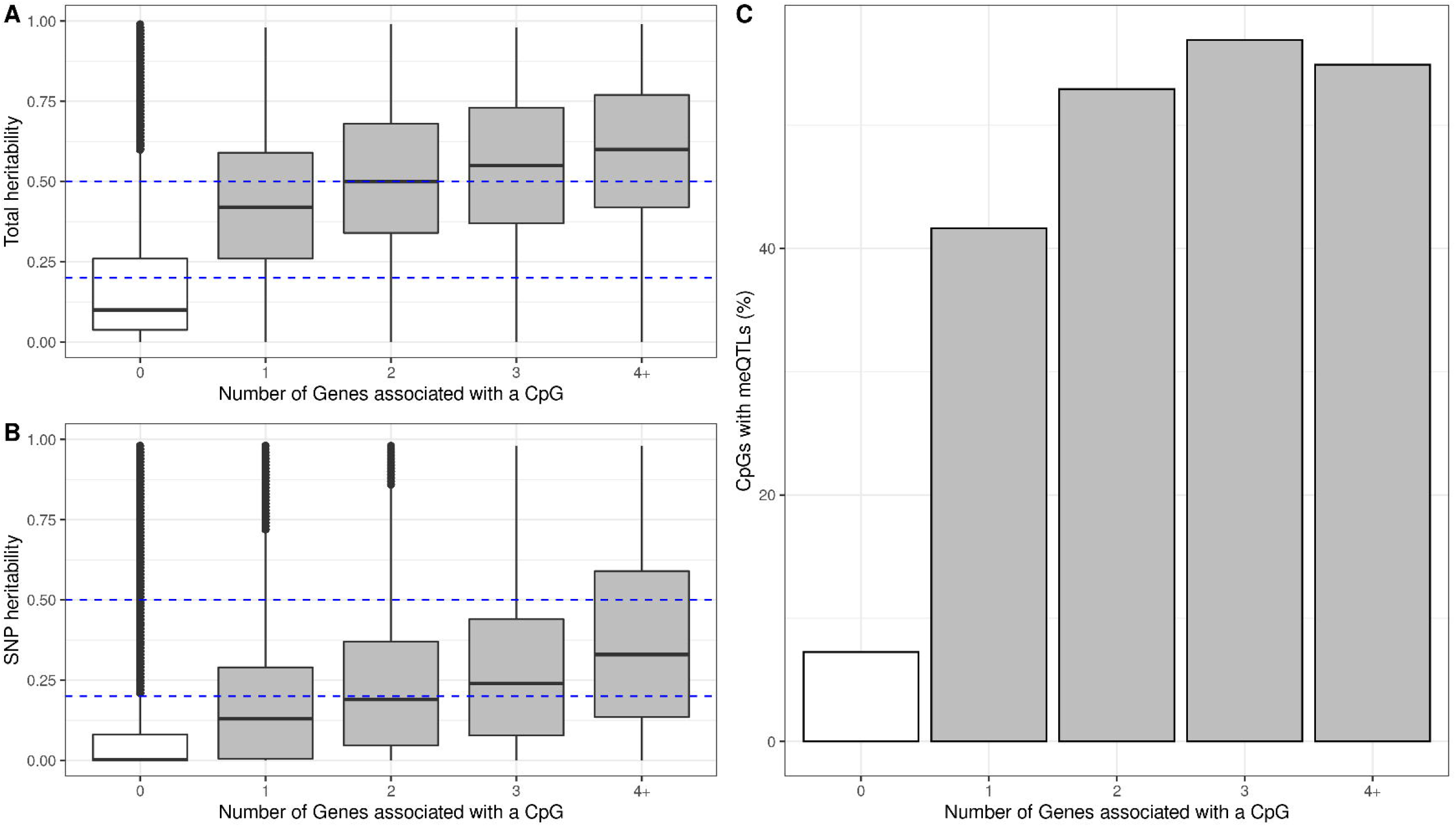
Enrichment of eCpGs with reliable measurement for different regulatory elements. Only eCpGs with reliable measurements (ICC >0.4) were considered (Sugden et al., 2020). eCpGs were classified in all (grey), inverse (yellow), and positive (green). The y-axis represents the odds ratio (OR) of the enrichment. In all cases, the enrichment was computed against non eCpGs. A) Enrichment for CpG island relative positions: CpG island, N- and S-shore, N- and S-shelf, and open sea. B) Enrichment for ROADMAP blood chromatin states (Roadmap Epigenomics Consortium et al., 2015): active TSS (TssA), flanking active TSS (TssAFlnk), transcription at 5’ and 3’ (TxFlnk), transcription region (Tx), weak transcription region (TxWk), enhancer (Enh); genic enhancer (EnhG), zinc finger genes and repeats (ZNF.Rpts), flanking bivalent region (BivFlnx), bivalent enhancer (EnhBiv), bivalent TSS (TssBiv), heterochromatin (Het), repressed Polycomb (ReprPC), weak repressed Polycomb (ReprPCWk), and quiescent region (Quies). Chromatin states can be grouped in active transcription start site proximal promoter states (TssProxProm), active transcribed states (ActTrans), enhancers (Enhancers), bivalent regulatory states (BivReg) and repressed Polycomb states (ReprPoly). C) Enrichment for groups of CpGs with different median methylation levels: low (0-0.3), medium (0.3-0.7), and high (0.7-1) (Huse et al., 2015).

**Figure 3 – figure supplement 2.**
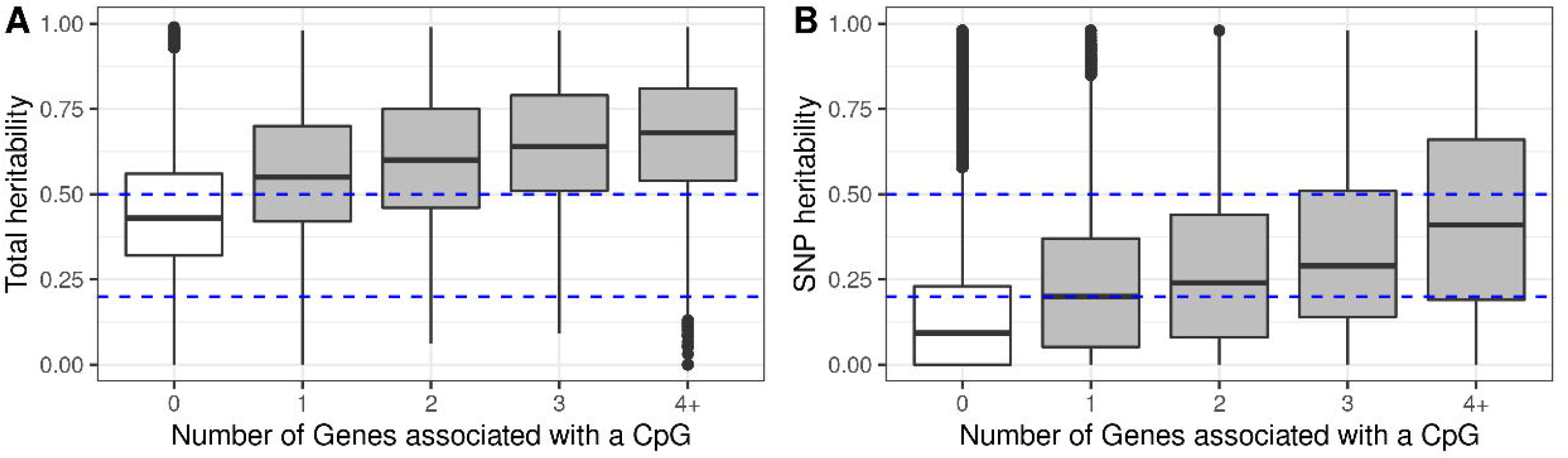
Enrichment of autosomal cis eCpGs in children’s blood for CpGs reported to be associated with phenotypic traits and/or environmental exposures. Enrichment for CpGs present in EWAS datasets: the EWAS Atlas (Li et al., 2019), and the EWAS Catalog (Battram et al., 2021). eCpGs were classified in all (grey), inverse (yellow), and positive (green). In all cases, the enrichment was computed against non eCpGs. The y-axis represents the odds ratio (OR) of the enrichment. A) Enrichment considering all CpGs. B) Enrichment considering only CpGs measured with reliable probes (ICC >0.4) (Sugden et al., 2020). intraclass correlation coefficient.

**Figure 4 – figure supplement 1.**
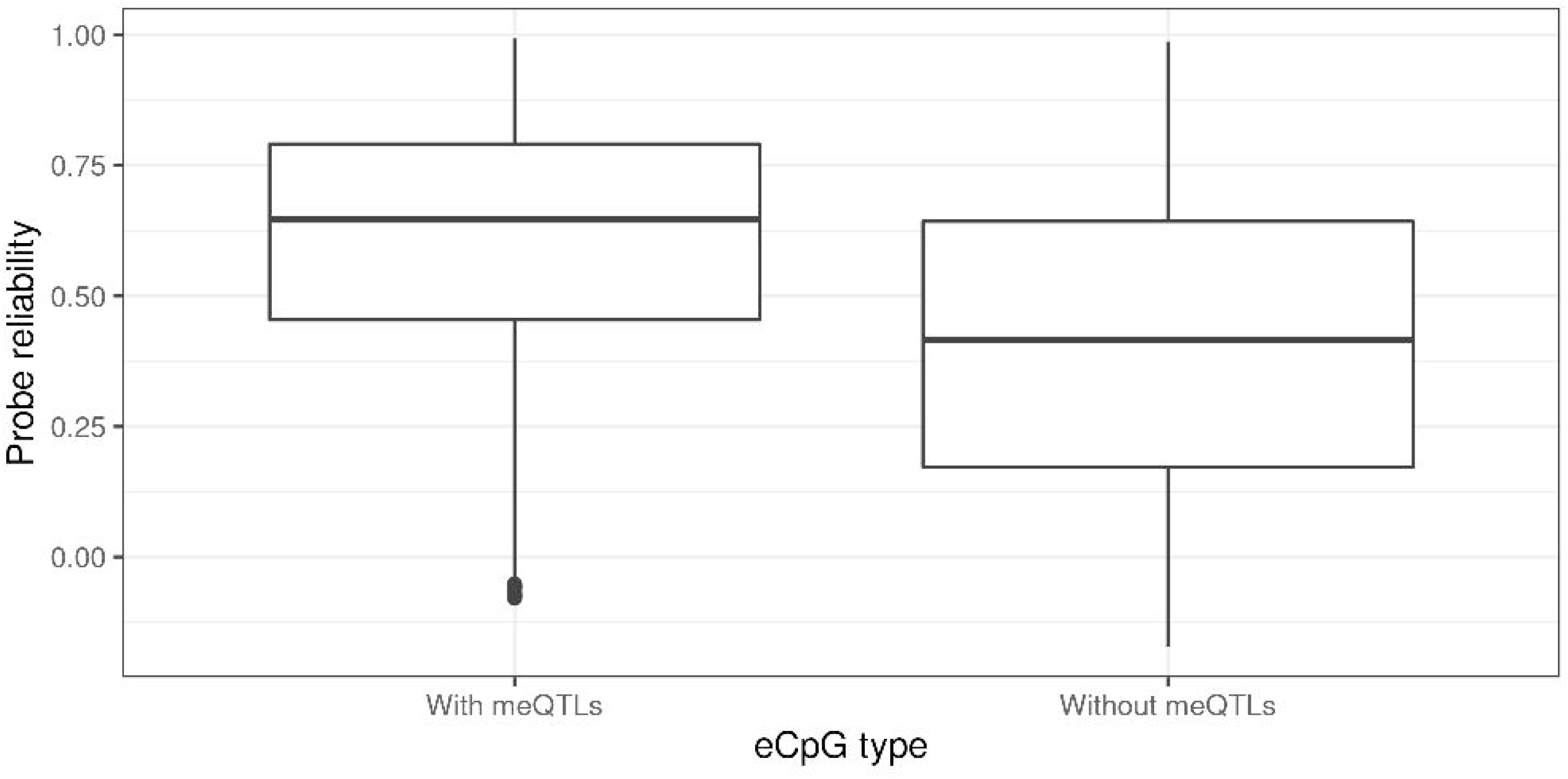
Heritability of methylation levels in CpGs with reliable measurements. Only CpGs measured with reliable probes (ICC >0.4) were considered (Sugden et al., 2020). CpGs were grouped by the number of Genes they were associated with, where 0 means that a CpG was not associated with any Gene (non eCpGs, in white). A) Total additive heritability as inferred by Van Dongen and colleagues (van Dongen et al., 2016), by each group of CpGs associated with a given number of Genes. B) SNP heritability as inferred by Van Dongen and colleagues (van Dongen et al., 2016), by each group of CpGs associated with a given number of Genes.

**Figure 4 – figure supplement 2.**
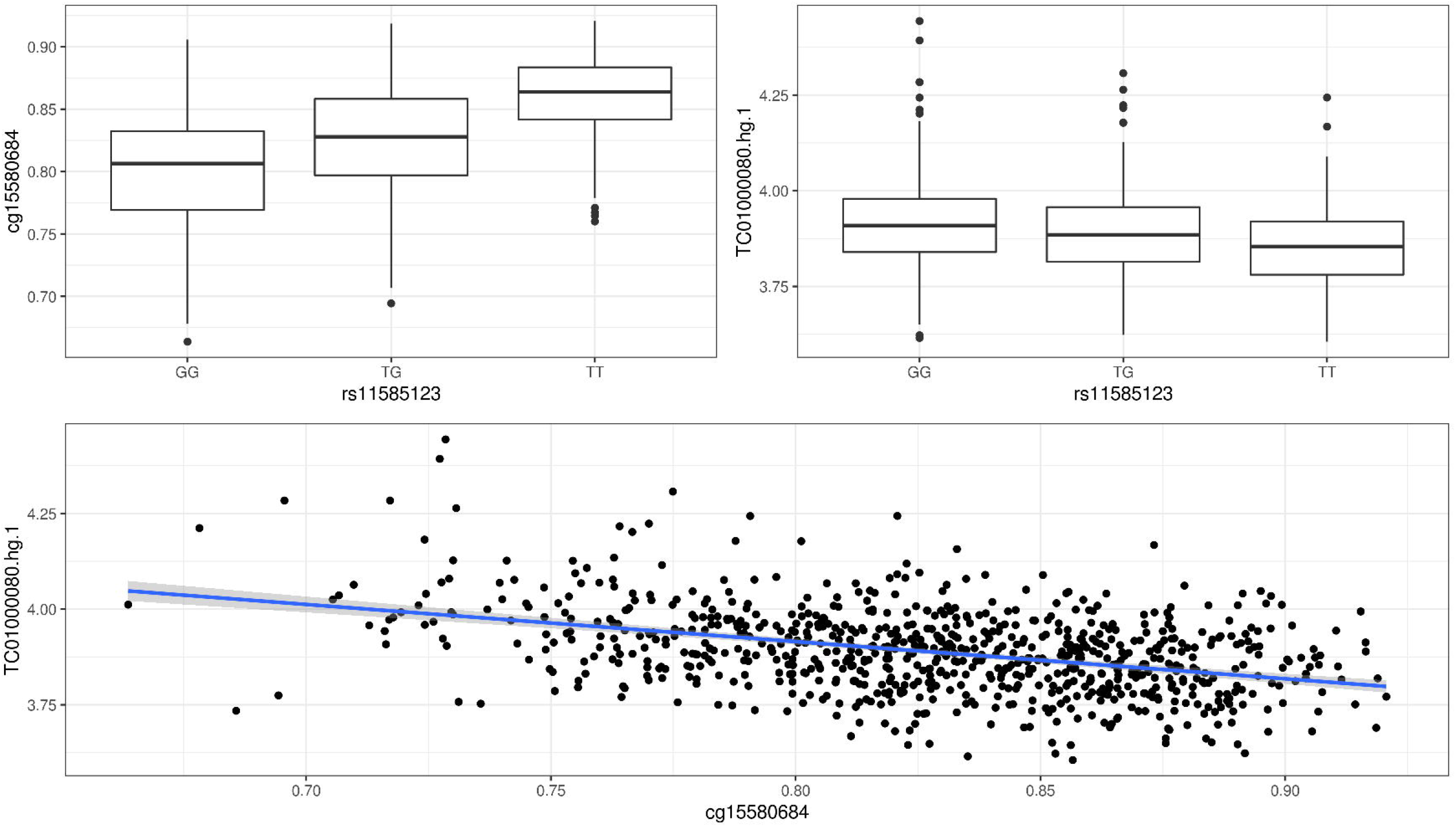
Proportion of CpGs having a meQTL (methylation quantitative trait loci) among CpGs with reliable measurements. Only CpGs measured with reliable probes (ICC >0.4) were considered (Sugden et al., 2020). CpGs were grouped by the number of Genes they were associated with.

**Figure 4 – figure supplement 3.**
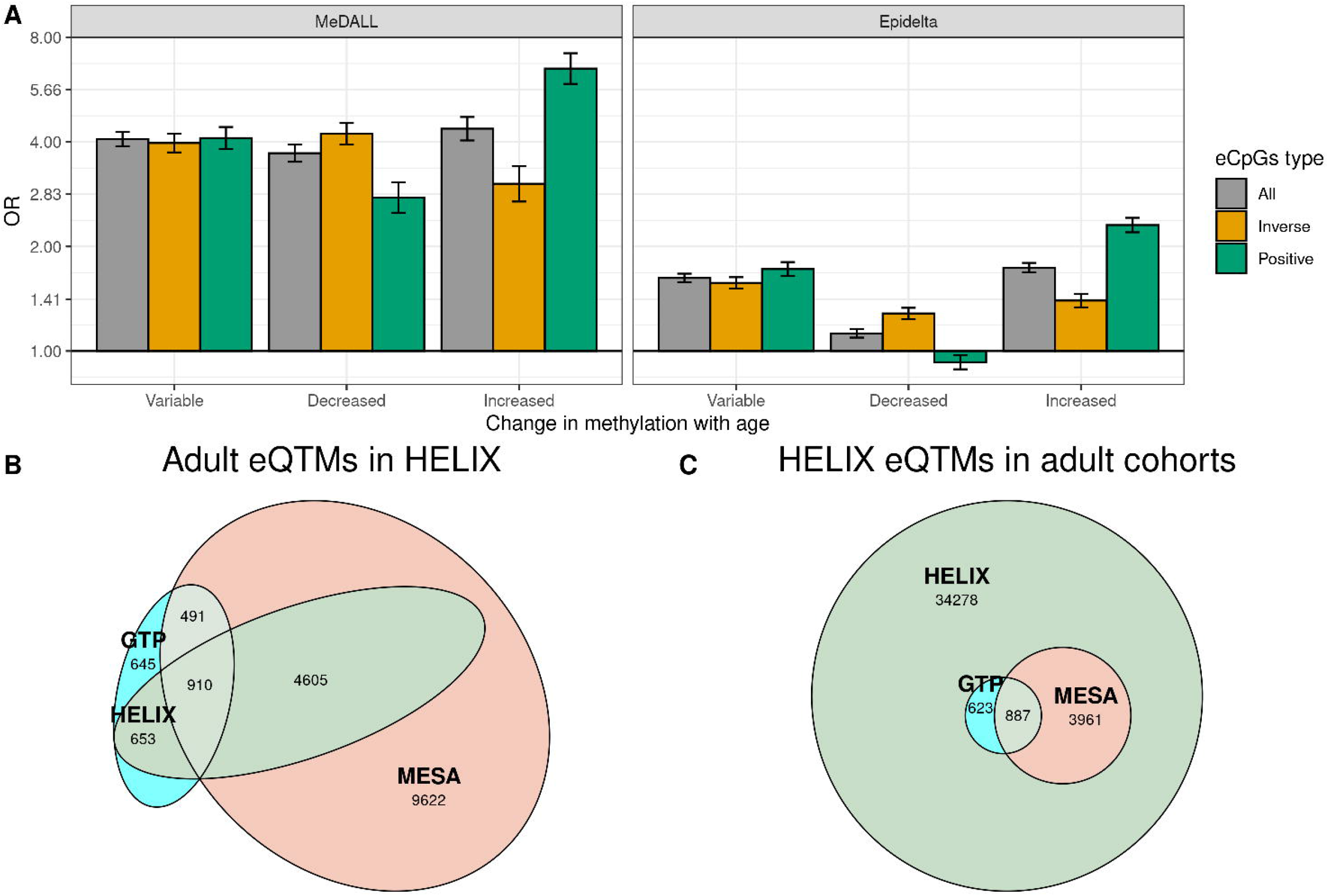
Probe reliability in autosomal cis eCpGs according to association with genetic variants. eCpGs were classified in two groups, depending on whether their methylation values were associated with any genetic variant. Probe reliabilities were based on intraclass correlations (ICCs) obtained from (Sugden et al., 2020).

**Figure 4 – figure supplement 4.**
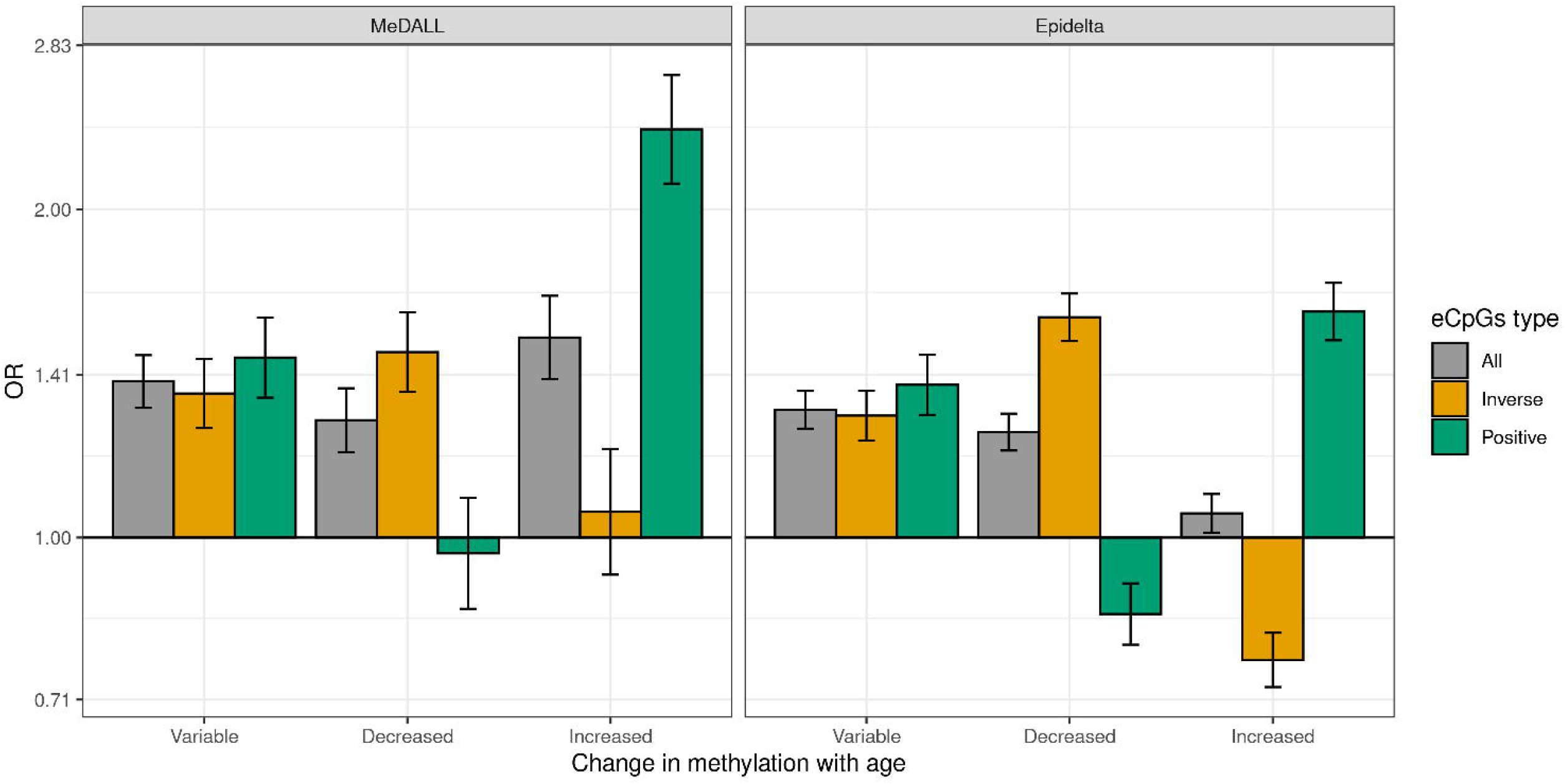
**Example of a trio of SNP-CpG-Gene**. A) Methylation levels (cg15580684) by SNP genotypes (rs11585123). B) Gene expression levels (TC01000080.hg.1, AJAP1 gene) by SNP genotypes (rs11585123). C) Correlation between gene expression (TC01000080.hg.1, AJAP1 gene) and methylation levels (cg15580684).

**Figure 5 – figure supplement 1.**
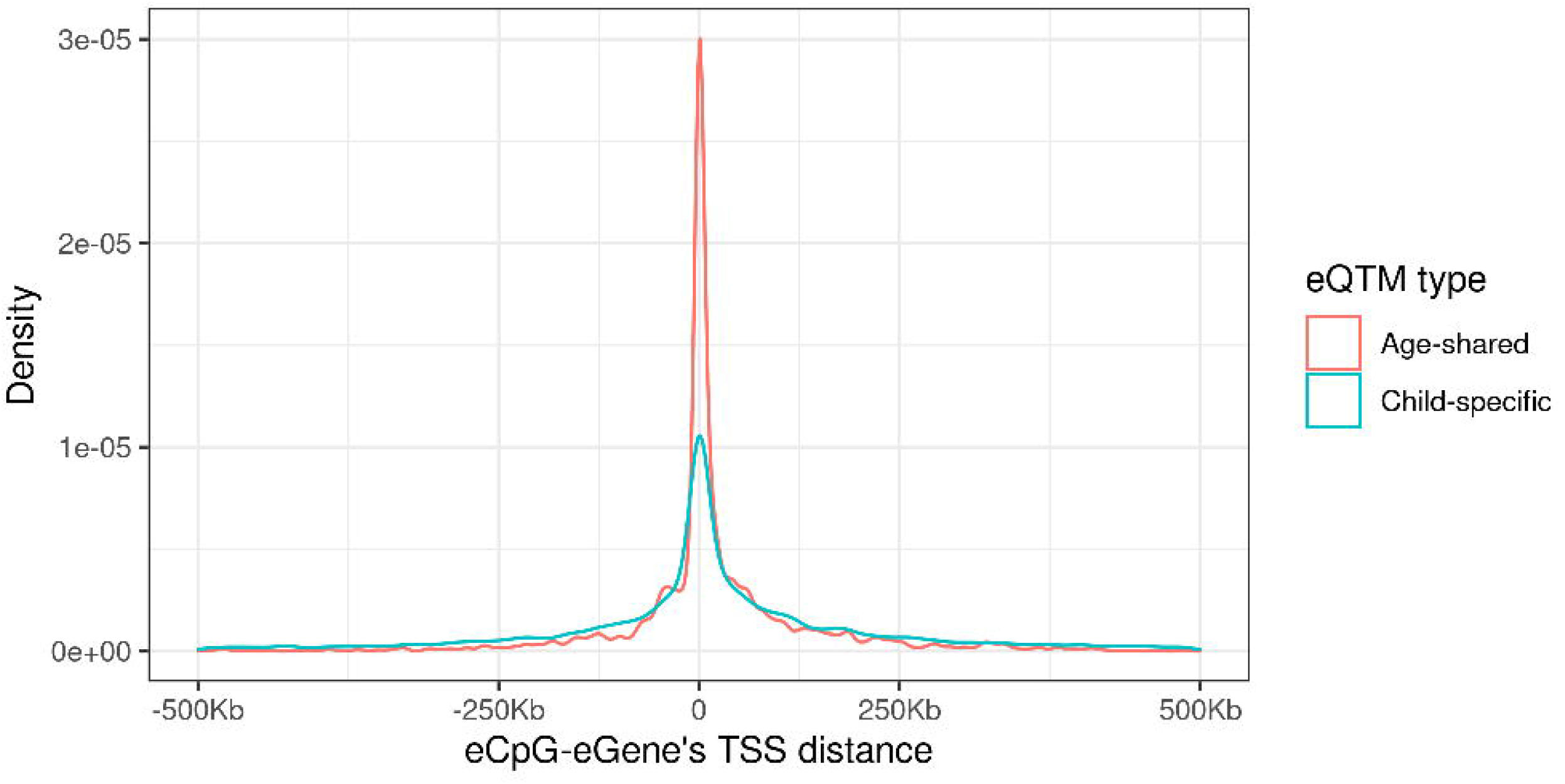
Enrichment of eCpGs with reliable measurements for CpGs with age-variable methylation levels. Only CpGs with reliable measurements (ICC >0.4) were considered (Sugden et al., 2020). eCpGs were classified in all (grey), inverse (yellow); and positive (green). Age-variable CpGs were retrieved from the MeDALL project (from birth to childhood (Xu et al., 2017)) and the Epidelta project (from birth to adolescence (RH et al., 2021)), and they were classified in: variable (CpGs with methylation levels that change with age), decreased (CpGs with methylation levels that decrease with age), and increased (CpGs with methylation levels that increase with age). The y-axis represents the odds ratio (OR) of the enrichment. For all eCpG types, the enrichment was computed against non eCpGs.

**Figure 5 – figure supplement 2.**
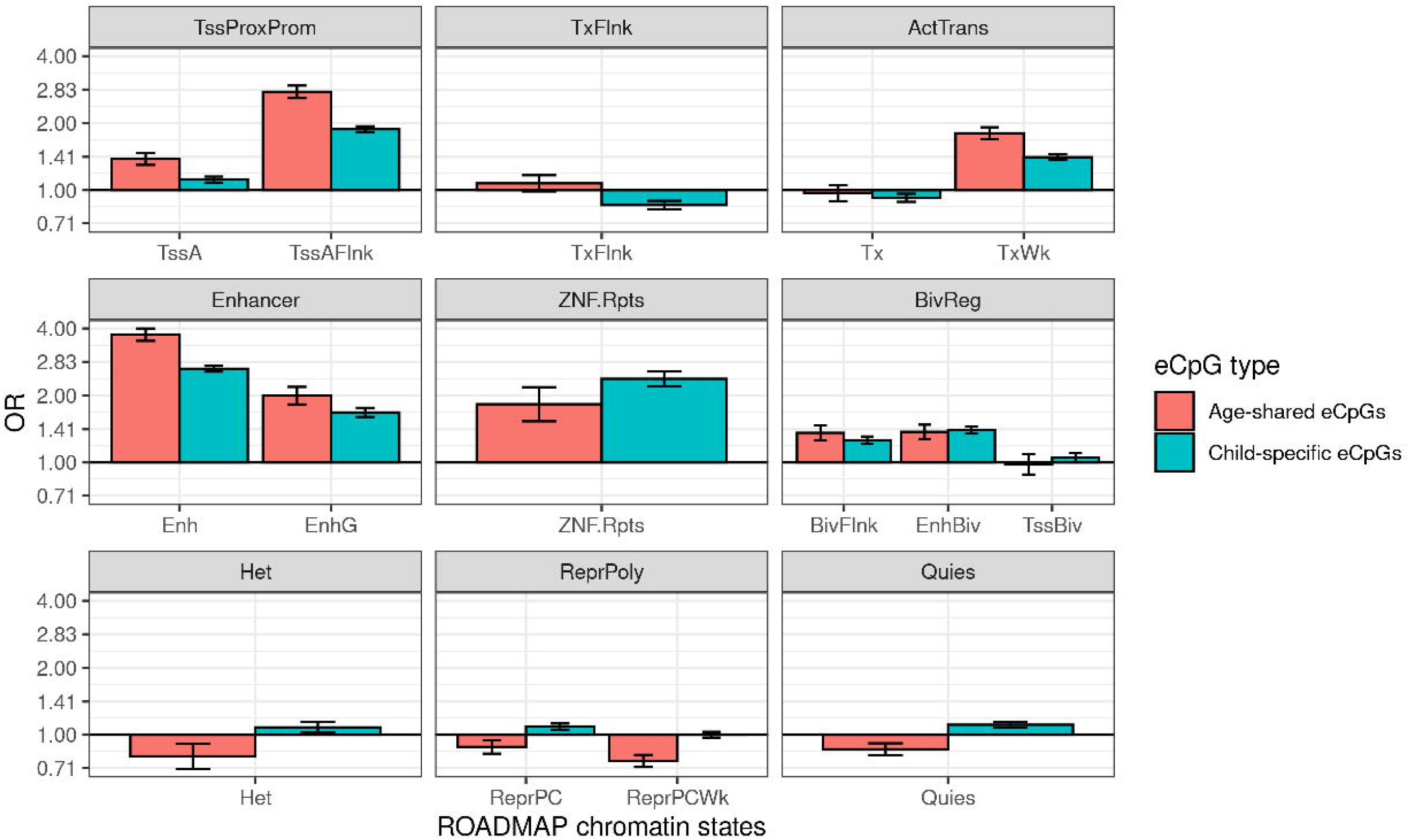
Probe reliability in eCpGs according to overlap with adult eQTMs. eCpGs were classified in age-shared eCpGs (eCpGs identified in HELIX children and also in adults from MESA and/or GTP studies, in red); and child-specific eCpGs (eCpGs only identified in HELIX children and not in the adult cohorts, in blue). Probe reliabilities were based on intraclass correlation coefficients (ICCs) obtained from (Sugden et al., 2020).

**Figure 5 – figure supplement 3.**
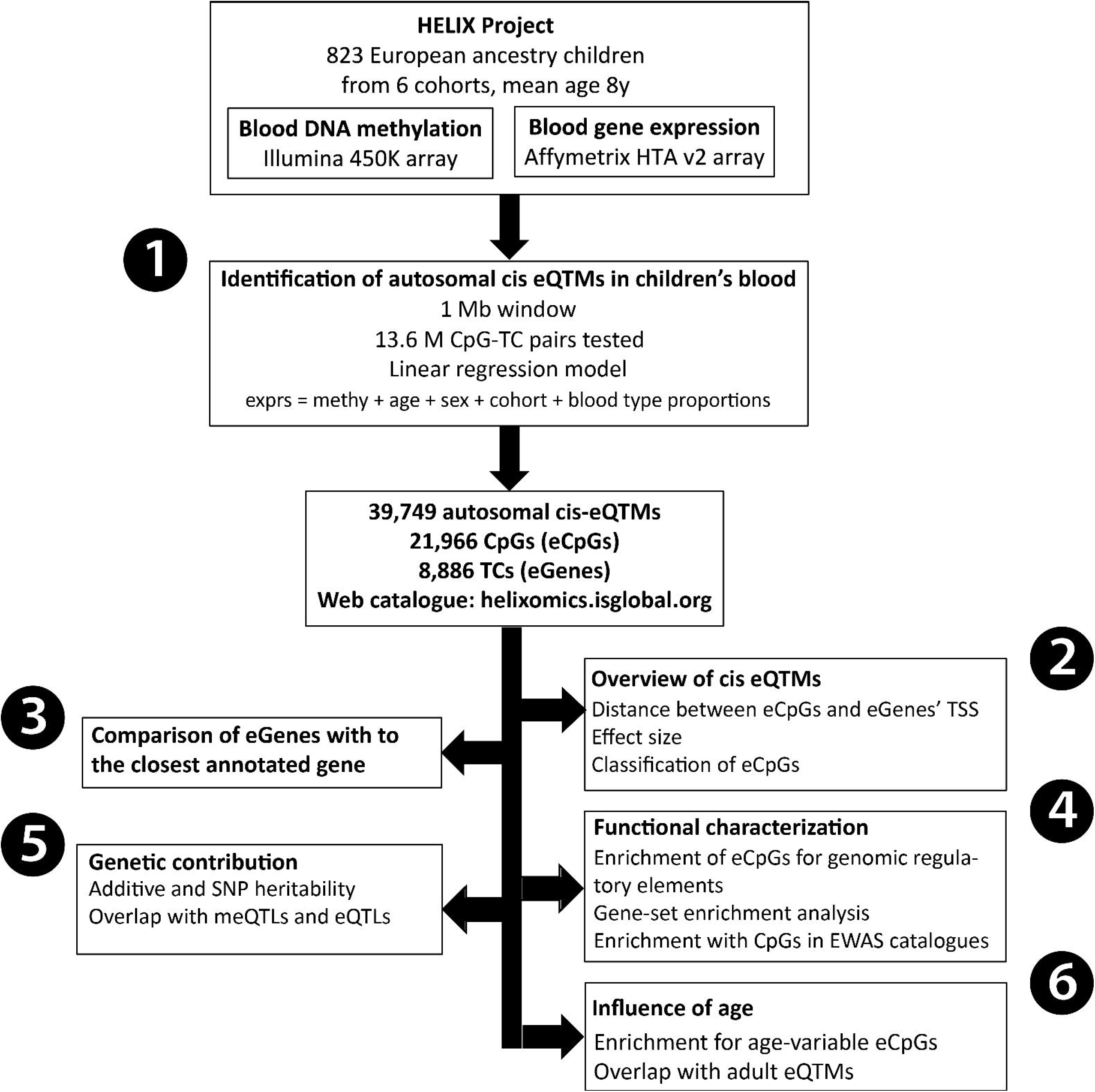
Distribution of the distance between CpG-Gene’s TSS by eQTM type. eQTMs were classified in age-shared (eQTMs identified in HELIX children and also in adults from MESA or GTP studies, in red); and child-specific (eQTMs only identified in HELIX children and not in adult cohorts, in blue). Distance between eCpG and eGene’s TSS is expressed in kb. Age- shared eQTMs median distance: 1.2 kb (IQR: -2.4; 35.4 kb). Child-specific eQTMs median distance: - 1.1 kb (IQR: -39.4; 70.7 kb).

**Figure 5 – figure supplement 4.**
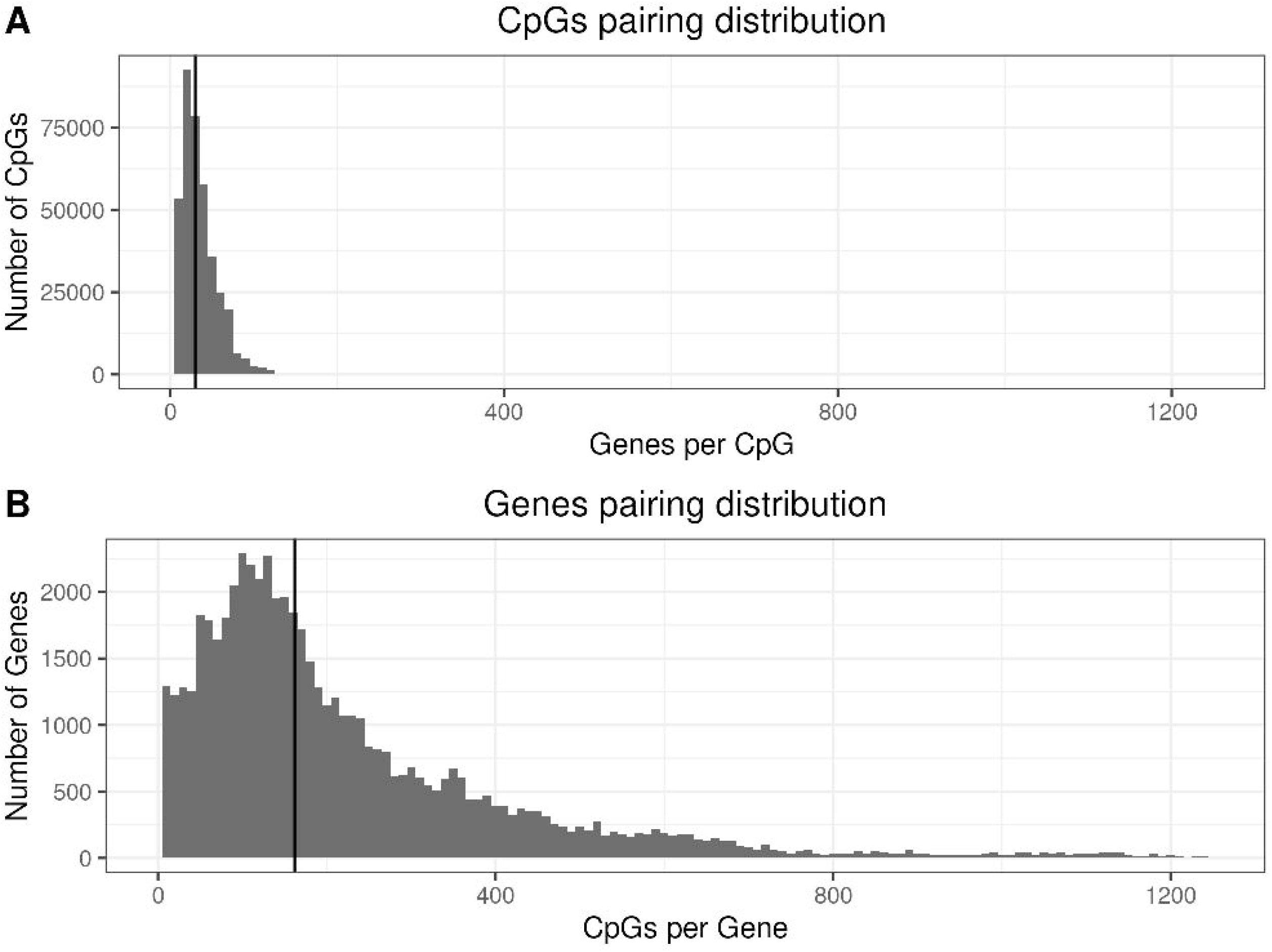
Enrichment of age-shared and child-specific eCpGs for blood ROADMAP blood chromatin states. eCpGs were classified in age-shared (eCpGs identified in HELIX children and also in adults from MESA or GTP studies, in red); and child-specific (eCpGs only identified in HELIX children and not in adult cohorts, in blue). ROADMAP blood chromatin states (Roadmap Epigenomics Consortium et al., 2015) are: active TSS (TssA), flanking active TSS (TssAFlnk), transcription at 5’ and 3’ (TxFlnk), transcription region (Tx), weak transcription region (TxWk), enhancer (Enh); genic enhancer (EnhG), zinc finger genes and repeats (ZNF.Rpts), flanking bivalent region (BivFlnx), bivalent enhancer (EnhBiv), bivalent TSS (TssBiv), heterochromatin (Het), repressed Polycomb (ReprPC), weak repressed Polycomb (ReprPCWk), and quiescent region (Quies). Chromatin states can be grouped in active transcription start site proximal promoter states (TssProxProm), active transcribed states (ActTrans), bivalent regulatory states (BivReg) and repressed Polycomb states (ReprPoly). The y-axis represents the odds ratio (OR) of the enrichment. For each regulatory element, the enrichment was computed against non eCpGs.

## File legends

**Supplementary tables (HELIX_MethExpr_SupTables.xlsx):** File with supplementary tables S1-S4.

**Source code file (SuplementaryCode.zip)**: compressed file with the code used to run the analyses and generate the tables and figures.

## Acknowledgments

The authors acknowledge the contribution of all the HELIX children and their families.

## Funding

The study has received funding from the European Community’s Seventh Framework Programme (FP7/2007-206) under grant agreement no 308333 (HELIX project); the H2020-EU.3.1.2. - Preventing Disease Programme under grant agreement no 874583 (ATHLETE project); from the European Union’s Horizon 2020 research and innovation programme under grant agreement no 733206 (LIFECYCLE project), and from the European Joint Programming Initiative “A Healthy Diet for a Healthy Life” (JPI HDHL and Instituto de Salud Carlos III) under the grant agreement no AC18/00006 (NutriPROGRAM project). The genotyping was supported by the project PI17/01225, funded by the Instituto de Salud Carlos III and co-funded by European Union (ERDF, “A way to make Europe”) and the Centro Nacional de Genotipado-CEGEN (PRB2-ISCIII).

BiB received core infrastructure funding from the Wellcome Trust (WT101597MA) and a joint grant from the UK Medical Research Council (MRC) and Economic and Social Science Research Council (ESRC) (MR/N024397/1). INMA data collections were supported by grants from the Instituto de Salud Carlos III, CIBERESP, and the Generalitat de Catalunya-CIRIT. KANC was funded by the grant of the Lithuanian Agency for Science Innovation and Technology (6-04-2014_31V-66). The Norwegian Mother, Father and Child Cohort Study is supported by the Norwegian Ministry of Health and Care Services and the Ministry of Education and Research. The Rhea project was financially supported by European projects (EU FP6-2003-Food-3-NewGeneris, EU FP6. STREP Hiwate, EU FP7 ENV.2007.1.2.2.2.

Project No 211250 Escape, EU FP7-2008-ENV-1.2.1.4 Envirogenomarkers, EU FP7- HEALTH-2009- single stage CHICOS, EU FP7 ENV.2008.1.2.1.6. Proposal No 226285 ENRIECO, EU- FP7- HEALTH-2012 Proposal No 308333 HELIX), and the Greek Ministry of Health (Program of Prevention of obesity and neurodevelopmental disorders in preschool children, in Heraklion district, Crete, Greece: 2011-2014; “Rhea Plus”: Primary Prevention Program of Environmental Risk Factors for Reproductive Health, and Child Health: 2012-15).

We acknowledge support from the Spanish Ministry of Science and Innovation through the “Centro de Excelencia Severo Ochoa 2019-2023” Program (CEX2018-000806-S), and support from the Generalitat de Catalunya through the CERCA Program.

MV-U and CR-A were supported by a FI fellowship from the Catalan Government (FI-DGR 2015 and #016FI_B 00272). MC received funding from Instituto Carlos III (Ministry of Economy and Competitiveness) (CD12/00563 and MS16/00128).

## Abbreviations

BivFlnx: flanking bivalent region
CpG: cytosine nucleotide followed by a guanine nucleotide
eCpG: CpG whose methylation is associated with gene expression; thus, it is part of an eQTM
eGene: gene whose expression is associated with CpG methylation; thus, it is part of an eQTM
eQTM: expression quantitative trait methylation (statistically significant associations of CpG- gene pairs)
eQTL: expression quantitative trait locus (SNP associated with gene expression)
Enh: enhancer
EnhBiv: bivalent enhancer
EnhG: genic enhancer
EWAS: epigenome-wide association study
FC: fold change
FDR: false discovery rate
GO: gene ontology
GWAS: genome-wide association study
HELIX: Human Early-Life Exposome project
Het: heterochromatin
ICC: intraclass correlation coefficient
IQR: interquartile range
meQTL: methylation quantitative trait locus (SNP associated with DNA methylation)
OR: odds ratio
Quies: quiescent region
ReprPC: repressed Polycomb
ReprPCWk: weak repressed Polycomb
SE: standard error
SNP: single nucleotide polymorphism
TC: transcript cluster
TSS: transcription start site
TssA: active transcription start site
TssAFlnk: flanking active transcription start site
TssBiv: bivalent transcription start site
TSS200: proximal promoter, from TSS to 200 bp
TSS1500: distal promoter, from 200 bp to 1,500 bp
Tx: transcription region
TxFlnk: transcription at 5’ and 3’
TxWk: weak transcription region
3’UTR: 3’ untranslated region
5’UTR: 5’ untranslated region
ZNF.Rpts: zinc finger genes and repeats

## References

Aryee MJ, Jaffe AE, Corrada-Bravo H, Ladd-Acosta C, Feinberg AP, Hansen KD, Irizarry RA. 2014. Minfi: a flexible and comprehensive Bioconductor package for the analysis of Infinium DNA methylation microarrays. Bioinformatics 30:1363–9. doi:10.1093/bioinformatics/btu049

Battram T, Yousefi P, Crawford G, Prince C, Babei MS, Sharp G, Hatcher C, Vega-Salas MJ, Khodabakhsh S, Whitehurst O, Langdon R, Mahoney L, Elliott HR, Mancano G, Lee M, Watkins SH, Lay AC, Hemani G, Gaunt TR, Relton CL, Staley JR, Suderman M. 2021. The EWAS Catalog: a database of epigenome-wide association studies. OSF Prepr 4. doi:10.31219/OSF.IO/837WN

Bonder MJ a., Kasela S, Kals M, Tamm R, Lokk K, Barragan I, Buurman WA, Deelen P, Greve JW, Ivanov M, Rensen SS, van Vliet-Ostaptchouk J V., Wolfs MG, Fu J, Hofker MH, Wijmenga C, Zhernakova A, Ingelman-Sundberg M, Franke L, Milani L. 2014. Genetic and epigenetic regulation of gene expression in fetal and adult human livers. BMC Genomics. doi:10.1186/1471-2164-15-860

Bonder MJ, Luijk R, Zhernakova D V, Moed M, Deelen P, Vermaat M, van Iterson M, van Dijk F, van Galen M, Bot J, Slieker RC, Jhamai PM, Verbiest M, Suchiman HED, Verkerk M, van der Breggen R, van Rooij J, Lakenberg N, Arindrarto W, Kielbasa SM, Jonkers I, van ’t Hof P, Nooren I, Beekman M, Deelen J, van Heemst D, Zhernakova A, Tigchelaar EF, Swertz MA, Hofman A, Uitterlinden AG, Pool R, van Dongen J, Hottenga JJ, Stehouwer CDA, van der Kallen CJH, Schalkwijk CG, van den Berg LH, van Zwet EW, Mei H, Li Y, Lemire M, Hudson TJ, Slagboom PE, Wijmenga C, Veldink JH, van Greevenbroek MMJ, van Duijn CM, Boomsma DI, Isaacs A, Jansen R, van Meurs JBJ, ’t Hoen PAC, Franke L, Heijmans BT, Heijmans BT. 2017. Disease variants alter transcription factor levels and methylation of their binding sites. Nat Genet 49:131–138. doi:10.1038/ng.3721

Buckberry S, Bent SJ, Bianco-Miotto T, Roberts CT. 2014. MassiR: A method for predicting the sex of samples in gene expression microarray datasets. Bioinformatics 30:2084– 2085. doi:10.1093/bioinformatics/btu161

Cavalli G, Heard E. 2019. Advances in epigenetics link genetics to the environment and disease. Nature. doi:10.1038/s41586-019-1411-0

Chang CC, Chow CC, Tellier LC, Vattikuti S, Purcell SM, Lee JJ. 2015. Second-generation PLINK: rising to the challenge of larger and richer datasets. Gigascience 4:7. doi:10.1186/s13742-015-0047-8

Chatzi L, Leventakou V, Vafeiadi M, Koutra K, Roumeliotaki T, Chalkiadaki G, Karachaliou M, Daraki V, Kyriklaki A, Kampouri M, Fthenou E, Sarri K, Vassilaki M, Fasoulaki M, Bitsios P, Koutis A, Stephanou EG, Kogevinas M. 2017. Cohort Profile: The Mother- Child Cohort in Crete, Greece (Rhea Study) 46:1392–1393k. doi:10.1093/ije/dyx084

Das S, Forer L, Schönherr S, Sidore C, Locke AE, Kwong A, Vrieze SI, Chew EY, Levy S, McGue M, Schlessinger D, Stambolian D, Loh P-R, Iacono WG, Swaroop A, Scott LJ, Cucca F, Kronenberg F, Boehnke M, Abecasis GR, Fuchsberger C. 2016. Next- generation genotype imputation service and methods. Nat Genet 48:1284–1287. doi:10.1038/ng.3656

Delahaye F, Do C, Kong Y, Ashkar R, Salas M, Tycko B, Wapner R, Hughes F. 2018. Genetic variants influence on the placenta regulatory landscape. PLoS Genet 14. doi:10.1371/journal.pgen.1007785

Dudbridge F, Gusnanto A. 2008. Estimation of significance thresholds for genomewide association scans. Genet Epidemiol 32:227. doi:10.1002/GEPI.20297

Feinberg AP. 2018. The Key Role of Epigenetics in Human Disease Prevention and Mitigation. N Engl J Med 378:1323–1334. doi:10.1056/nejmra1402513

Felix JF, Joubert BR, Baccarelli AA, Sharp GC, Almqvist C, Annesi-Maesano I, Arshad H, Baiz N, Bakermans-Kranenburg MJ, Bakulski KM, Binder EB, Bouchard L, Breton C V., Brunekreef B, Brunst KJ, Burchard EG, Bustamante M, Chatzi L, Munthe-Kaas MC, Corpeleijn E, Czamara D, Dabelea D, Smith GD, De Boever P, Duijts L, Dwyer T, Eng C, Eskenazi B, Everson TM, Falahi F, Fallin MD, Farchi S, Fernandez MF, Gao L, Gaunt TR, Ghantous A, Gillman MW, Gonseth S, Grote V, Gruzieva O, Håberg SE, Herceg Z, Hivert MF, Holland N, Holloway JW, Hoyo C, Hu D, Huang RC, Huen K, Järvelin MR, Jima DD, Just AC, Karagas MR, Karlsson R, Karmaus W, Kechris KJ, Kere J, Kogevinas M, Koletzko B, Koppelman GH, Kupers LK, Ladd-Acosta C, Lahti J, Lambrechts N, Langie SAS, Lie RT, Liu AH, Magnus MC, Magnus P, Maguire RL, Marsit CJ, McArdle W, Melen E, Melton P, Murphy SK, Nawrot TS, Nisticò L, Nohr EA, Nordlund B, Nystad W, Oh SS, Oken E, Page CM, Perron P, Pershagen G, Pizzi C, Plusquin M, Raikkonen K, Reese SE, Reischl E, Richiardi L, Ring S, Roy RP, Rzehak P, Schoeters G, Schwartz DA, Sebert S, Snieder H, Sørensen TIA, Starling AP, Sunyer J, Taylor JA, Tiemeier H, Ullemar V, Vafeiadi M, Van Ijzendoorn MH, Vonk JM, Vriens A, Vrijheid M, Wang P, Wiemels JL, Wilcox AJ, Wright RJ, Xu CJ, Xu Z, Yang I V., Yousefi P, Zhang H, Zhang W, Zhao S, Agha G, Relton CL, Jaddoe VWV, London SJ. 2018. Cohort profile: Pregnancy and childhood epigenetics (PACE) consortium. Int J Epidemiol 47:22–23u. doi:10.1093/ije/dyx190

Fortin J-P, Fertig E, Hansen K. 2014a. shinyMethyl: interactive quality control of Illumina 450k DNA methylation arrays in R. F1000Research 3:175. doi:10.12688/f1000research.4680.2

Fortin J-P, Labbe A, Lemire M, Zanke BW, Hudson TJ, Fertig EJ, Greenwood C, Hansen KD. 2014b. Functional normalization of 450k methylation array data improves replication in large cancer studies. Genome Biol 15:503. doi:10.1186/s13059-014-0503-2

Fuchsberger C, Abecasis GR, Hinds DA. 2015. Minimac2: Faster genotype imputation. Bioinformatics 31:782–784. doi:10.1093/bioinformatics/btu704

Gamazon ER, Segrè A V., Van De Bunt M, Wen X, Xi HS, Hormozdiari F, Ongen H, Konkashbaev A, Derks EM, Aguet F, Quan J, Nicolae DL, Eskin E, Kellis M, Getz G, McCarthy MI, Dermitzakis ET, Cox NJ, Ardlie KG. 2018. Using an atlas of gene regulation across 44 human tissues to inform complex disease- and trait-associated variation. Nat Genet 50:956–967. doi:10.1038/s41588-018-0154-4

Gaunt TR, Shihab HA, Hemani G, Min JL, Woodward G, Lyttleton O, Zheng J, Duggirala A, McArdle WL, Ho K, Ring SM, Evans DM, Davey Smith G, Relton CL. 2016. Systematic identification of genetic influences on methylation across the human life course. Genome Biol 17:61. doi:10.1186/s13059-016-0926-z

Gondalia R, Baldassari A, Holliday KM, Justice AE, Méndez-Giráldez R, Stewart JD, Liao D, Yanosky JD, Brennan KJM, Engel SM, Jordahl KM, Kennedy E, Ward-Caviness CK, Wolf K, Waldenberger M, Cyrys J, Peters A, Bhatti P, Horvath S, Assimes TL, Pankow JS, Demerath EW, Guan W, Fornage M, Bressler J, North KE, Conneely KN, Li Y, Hou L, Baccarelli AA, Whitsel EA. 2019. Methylome-wide association study provides evidence of particulate matter air pollution-associated DNA methylation. Environ Int 132. doi:10.1016/j.envint.2019.03.071

Grazuleviciene R, Danileviciute A, Nadisauskiene R, Vencloviene J. 2009. Maternal smoking, GSTM1 and GSTT1 polymorphism and susceptibility to adverse pregnancy outcomes. Int J Environ Res Public Health 6:1282–1297. doi:10.3390/ijerph6031282

Gutierrez-Arcelus M, Lappalainen T, Montgomery SB, Buil A, Ongen H, Yurovsky A, Bryois J, Giger T, Romano L, Planchon A, Falconnet E, Bielser D, Gagnebin M, Padioleau I, Borel C, Letourneau A, Makrythanasis P, Guipponi M, Gehrig C, Antonarakis SE, Dermitzakis ET. 2013. Passive and active DNA methylation and the interplay with genetic variation in gene regulation. Elife 2. doi:10.7554/eLife.00523

Gutierrez-Arcelus M, Ongen H, Lappalainen T, Montgomery SB, Buil A, Yurovsky A, Bryois J, Padioleau I, Romano L, Planchon A, Falconnet E, Bielser D, Gagnebin M, Giger T, Borel C, Letourneau A, Makrythanasis P, Guipponi M, Gehrig C, Antonarakis SE, Dermitzakis ET. 2015. Tissue-Specific Effects of Genetic and Epigenetic Variation on Gene Regulation and Splicing. PLoS Genet 11. doi:10.1371/journal.pgen.1004958

Guxens M, Ballester F, Espada M, Fernández MF, Grimalt JO, Ibarluzea J, Olea N, Rebagliato M, Tardón A, Torrent M, Vioque J, Vrijheid M, Sunyer J. 2012. Cohort Profile: the INMA--INfancia y Medio Ambiente--(Environment and Childhood) Project. Int J Epidemiol 41:930–40. doi:10.1093/ije/dyr054

Hansen K. n.d. IlluminaHumanMethylation450kanno.ilmn12.hg19: Annotation for Illumina’s 450k methylation arrays. doi:10.18129/B9.bioc.IlluminaHumanMethylation450kanno.ilmn12.hg19

Heude B, Forhan A, Slama R, Douhaud L, Bedel S, Saurel-Cubizolles M-JJ, Hankard R, Thiebaugeorges O, de Agostini M, Annesi-Maesano I, Kaminski M, Charles M-AA, Annesi-Maesano I, Bernard JY, Botton J, Charles M-AA, Dargent-Molina P, de Lauzon- Guillain B, Ducimetière P, de Agostini M, Foliguet B, Forhan A, Fritel X, Germa A, Goua V, Hankard R, Heude B, Kaminski M, Larroque B, Lelong N, Lepeule J, Magnin G, Marchand L, Nabet C, Pierre F, Slama R, Saurel-Cubizolles M-JJ, Schweitzer M, Thiebaugeorges O, EDEN mother-child cohort study group. 2016. Cohort Profile: The EDEN mother-child cohort on the prenatal and early postnatal determinants of child health and development. Int J Epidemiol 45:353–363. doi:10.1093/ije/dyv151

Houseman EAE, Accomando WP, Koestler DDC, Christensen BBC, Marsit CCJ, Nelson HH, Wiencke JK, Kelsey KTK, Natoli G, Ji H, Ehrlich L, Seita J, Murakami P, Doi A, Lindau P, Lee H, Aryee M, Irizarry R, Kim K, Rossi D, Inlay M, Serwold T, Karsunky H, Ho L, Daley G, Weissman I, Feinberg A, Khavari D, Sen G, Rinn J, Baron U, Turbachova I, Hellwag A, Eckhardt F, Berlin K, Hoffmuller U, Gardina P, Olek S, Wieczorek G, Asemissen A, Model F, Turbachova I, Floess S, Liebenberg V, Baron U, Stauch D, Kotsch K, Pratschke J, Hamann A, Loddenkemper C, Stein H, Volk H, Hoffmuller U, Grutzkau A, Mustea A, Huehn J, Scheibenbogen C, Olek S, Sehouli J, Loddenkemper C, Cornu T, Schwachula T, Hoffmuller U, Grutzkau A, Lohneis P, Dickhaus T, Grone J, Kruschewski M, Mustea A, Turbachova I, Baron U, Olek S, Hanahan D, Weinberg R, Ostrand-Rosenberg S, Lynch L, O’Connell J, Kwasnik A, Cawood T, O’Farrelly C, O’Shea D, Anderson E, Gutierrez D, Hasty A, Chua W, Charles K, Baracos V, Clarke S, Carroll R, Ruppert D, Stefanski L, Gaujoux R, Seoighe C, Gong T, Hartmann N, Kohane I, Brinkmann V, Staedtler F, Letzkus M, Bongiovanni S, Szustakowski J, Shen- Orr S, Tibshirani R, Khatri P, Bodian D, Staedtler F, Perry N, Hastie T, Sarwal M, Davis M, Butte A, Wang S, Petronis A, Smyth G, Leek J, Storey J, Teschendorff A, Zhuang J, Widschwendte R, Goldfarb D, Idnani A, Peters E, McClean M, Liu M, Eisen E, Mueller N, Kelsey KTK, Teschendorff A, Menon U, Gentry-Maharaj A, Ramus S, Gayther S, Apostolidou S, Jones A, Lechner M, Beck S, Jacobs I, Widschwendter M, Kerkel K, Schupf N, Hatta K, Pang D, Salas M, Kratz A, Minden M, Murty V, Zigman W, Mayeux R, Jenkins E, Torkamani A, Schork N, Silverman W, Croy B, Tycko B, Wang X, Zhu H, Snieder H, Su S, Munn D, Harshfield G, Maria B, Dong Y, Treiber F, Gutin B, Shi H, Trellakis S, Bruderek K, Dumitru C, Gholaman H, Gu X, Bankfalvi A, Scherag A, Hutte J, Dominas N, Lehnerdt G, Hoffmann T, Lang S, Brandau S, Kuss I, Hathaway B, Ferris R, Gooding W, Whiteside T, Kuss I, Hathaway B, Ferris R, Gooding W, Whiteside T, Mold J, Venkatasubrahmanyam S, Burt T, Michaelsson J, Rivera J, Galkina S, Weinberg K, Stoddart C, McCune J, Ouden M den, Ubachs J, Stoot J, Wersch J van, Bishara S, Griffin M, Cargill A, Bali A, Gore M, Kaye S, Shepherd J, Trappen P Van, Cho H, Hur H, Kim SS, Kim SS, Kim J, Kim Y, Lee K, Verstegen R, Kusters M, Gemen E, Vries E De, Ram G, Chinen J, Thurston S, Spiegelman D, Ruppert D, Li B, Yin X, Goeman J, Buhlmann P, Subramanian A, Tamayo P, Mootha V, Mukherjeed S, Ebert B, Gillette M, Paulovich A, Pomeroy S, Golub T, Lander E, Mesirov J, Carroll R, Galindo C, Little R, Rubin D, Koestler DDC, Marsit CCJ, Christensen BBC, Karagas M, Bueno R, Sugarbaker D, Kelsey KTK, Houseman EAE, Marsit CCJ, Koestler DDC, Christensen BBC, Karagas M, Houseman EAE, Kelsey KTK, Pedersen K, Bamlet W, Oberg A, Andrade M de, Matsumoto M, Tang H, Thibodeau S, Petersen G, Wang L, Bocklandt S, Lin W, Sehl M, Sanchez F, Sinsheimer J, Horvath S, Vilain E, Chu M, Siegmund K, Hao Q, Crooks G, Tavare S, Shibata D, Doi A, Park I, Wen B, Murakami P, Aryee M, Houseman EAE, Christensen BBC, Yeh R, Marsit CCJ, Karagas M, Alberts B, Johnson A, Lewis J, Raff M, Roberts K, Showe M, Vachani A, Kossenkov A, Yousef M, Nichols C, Kossenkov A, Vachani A, Chang C, Nichols C, Billouin S, Watkins N, Gusnanto A, Bono B de, De S, Miranda-Saavedra D, Ginns L, Goldenheim P, Miller L, Burton R, Gillick L, Mazzoccoli G, Balzanelli M, Giuliani A, Cata A De, Viola M La. 2012. DNA methylation arrays as surrogate measures of cell mixture distribution. BMC Bioinformatics 13:86. doi:10.1186/1471-2105-13-86

Huse SM, Gruppuso PA, Boekelheide K, Sanders JA. 2015. Patterns of gene expression and DNA methylation in human fetal and adult liver. BMC Genomics 16:981. doi:10.1186/s12864-015-2066-3

Husquin LT, Rotival M, Fagny M, Quach H, Zidane N, McEwen LM, MacIsaac JL, Kobor MS, Aschard H, Patin E, Quintana-Murci L. 2018. Exploring the genetic basis of human population differences in DNA methylation and their causal impact on immune gene regulation 06 Biological Sciences 0604 Genetics. Genome Biol 19. doi:10.1186/s13059-018-1601-3

J AA and R. 2010. topGO: topGO: Enrichment analysis for Gene Ontology. No Title.

Johnson ND, Wiener HW, Smith AK, Nishitani S, Absher DM, Arnett DK, Aslibekyan S, Conneely KN. 2017. Non-linear patterns in age-related DNA methylation may reflect CD4+ T cell differentiation. Epigenetics 12:492–503. doi:10.1080/15592294.2017.1314419

Johnson WE, Li C, Rabinovic A. 2007. Adjusting batch effects in microarray expression data using empirical Bayes methods. Biostatistics 8:118–27. doi:10.1093/biostatistics/kxj037

Jones PA. 2012. Functions of DNA methylation: Islands, start sites, gene bodies and beyond. Nat Rev Genet. doi:10.1038/nrg3230

Kennedy EM, Goehring GN, Nichols MH, Robins C, Mehta D, Klengel T, Eskin E, Smith AK, Conneely KN. 2018. An integrated -omics analysis of the epigenetic landscape of gene expression in human blood cells. BMC Genomics 19. doi:10.1186/s12864-018-4842-3

Kim S, Forno E, Zhang R, Park HJ, Xu Z, Yan Q, Boutaoui N, Acosta-Pérez E, Canino G, Chen W, Celedón JC. 2020. Expression Quantitative Trait Methylation Analysis Reveals Methylomic Associations With Gene Expression in Childhood Asthma. Chest. doi:10.1016/j.chest.2020.05.601

Küpers LK, Monnereau C, Sharp GC, Yousefi P, Salas LA, Ghantous A, Page CM, Reese SE, Wilcox AJ, Czamara D, Starling AP, Novoloaca A, Lent S, Roy R, Hoyo C, Breton C V., Allard C, Just AC, Bakulski KM, Holloway JW, Everson TM, Xu CJ, Huang RC, van der Plaat DA, Wielscher M, Merid SK, Ullemar V, Rezwan FI, Lahti J, van Dongen J, Langie SAS, Richardson TG, Magnus MC, Nohr EA, Xu Z, Duijts L, Zhao S, Zhang W, Plusquin M, DeMeo DL, Solomon O, Heimovaara JH, Jima DD, Gao L, Bustamante M, Perron P, Wright RO, Hertz-Picciotto I, Zhang H, Karagas MR, Gehring U, Marsit CJ, Beilin LJ, Vonk JM, Jarvelin MR, Bergström A, Örtqvist AK, Ewart S, Villa PM, Moore SE, Willemsen G, Standaert ARL, Håberg SE, Sørensen TIA, Taylor JA, Räikkönen K, Yang I V., Kechris K, Nawrot TS, Silver MJ, Gong YY, Richiardi L, Kogevinas M, Litonjua AA, Eskenazi B, Huen K, Mbarek H, Maguire RL, Dwyer T, Vrijheid M, Bouchard L, Baccarelli AA, Croen LA, Karmaus W, Anderson D, de Vries M, Sebert S, Kere J, Karlsson R, Arshad SH, Hämäläinen E, Routledge MN, Boomsma DI, Feinberg AP, Newschaffer CJ, Govarts E, Moisse M, Fallin MD, Melén E, Prentice AM, Kajantie E, Almqvist C, Oken E, Dabelea D, Boezen HM, Melton PE, Wright RJ, Koppelman GH, Trevisi L, Hivert MF, Sunyer J, Munthe-Kaas MC, Murphy SK, Corpeleijn E, Wiemels J, Holland N, Herceg Z, Binder EB, Davey Smith G, Jaddoe VWV, Lie RT, Nystad W, London SJ, Lawlor DA, Relton CL, Snieder H, Felix JF. 2019. Meta-analysis of epigenome-wide association studies in neonates reveals widespread differential DNA methylation associated with birthweight. Nat Commun 10. doi:10.1038/s41467-019-09671-3

Lappalainen T, Greally JM. 2017. Associating cellular epigenetic models with human phenotypes. Nat Rev Genet. doi:10.1038/nrg.2017.32

Leek JT, Storey JD, Qiu X, Xiao Y, Gordon A, Yakovlev A, Klebanov L, Yakovlev A, Kerr M, Martin M, Churchill G, Kerr M, Churchill G, Holter N, Mitra M, Maritan A, Cieplak M, Banavar J, Gasch A, Spellman P, Kao C, Carmel-Harel O, Eisen M, Rodwell G, Sonu R, Zahn J, Lund J, Wilhelmy J, Storey J, Xiao W, T L, Tompkins R, Davis R, DeRisi J, Iyer V, Brown P, Brem R, Yvert G, Clinton R, Kruglyak L, Schadt E, Monks S, Drake T, Lusis A, Che N, Tseng G, Oh M, Rohlin L, Liao J, Wong W, Yang Y, Dudoit S, Luu P, Lin D, Peng V, Qui X, Klebanov L, Yakovlev A, Morley M, Molony C, Weber T, Devlin J, Ewens K, Rhodes D, Chinnaiyan A, Nguyen D, Sam K, Tsimelzon A, Li X, Wong H, Amundson S, Bittner M, Chen Y, Trent J, Meltzer P, Lamb J, Crawford E, Peck D, Modell J, Blat I, Dabney A, Storey J, Brem R, Storey J, Whittle J, Kruglyak L, Hedenfalk I, Duggan D, Chen Y, Radmacher M, Bittner M, Storey J, Tibshirani R, Dabney A, Storey J, Rice J, Storey J, Buja A, Eyuboglu N, Lehman E, Romano J, Owen A, Qiu X, Yakovlev A, Efron B, Efron B, Cai G, Sarkar S, Benjamini Y, Yekultieli D, Pawitan Y, Calza S, Ploner A, Yvert G, Brem R, Whittle J, Akey J, Foss E, Eisen M, Spellman P, Brown P, Botstein D, Hedenfalk I, Ringer M, Ben-Dor A, Yakhini Z, Chen Y, Mardia K, Kent J, Bibby J, Alter O, Brown P, Botstein D, Price A, Patterson N, Plenge R, Weinblatt M, SN A, Storey J, Akey J, Kruglyak L, Hastie T, Tibshirani R. 2007. Capturing heterogeneity in gene expression studies by surrogate variable analysis. PLoS Genet 3:1724–1735. doi:10.1371/journal.pgen.0030161

Lehne B, Drong AW, Loh M, Zhang W, Scott WR, Tan S-T, Afzal U, Scott J, Jarvelin M-R, Elliott P, McCarthy MI, Kooner JS, Chambers JC. 2015. A coherent approach for analysis of the Illumina HumanMethylation450 BeadChip improves data quality and performance in epigenome-wide association studies. Genome Biol 16:37. doi:10.1186/s13059-015-0600-x

Leland Taylor D, Jackson AU, Narisu N, Hemani G, Erdos MR, Chines PS, Swift A, Idol J, Didion JP, Welch RP, Kinnunen L, Saramies J, Lakka TA, Laakso M, Tuomilehto J, Parker SCJ, Koistinen HA, Smith GD, Boehnke M, Scott LJ, Birney E, Collins FS. 2019. Integrative analysis of gene expression, DNA methylation, physiological traits, and genetic variation in human skeletal muscle. Proc Natl Acad Sci U S A 166:10883– 10888. doi:10.1073/pnas.1814263116

Li M, Zou D, Li Z, Gao R, Sang J, Zhang Y, Li R, Xia L, Zhang T, Niu G, Bao Y, Zhang Z. 2019. EWAS Atlas: A curated knowledgebase of epigenome-wide association studies. Nucleic Acids Res 47:D983–D988. doi:10.1093/nar/gky1027

Lin X, Teh AL, Chen L, Lim IY, Tan PF, MacIsaac JL, Morin AM, Yap F, Tan KH, Saw SM, Lee YS, Holbrook JD, Godfrey KM, Meaney MJ, Kobor MS, Chong YS, Gluckman PD, Karnani N. 2017. Choice of surrogate tissue influences neonatal EWAS findings. BMC Med 15. doi:10.1186/s12916-017-0970-x

Liu Y, Ding J, Reynolds LM, Lohman K, Register TC, De la Fuente A, Howard TD, Hawkins GA, Cui W, Morris J, Smith SG, Barr RG, Kaufman JD, Burke GL, Post W, Shea S, Mccall CE, Siscovick D, Jacobs DR, Tracy RP, Herrington DM, Hoeschele I. 2013. Methylomics of gene expression in human monocytes. Hum Mol Genet 22:5065–5074. doi:10.1093/hmg/ddt356

Loh PR, Danecek P, Palamara PF, Fuchsberger C, Reshef YA, Finucane HK, Schoenherr S, Forer L, McCarthy S, Abecasis GR, Durbin R, Price AL. 2016. Reference-based phasing using the Haplotype Reference Consortium panel. Nat Genet 48:1443–1448. doi:10.1038/ng.3679

Lu Y, Wang B, Jiang F, Mo X, Wu L, He P, Lu X, Deng F, Lei S. 2019. Multi-omics integrative analysis identified SNP-methylation-mRNA: Interaction in peripheral blood mononuclear cells. J Cell Mol Med 23:4601. doi:10.1111/JCMM.14315

Magnus P, Birke C, Vejrup K, Haugan A, Alsaker E, Daltveit AK, Handal M, Haugen M, Høiseth G, Knudsen GP, Paltiel L, Schreuder P, Tambs K, Vold L, Stoltenberg C. 2016. Cohort Profile Update: The Norwegian Mother and Child Cohort Study (MoBa). Int J Epidemiol 45:382–388. doi:10.1093/ije/dyw029

Maitre L, De Bont J, Casas M, Robinson O, Aasvang GM, Agier L, Andrušaitytė S, Ballester F, Basagaña X, Borràs E, Brochot C, Bustamante M, Carracedo A, De Castro M, Dedele A, Donaire-Gonzalez D, Estivill X, Evandt J, Fossati S, Giorgis-Allemand L, Gonzalez JR, Granum B, Grazuleviciene R, Gützkow KB, Haug LS, Hernandez-Ferrer C, Heude B, Ibarluzea J, Julvez J, Karachaliou M, Keun HC, Krog NH, Lau CHE, Leventakou V, Lyon-Caen S, Manzano C, Mason D, McEachan R, Meltzer HM, Petraviciene I, Quentin J, Roumeliotaki T, Sabido E, Saulnier PJ, Siskos AP, Siroux V, Sunyer J, Tamayo I, Urquiza J, Vafeiadi M, Van Gent D, Vives-Usano M, Waiblinger D, Warembourg C, Chatzi L, Coen M, Van Den Hazel P, Nieuwenhuijsen MJ, Slama R, Thomsen C, Wright J, Vrijheid M. 2018. Human Early Life Exposome (HELIX) study: A European population-based exposome cohort. BMJ Open 8. doi:10.1136/bmjopen-2017-021311

McCarthy S, Das S, Kretzschmar W, Delaneau O, Wood AR, Teumer A, Kang HM, Fuchsberger C, Danecek P, Sharp K, Luo Y, Sidore C, Kwong A, Timpson N, Koskinen S, Vrieze S, Scott LJ, Zhang H, Mahajan A, Veldink J, Peters U, Pato C, van Duijn CM, Gillies CE, Gandin I, Mezzavilla M, Gilly A, Cocca M, Traglia M, Angius A, Barrett JC, Boomsma D, Branham K, Breen G, Brummett CM, Busonero F, Campbell H, Chan A, Chen S, Chew E, Collins FS, Corbin LJ, Smith GD, Dedoussis G, Dorr M, Farmaki A-E, Ferrucci L, Forer L, Fraser RM, Gabriel S, Levy S, Groop L, Harrison T, Hattersley A, Holmen OL, Hveem K, Kretzler M, Lee JC, McGue M, Meitinger T, Melzer D, Min JL, Mohlke KL, Vincent JB, Nauck M, Nickerson D, Palotie A, Pato M, Pirastu N, McInnis M, Richards JB, Sala C, Salomaa V, Schlessinger D, Schoenherr S, Slagboom PE, Small K, Spector T, Stambolian D, Tuke M, Tuomilehto J, Van den Berg LH, Van Rheenen W, Volker U, Wijmenga C, Toniolo D, Zeggini E, Gasparini P, Sampson MG, Wilson JF, Frayling T, de Bakker PIW, Swertz MA, McCarroll S, Kooperberg C, Dekker A, Altshuler D, Willer C, Iacono W, Ripatti S, Soranzo N, Walter K, Swaroop A, Cucca F, Anderson CA, Myers RM, Boehnke M, McCarthy MI, Durbin R, Abecasis G, Marchini J. 2016. A reference panel of 64,976 haplotypes for genotype imputation. Nat Genet 48:1279–1283. doi:10.1038/ng.3643

Melé M, Ferreira PG, Reverter F, DeLuca DS, Monlong J, Sammeth M, Young TR, Goldmann JM, Pervouchine DD, Sullivan TJ, Johnson R, Segrè A V., Djebali S, Niarchou A, Wright FA, Lappalainen T, Calvo M, Getz G, Dermitzakis ET, Ardlie KG, Guigó R. 2015. The human transcriptome across tissues and individuals. Science (80-) 348:660–665. doi:10.1126/science.aaa0355

Pedersen BS, Quinlan AR. 2017. Who’s Who? Detecting and Resolving Sample Anomalies in Human DNA Sequencing Studies with Peddy. Am J Hum Genet 100:406–413. doi:10.1016/j.ajhg.2017.01.017

Purcell S, Neale B, Todd-Brown K, Thomas L, Ferreira MAR, Bender D, Maller J, Sklar P, de Bakker PIW, Daly MJ, Sham PC. 2007. PLINK: A Tool Set for Whole-Genome Association and Population-Based Linkage Analyses. Am J Hum Genet 81:559–575. doi:10.1086/519795

Reinius LE, Acevedo N, Joerink M, Pershagen G, Dahlén SE, Greco D, Söderhäll C, Scheynius A, Kere J. 2012. Differential DNA methylation in purified human blood cells: Implications for cell lineage and studies on disease susceptibility. PLoS One 7:e41361. doi:10.1371/journal.pone.0041361

RH M, A N, CAM C, E W, LC H, AJ S, J R, BT H, TR G, JF F, VWV J, MJ B-K, H T, CL R, MH van Ij, M S. 2021. Epigenome-wide change and variation in DNA methylation in childhood: trajectories from birth to late adolescence. Hum Mol Genet 30:119–134. doi:10.1093/HMG/DDAA280

Roadmap Epigenomics Consortium RE, Kundaje A, Meuleman W, Ernst J, Bilenky M, Yen A, Heravi-Moussavi A, Kheradpour P, Zhang Z, Wang J, Ziller MJ, Amin V, Whitaker JW, Schultz MD, Ward LD, Sarkar A, Quon G, Sandstrom RS, Eaton ML, Wu Y-C, Pfenning AR, Wang X, Claussnitzer M, Liu Y, Coarfa C, Harris RA, Shoresh N, Epstein CB, Gjoneska E, Leung D, Xie W, Hawkins RD, Lister R, Hong C, Gascard P, Mungall AJ, Moore R, Chuah E, Tam A, Canfield TK, Hansen RS, Kaul R, Sabo PJ, Bansal MS, Carles A, Dixon JR, Farh K-H, Feizi S, Karlic R, Kim A-R, Kulkarni A, Li D, Lowdon R, Elliott G, Mercer TR, Neph SJ, Onuchic V, Polak P, Rajagopal N, Ray P, Sallari RC, Siebenthall KT, Sinnott-Armstrong NA, Stevens M, Thurman RE, Wu J, Zhang B, Zhou X, Beaudet AE, Boyer LA, De Jager PL, Farnham PJ, Fisher SJ, Haussler D, Jones SJM, Li W, Marra MA, McManus MT, Sunyaev S, Thomson JA, Tlsty TD, Tsai L-H, Wang W, Waterland RA, Zhang MQ, Chadwick LH, Bernstein BE, Costello JF, Ecker JR, Hirst M, Meissner A, Milosavljevic A, Ren B, Stamatoyannopoulos JA, Wang T, Kellis M. 2015. Integrative analysis of 111 reference human epigenomes. Nature 518:317–30. doi:10.1038/nature14248

Shabalin AA. 2012. Matrix eQTL: Ultra fast eQTL analysis via large matrix operations, Bioinformatics. Bioinformatics. doi:10.1093/bioinformatics/bts163

Sharp GC, Salas LA, Monnereau C, Allard C, Yousefi P, Everson TM, Bohlin J, Xu Z, Huang RC, Reese SE, Xu CJ, Baïz N, Hoyo C, Agha G, Roy R, Holloway JW, Ghantous A, Merid SK, Bakulski KM, Küpers LK, Zhang H, Richmond RC, Page CM, Duijts L, Lie RT, Melton PE, Vonk JM, Nohr EA, Williams-DeVane CL, Huen K, Rifas-Shiman SL, Ruiz-Arenas C, Gonseth S, Rezwan FI, Herceg Z, Ekström S, Croen L, Falahi F, Perron P, Karagas MR, Quraishi BM, Suderman M, Magnus MC, Jaddoe VWV, Taylor JA, Anderson D, Zhao S, Smit HA, Josey MJ, Bradman A, Baccarelli AA, Bustamante M, Håberg SE, Pershagen G, Hertz-Picciotto I, Newschaffer C, Corpeleijn E, Bouchard L, Lawlor DA, Maguire RL, Barcellos LF, Smith GD, Eskenazi B, Karmaus W, Marsit CJ, Hivert MF, Snieder H, Fallin MD, Melén E, Munthe-Kaas MC, Arshad H, Wiemels JL, Annesi-Maesano I, Vrijheid M, Oken E, Holland N, Murphy SK, Sørensen TIA, Koppelman GH, Newnham JP, Wilcox AJ, Nystad W, London SJ, Felix JF, Relton CL. 2017. Maternal BMI at the start of pregnancy and offspring epigenome-wide DNA methylation: Findings from the pregnancy and childhood epigenetics (PACE) consortium. Hum Mol Genet 26:4067–4085. doi:10.1093/hmg/ddx290

Sugden K, Hannon EJ, Arseneault L, Belsky DW, Corcoran DL, Fisher HL, Houts RM, Kandaswamy R, Moffitt TE, Poulton R, Prinz JA, Rasmussen LJH, Williams BS, Wong CCY, Mill J, Caspi A. 2020. Patterns of Reliability: Assessing the Reproducibility and Integrity of DNA Methylation Measurement. Patterns 1:100014. doi:10.1016/j.patter.2020.100014

Tsai PC, Glastonbury CA, Eliot MN, Bollepalli S, Yet I, Castillo-Fernandez JE, Carnero- Montoro E, Hardiman T, Martin TC, Vickers A, Mangino M, Ward K, Pietiläinen KH, Deloukas P, Spector TD, Viñuela A, Loucks EB, Ollikainen M, Kelsey KT, Small KS, Bell JT. 2018. Smoking induces coordinated DNA methylation and gene expression changes in adipose tissue with consequences for metabolic health 06 Biological Sciences 0604 Genetics. Clin Epigenetics 10. doi:10.1186/s13148-018-0558-0

van Dongen J, Nivard MG, Willemsen G, Hottenga J-J, Helmer Q, Dolan C V., Ehli EA, Davies GE, van Iterson M, Breeze CE, Beck S, Hoen PAC’., Pool R, van Greevenbroek MMJ, Stehouwer CDA, Kallen CJH van der, Schalkwijk CG, Wijmenga C, Zhernakova S, Tigchelaar EF, Beekman M, Deelen J, van Heemst D, Veldink JH, van den Berg LH, van Duijn CM, Hofman BA, Uitterlinden AG, Jhamai PM, Verbiest M, Verkerk M, van der Breggen R, van Rooij J, Lakenberg N, Mei H, Bot J, Zhernakova D V., van’t Hof P, Deelen P, Nooren I, Moed M, Vermaat M, Luijk R, Bonder MJ, van Dijk F, van Galen M, Arindrarto W, Kielbasa SM, Swertz MA, van Zwet EW, Isaacs A, Franke L, Suchiman HE, Jansen R, van Meurs JB, Heijmans BT, Slagboom PE, Boomsma DI. 2016. Genetic and environmental influences interact with age and sex in shaping the human methylome. Nat Commun 7:11115. doi:10.1038/ncomms11115

van Iterson M, Tobi EW, Slieker RC, den Hollander W, Luijk R, Slagboom PE, Heijmans BT. 2014. MethylAid: Visual and interactive quality control of large Illumina 450k data sets. Bioinformatics 30:3435–3437. doi:10.1093/bioinformatics/btu566

Wagner JR, Busche S, Ge B, Kwan T, Pastinen T, Blanchette M. 2014. The relationship between DNA methylation, genetic and expression inter-individual variation in untransformed human fibroblasts. Genome Biol 15. doi:10.1186/gb-2014-15-2-r37

Wright J, Small N, Raynor P, Tuffnell D, Bhopal R, Cameron N, Fairley L, A Lawlor D, Parslow R, Petherick ES, Pickett KE, Waiblinger D, West J. 2013. Cohort profile: The born in bradford multi-ethnic family cohort study. Int J Epidemiol 42:978–991. doi:10.1093/ije/dys112

Wu Y, Zeng J, Zhang F, Zhu Z, Qi T, Zheng Z, Lloyd-Jones LR, Marioni RE, Martin NG, Montgomery GW, Deary IJ, Wray NR, Visscher PM, McRae AF, Yang J. 2018. Integrative analysis of omics summary data reveals putative mechanisms underlying complex traits. Nat Commun 9. doi:10.1038/s41467-018-03371-0

Xu C-J, Bonder MJ, Söderhäll C, Bustamante M, Baïz N, Gehring U, Jankipersadsing SA, van der Vlies P, van Diemen CC, van Rijkom B, Just J, Kull I, Kere J, Antó JM, Bousquet J, Zhernakova A, Wijmenga C, Annesi-Maesano I, Sunyer J, Melén E, Li Y, Postma DS, Koppelman GH. 2017. The emerging landscape of dynamic DNA methylation in early childhood. BMC Genomics 18:25. doi:10.1186/s12864-016-3452-1

