## SupplementalCode for "Identification of autosomal cis expression quantitative trait methylation (cis eQTMs) in children’s blood": 1.createListCommonIDs.R

```
#'##############################################################################
#' Create list of common IDs between methylation and gene Expression
#'##############################################################################

# Load libraries
library(minfi)
library(SummarizedExperiment)
library(openxlsx)

# Load Methylation data ####
load("./data/methylome_subcohort_ComBatSlide_6cells_notfitr_v3.Rdata")
meth <- methylome_subcohort_ComBatSlide_6cells_notfitr

# Load Gene Expression data ####
load("data/transcriptome_subcohort_notfitr_inclsex_v3.RData")
gexp <- transcriptome_subcohort_notfitr_inclsex

## Load ethnicity from genotypes
ethni <- read.xlsx("data/HELIX_GWAS_1000G_ethnicity_20181212.xlsx", sheet = "Prob_ancestry",
                   startRow = 5)
rownames(ethni) <- ethni$sample_id

# Filter Gene Expression dataset
## Discard panel 1B samples
gexp_sel <- gexp[, gexp$Period != "1B"]

## Select European
gexp_samps <- colnames(gexp_sel)

### Select individuals with EUR ancestry from peddy. 
selEUR <- ethni[gexp_samps, "FINAL_ancestry"] == "EUR"
names(selEUR) <- gexp_samps

### If not available, select individuals based on questionary
selEUR[is.na(selEUR)] <- gexp_sel[, names(selEUR[is.na(selEUR)])]$h_ethnicity_cauc == "yes"

selEUR <- selEUR[selEUR]

## Select common samples between Exprs and Methylation
comIds <- intersect(names(selEUR), colnames(meth))
save(comIds, file = "results/preprocessFiles/comIds.Rdata")
```
