## SupplementalCode for "Identification of autosomal cis expression quantitative trait methylation (cis eQTMs) in children’s blood": 2.cromatin.R

```
###############################################################
# Chromatin STATE (15-STATE)
###############################################################

## Run in home
crom <- read.delim("~/data/WS_HELIX/HELIX_preproc/methylation/annotation/Illumina450K_MQtlMappingFile_Original37_v.1.2_incStates15.txt", header = T)
sort(names(crom))

# Searching function:
# Searches if there is 'overlapping' in at least one cell type
anyt <- function(x) {
  any(grepl("overlapping", x))
}

# TssA
tssa <- crom[,grep("TssA", colnames(crom))]
tssa <- tssa[,-grep("TssAFlnk", colnames(tssa))]
tssa <- apply(tssa, 1, anyt)

# TssAFlnk
tssaf <- crom[,grep("TssAFlnk", colnames(crom))]
tssaf <- apply(tssaf, 1, anyt)

# TxFlnk
txf <- crom[,grep("TxFlnk", colnames(crom))]
txf <- apply(txf, 1, anyt)

# TxWk
txw <- crom[,grep("TxWk", colnames(crom))]
txw <- apply(txw, 1, anyt)

# Tx
tx <- crom[,grep("Tx", colnames(crom))]
tx <- tx[,-grep("TxWk|TxFlnk", colnames(tx))]
tx <- apply(tx, 1, anyt)

# EnhG
enhg <- crom[,grep("EnhG", colnames(crom))]
enhg <- apply(enhg, 1, anyt)

# Enh
enh <- crom[,grep("Enh", colnames(crom))]
enh <- enh[,-grep("EnhG", colnames(enh))]
enh <- enh[,-grep("EnhBiv", colnames(enh))]
enh <- apply(enh, 1, anyt)

# ZNF/Rpts
zr <- crom[,grep("ZNF.Rpts", colnames(crom))]
zr <- apply(zr, 1, anyt)

# Het
het <- crom[,grep("Het", colnames(crom))]
het <- apply(het, 1, anyt)

# TssBiv
tssb <- crom[,grep("TssBiv", colnames(crom))]
tssb <- apply(tssb, 1, anyt)

# BivFlnk
bf <- crom[,grep("BivFlnk", colnames(crom))]
bf <- apply(bf, 1, anyt)

# EnhBiv
eb <- crom[,grep("EnhBiv", colnames(crom))]
eb <- apply(eb, 1, anyt)

# ReprPC
rp <- crom[,grep("ReprPC", colnames(crom))]
rp <- rp[,-grep("ReprPCWk", colnames(rp))]
rp <- apply(rp, 1, anyt)

# ReprPCWk
rpcwk <- crom[,grep("ReprPCWk", colnames(crom))]
rpcwk <- apply(rpcwk, 1, anyt)

# Quies
quies <- crom[,grep("Quies", colnames(crom))]
quies <- apply(quies, 1, anyt)

# Final table
crom.df1 <- crom[,2:7]
crom.df2 <- data.frame("TssA" = tssa, "TssAFlnk" = tssaf, "TxFlnk" = txf,
	"TxWk" = txw, "Tx" = tx, "EnhG" = enhg, "Enh" = enh, "ZNF/Rpts" = zr,
	"Het" = het, "TssBiv" = tssb, "BivFlnk" = bf, "EnhBiv" = eb, "ReprPC" = rp,
	"ReprPCWk" = rpcwk, "Quies" = quies)
crom.df <- cbind(crom.df1, crom.df2)
write.csv(crom.df, "data/crom15.csv")
```
