## SupplementalCode for "Identification of autosomal cis expression quantitative trait methylation (cis eQTMs) in children’s blood": 3.dhs.R

```
###############################################################
# DHS
###############################################################

dhs <- read.delim("/Lacie_CRW10023/HELIX_preproc/methylation/annotation/Illumina450K_MQtlMappingFile_Original37_v.1.2_HistoneMarkAndDNASE1.txt", header = T)
sort(names(dhs))

# Searching function:
# Searches if there is 'overlapping' in at least one cell type
anyt <- function(x) {
  any(grepl("overlapping", x))
}

# DNase
dnase <- dhs[,grep("DNase.", colnames(dhs))]
dnase <- dnase[,-grep("DNase.macs2", colnames(dnase))]
ndnase <- gsub("DNase.", "", names(dnase))
dnase <- apply(dnase, 1, anyt)

# DNase.macs2
dnm <- dhs[,grep("DNase.macs2.", colnames(dhs))]
ndnm <- gsub("DNase.macs2.", "", names(dnm))
dnm <- apply(dnm, 1, anyt)

# H2A.Z
h2a <- dhs[,grep("H2A.Z.", colnames(dhs))]
nh2a <- gsub("H2A.Z.", "", names(h2a))
h2a <- apply(h2a, 1, anyt)

# H3K27ac
k27ac <- dhs[,grep("H3K27ac.", colnames(dhs))]
nk27ac <- gsub("H3K27ac.", "", names(k27ac))
k27ac <- apply(k27ac, 1, anyt)

# H3K27me3
k27me3 <- dhs[,grep("H3K27me3.", colnames(dhs))]
nk27me3 <- gsub("H3K27me3.", "", names(k27me3))
k27me3 <- apply(k27me3, 1, anyt)

# H3K36me3
k36me3 <- dhs[,grep("H3K36me3.", colnames(dhs))]
nk36me3 <- gsub("H3K36me3.", "", names(k36me3))
k36me3 <- apply(k36me3, 1, anyt)

# H3K4me1
k4me1 <- dhs[,grep("H3K4me1.", colnames(dhs))]
nk4me1 <- gsub("H3K4me1.", "", names(k4me1))
k4me1 <- apply(k4me1, 1, anyt)

# H3K4me2
k4me2 <- dhs[,grep("H3K4me2.", colnames(dhs))]
nk4me2 <- gsub("H3K4me2.", "", names(k4me2))
k4me2 <- apply(k4me2, 1, anyt)

# H3K4me3
k4me3 <- dhs[,grep("H3K4me3.", colnames(dhs))]
nk4me3 <- gsub("H3K4me3.", "", names(k4me3))
k4me3 <- apply(k4me3, 1, anyt)

# H3K79me2
k79me2 <- dhs[,grep("H3K79me2.", colnames(dhs))]
nk79me2 <- gsub("H3K79me2.", "", names(k79me2))
k79me2 <- apply(k79me2, 1, anyt)

# H3K9ac
k9ac <- dhs[,grep("H3K9ac.", colnames(dhs))]
nk9ac <- gsub("H3K9ac.", "", names(k9ac))
k9ac <- apply(k9ac, 1, anyt)

# H3K9me3
k9me3 <- dhs[,grep("H3K9me3.", colnames(dhs))]
nk9me3 <- gsub("H3K9me3.", "", names(k9me3))
k9me3 <- apply(k9me3, 1, anyt)

# H4K20me1
k20me1 <- dhs[,grep("H4K20me1.", colnames(dhs))]
nk20me1 <- gsub("H4K20me1.", "", names(k20me1))
k20me1 <- apply(k20me1, 1, anyt)

# Final table
dhs.df1 <- dhs[,2:7]
dhs.df2 <- data.frame("DNase" = dnase, "DNase.macs2" = dnm, "H2A.Z" = h2a,
                      "H3K27ac" = k27ac, "H3K27me3" = k27me3, "H3K36me3" = k36me3,
                      "H3K4me1" = k4me1, "H3K4me2" = k4me2, "H3K4me3" = k4me3,
                      "H3K79me2" = k79me2, "H3K9ac" = k9ac, "H3K9me3" = k9me3,
                      "H4K20me1" = k20me1)

dhs.df <- cbind(dhs.df1, dhs.df2)
write.csv(dhs.df, "dhs.csv")
```
