## SupplementalCode for "Identification of autosomal cis expression quantitative trait methylation (cis eQTMs) in children’s blood": 4.createGRSet.R

```
###############################################################################
#' Create GenomicRatioSet for analysis (methylation)
###############################################################################

library('minfi', verbose = FALSE)

# Loading GRSet
load("./data/methylome_subcohort_ComBatSlide_notfitr_v4.Rdata")
gset <- methylome_subcohort_ComBatSlide_notfitr

## Load common IDs
load("results/preprocessFiles/comIds.Rdata")

# Select common IDs
gset <- gset[ , comIds]

## Remove crosshibridizing and SNPs probes
load("./data/badProbes.Rdata")
gset <- gset[!rownames(gset) %in% c(crossProbes, SNPsProbes), ]

save(gset, file = "results/preprocessFiles/Methylation_GRSet.RData")
```
