## SupplementalCode for "Identification of autosomal cis expression quantitative trait methylation (cis eQTMs) in children’s blood": 5.createSummExp.R

```
###############################################################################
#' Create SummarizedExperiment for analysis (Gene Expression)
###############################################################################

library(Biobase, verbose = FALSE)
library(SummarizedExperiment, verbose = FALSE)
library(sva, verbose = FALSE)
library(isva, verbose = FALSE)
library(limma, verbose = FALSE)    
library(SmartSVA, verbose = FALSE) 

## Load SummarizedExperiment
load("./data/transcriptome_subcohort_notfitr_inclsex_v3.RData")
gexp <- transcriptome_subcohort_notfitr_inclsex

## Load common IDs
load("results/preprocessFiles/comIds.Rdata")

# Select common IDs
gexp <- gexp[ , comIds]

## Remove probes with low call rate
gexp <- gexp[fData(gexp)$fil1 == "no", ]

# Making SE
se <- makeSummarizedExperimentFromExpressionSet(gexp)

## Change ranges to be TSS
gr <- rowRanges(se)
gr$TC_Start <- start(gr)
gr$TC_end <- end(gr)
start(gr) <- gr$TSS_Affy
end(gr) <- gr$TSS_Affy
rowRanges(se) <- gr

save(se, file = "results/preprocessFiles/Expression_SE_raw.RData")

## Compute Residuals ####
mat <- assay(se)
pd <- colData(se)

# Protegim les variables que llavors voldrem eliminar a traves dels residuals
Y.r <- t(resid(lm(t(mat) ~ cohort + e3_sex + age_sample_years +
                    NK_6 + Bcell_6 + CD4T_6 + CD8T_6 + Gran_6 +
                    Mono_6, data = pd)))

# Calculem quantes SVs necessitem
n.sv <- EstDimRMT(Y.r, FALSE)$dim + 1
print(n.sv)
# 57 SVs

# Busquem les SVs
mod <- model.matrix(~ cohort + e3_sex + age_sample_years +
                      NK_6 + Bcell_6 + CD4T_6 + CD8T_6 + Gran_6 +
                      Mono_6, data = pd)
sv.obj <- smartsva.cpp(mat, mod, mod0 = NULL, n.sv = n.sv)

# Residuals de només els SVs.
res <- residuals(lmFit(mat, sv.obj$sv), mat)

# Save
assay(se) <- res
save(se, file = "results/preprocessFiles/Expression_SE_residuals.RData")
```
