## SupplementalCode for "Identification of autosomal cis expression quantitative trait methylation (cis eQTMs) in children’s blood": 6.createAnnotation.R

```
#'##############################################################################
#' Create files with annotation for analysis
#'##############################################################################

library(GenomicRanges)
library(SummarizedExperiment)
library(minfi)
library(dplyr)
library(matrixStats)
library(readxl)

# DNA Methylation Annotation ####
load("results/preprocessFiles/Methylation_GRSet.RData")
rd <- getAnnotation(gset)
rd <- rd[,-c(5:14, 16, 20:21)]

## Add bimodal data
gaps <- gaphunter(gset)

### Mark CpGs having a bimodal distribution containing at least 20% of samples
bimod <- rownames(gaps$proberesults)[rowMaxs(data.matrix(gaps$proberesults)) < 0.8*814]
rd$bimod <- rownames(rd) %in% bimod
### Almost all bimodal CpGs are in sexual chromosomes!!!!

## Add median methylation
median_meth <- rowMedians(getBeta(gset))
names(median_meth) <- rownames(gset)
rd$median <- median_meth[rownames(rd)]

### Categorize in three categories (PMID:26589361)
rd$median_cat <- cut(rd$median, c(0, 0.3, 0.7, 1), labels = c("low", "medium", "high"))

## Mark invariant CpGs (PMID:29202839)
ranges <- apply(getBeta(gset), 1, function(x) {
  a <- quantile(x, c(0.99, 0.01))
  a[1] - a[2]
  })
rd$meth_range <- ranges
rd$variability <- ifelse(ranges > 0.05, "variant", "invariant")


# Adding phantom summary
rd$Phantom_S <- ifelse(grepl("low", rd$Phantom), "low",
                       ifelse(grepl("high", rd$Phantom),"high", ""))

# Adding group columns
rd$TSS200 <- ifelse(grepl("TSS200", rd$UCSC_RefGene_Group), T, F)
rd$TSS1500 <- ifelse(grepl("TSS1500", rd$UCSC_RefGene_Group), T, F)
rd$UTR5 <- ifelse(grepl("5'UTR", rd$UCSC_RefGene_Group), T, F)
rd$FirstExon <- ifelse(grepl("1stExon", rd$UCSC_RefGene_Group), T, F)
rd$Body <- ifelse(grepl("Body", rd$UCSC_RefGene_Group), T, F)
rd$UTR3 <- ifelse(grepl("3'UTR", rd$UCSC_RefGene_Group), T, F)

# Adding dhs and crom15
crom15 <- read.csv("data/crom15.csv", header = T, row.names = 1)
crom15 <- crom15[,c(1,7:21)]
dhs <- read.csv("data/dhs.csv", header = T, row.names = 1)
dhs <- dhs[,c(1,7:19)]

states <- merge(crom15, dhs, by = "HT12v4.ArrayAddress", sort = F)
rd <- merge(rd, states, by.x = "row.names", by.y = "HT12v4.ArrayAddress", sort = F)

a <- strsplit(rd$UCSC_RefGene_Name, ";")
a[a == "character(0)"] <- ""
rd$UCSC_RefGene_Name <- a

# Add reliability
rel <- read_xlsx("data/PMID32885222_reliability.xlsx", skip = 2) %>%
  mutate(Name = `Illumina Probe ID`) %>%
  dplyr::select(Name, Reliability) 

rd <- merge(rd, rel, by = "Name")
methyAnnot <- rd
save(methyAnnot, file = "results/preprocessFiles/methyAnnotation.Rdata")

# Gene Expression Annotation ####
load("results/preprocessFiles/Expression_SE_residuals.RData")
annot <- read.csv("/home/isglobal.lan/cruiz/data/WS_HELIX/HELIX_preproc/gene_expression/annotation/HTA-2_0.na36.hg19.transcript.csv", comment.char="#", as.is = TRUE)

### Mark as coding those TC having data in Swiss-Prot
### Consider expressed TCs with call rate > 10%
expAnnot <- annot %>%
  as_tibble() %>%
  right_join(as_tibble(mcols(rowRanges(se))[, c(1, 7:11, 20)]), by = "transcript_cluster_id") %>%
  mutate(Coding = ifelse(swissprot != "---", "coding", "non-coding"), 
         Expressed = ifelse(CallRate > 10, "Expressed", "Not-Expressed"), 
         TC = transcript_cluster_id)
save(expAnnot, file = "results/preprocessFiles/gexpAnnotation.Rdata")


# Overlaps ####
gr <- makeGRangesFromDataFrame(expAnnot)
gr$TC <- expAnnot$TC
start(gr) <- expAnnot$TSS_Affy
end(gr) <- expAnnot$TSS_Affy
allOver <- findOverlaps(gr + 5e5, rowRanges(gset))
overDF <- data.frame(CpG = rownames(gset)[to(allOver)], TC = gr$TC[from(allOver)], 
                     CpG_Pos = start(rowRanges(gset))[to(allOver)], 
                     TC_Pos = start(gr)[from(allOver)], 
                     TC_strand = strand(gr)[from(allOver)])
overDF$Distance <- ifelse(overDF$TC_strand == "+", overDF$CpG_Pos - overDF$TC_Pos, 
                          overDF$TC_Pos - overDF$CpG_Pos)
save(overDF, file = "results/preprocessFiles/allOverlaps.Rdata")
```
