## SupplementalCode for "Identification of autosomal cis expression quantitative trait methylation (cis eQTMs) in children’s blood": README

These scripts contains the files to prepare our data to run the eQTM analysis. They can be run in this order. The main idea is to generate a GenomicRatioSet with the DNA methylation and a SummarizedExperiment with the gene expression data, having the same individuals and in the same order. These scripts are very specific to our data.

1.createListCommonIDs.R: Select individuals in common between DNA methylation and gene expression. It also selects European individuals. 
2.cromatin.R: Preprocess data from chromatin states to be added to methylation data.
3.dhs.R: Preprocess data from histones accessibility.
4.createGRSet.R: Converts DNA methylation data to a GenomicRatioSet and filters individuals.
5.createSummExp.R: Converts gene expression data to a SummarizedExperiments and filters individuals.
6.createAnnotation.R: Creates annotation for DNA methylation and gene expression objects.
README: this file
