## SupplementalCode for "Identification of autosomal cis expression quantitative trait methylation (cis eQTMs) in children’s blood": 1.mainModel_splitDataByChr.sh

```
#!/bin/sh

###############################################################################
# Methylation vs Expression Analysis using:
#   - Methylation: ComBat
#   - Expression: SVA Residuals protecting age, sex, cohort and cell counts
#   - Model: adjust for age, sex, cohort
#   - All Samples
#   - Autosome chromosomes
###############################################################################

#set the job name
#SBATCH --job-name=noCell

#set the number of CPUS per task
#SBATCH --ntasks-per-node=1

#set the memory
#SBATCH --mem=15000

# job output file information
#SBATCH -o nocellPrepare.out

# job errors file
#SBATCH -e nocellPrepare.err

# set the partition where the job will run
#SBATCH --partition=normal
 
# set the number of nodes
#SBATCH --nodes=1
 
# set max wallclock time
#SBATCH --time=1:00:00
 
# mail alert at start, end and abortion of execution
#SBATCH --mail-type=END
 
# send mail to this address
#SBATCH --mail-user=

module load R/3.5.1-foss-2018b

## Define paths to omic files
meth="results/preprocessFiles/Methylation_GRSet.RData"
gexp="results/preprocessFiles/Expression_SE_residuals.RData"
resFolder="results/MethComBatExpResidualsNoCellAdj"

methInput="${resFolder}/methyInput.Rdata"
gexpInput="${resFolder}/gexpInput.Rdata"

## Create folder for result
mkdir $resFolder
  

## Generate Input data for analysis
R CMD BATCH --vanilla '--args methy="'$meth'" exprs="'$gexp'" out_fold="'$resFolder'" autosomes' src/createInputFiles.R  $resFolder/input.out

## Divide data for analysis
R CMD BATCH --vanilla '--args methy="'$methInput'" exprs="'$gexpInput'" out_fold="'$resFolder'"' src/divide_data.R  $resFolder/divide.out
```
