## SupplementalCode for "Identification of autosomal cis expression quantitative trait methylation (cis eQTMs) in children’s blood": 2.mainModel_runPermutations.sh

```
#!/bin/sh

###############################################################################
# Run linear models for all the different models
###############################################################################

#set the job name
#SBATCH --job-name=nocellSims

#set the number of CPUS per task
#SBATCH --ntasks-per-node=16

#set the memory
#SBATCH --mem=35000

# job output file information
#SBATCH -o nocellSims.out

# job errors file
#SBATCH -e nocellSims.err

# set the partition where the job will run
#SBATCH --partition=normal
 
# set the number of nodes
#SBATCH --nodes=1
 
# set max wallclock time
#SBATCH --time=7:00:00
 
# mail alert at start, end and abortion of execution
#SBATCH --mail-type=END
 
# send mail to this address
#SBATCH --mail-user=

module load R/3.5.1-foss-2018b

## No cell adjusted
echo No Cell Adjusted
resFolder="results/MethComBatExpResidualsNoCellAdj"

simFolder="$resFolder/sim${SLURM_ARRAY_TASK_ID}"
mkdir $simFolder


for i in {1..22}
do
  echo $i
  R CMD BATCH '--args data_fold="'$resFolder'" chr="chr'$i'" model="nocell" out_fold="'$resFolder'" "'${SLURM_ARRAY_TASK_ID}'"' src/runLinearModelSubset.R $simFolder/modchr$i.out
done
mv $simFolder /gpfs42/projects/lab_helix_omics/shared_data/methExprsHELIX/$resFolder
```
