## SupplementalCode for "Identification of autosomal cis expression quantitative trait methylation (cis eQTMs) in children’s blood": 3.mainModel_geteQTMs.sh

```
#!/bin/sh

###############################################################################
# Methylation vs Expression Analysis using:
#   - Methylation: ComBat
#   - Expression: SVA Residuals protecting age, sex, cohort and cell counts
#   - Model: adjust for age, sex, cohort
#   - All Samples
#   - Autosome chromosomes
###############################################################################

resFolder="results/MethComBatExpResidualsNoCellAdj"

## Run linear models
for i in {1..22}
do
  echo $i
  Rscript src/runLinearModelSubset.R data_fold="'$resFolder'" chr="'chr$i'" model="'nocell'" out_fold="'$resFolder'"
done

## Generate beta dsitributions per CpG
Rscript src/getBetaDistributionsCpGs.R folder="'$resFolder'" type="'autosome'"
Rscript src/getSignificantPairs.R resFolder="'$resFolder'" base="'cpgs'" distribution="'$resFolder/CpGsDistr.Rdata'"
```
