## SupplementalCode for "Identification of autosomal cis expression quantitative trait methylation (cis eQTMs) in children’s blood": 7.meQTL_analysis.R

```
#'##############################################################################
#'##############################################################################
#' Script to run eQTM analysis
#'  - Adjust model for age, sex, cohort, cell types and 10 genetic PCs
#'##############################################################################
#'##############################################################################

## Run PCA
plink --bfile ~/data/WS_HELIX/HELIX_preproc/gwas/Final_data_HRCimp_QC2/HELIX.impQC.rs --pca --out ~/data/WS_HELIX/HELIX_analyses/expr_met_SM/data/snpPCA
## create folder
mkdir results/eQTLanalysis

## Load libraries ####
library(snpStats)
library(dplyr)
library(MatrixEQTL)
library(SummarizedExperiment)
library(minfi)
library(mediation)

# Prepare data ####
## Load eQTM catalogue (http://www.mqtldb.org/download.htm)
mQTLs <- read.table("data/ARIES_mQTLs.tab", header = TRUE, as.is = TRUE)

selSNPs <- unique(mQTLs$SNP)
selCpGs <- unique(mQTLs$gene)

## Load Genotypes
plinkSNPs <- read.table("~/data/WS_HELIX/HELIX_preproc/gwas/Final_data_HRCimp_QC2/HELIX.impQC.rs.bim")
comSNPs <- intersect(as.character(plinkSNPs$V2), selSNPs)
plink <- read.plink("/home/isglobal.lan/cruiz/data/WS_HELIX/HELIX_preproc/gwas/Final_data_HRCimp_QC2/HELIX.impQC.rs",
                   select.snps = comSNPs)

## Load methylation
load("results/preprocessFiles/Methylation_GRSet.RData")
colnames(gset) <- gset$HelixID

## Select common samples
comSamps <- intersect(rownames(plink$geno), colnames(gset))
geno <- plink$geno[comSamps, ]

## Select common samples and CpGs
comCpGs <- intersect(selCpGs, rownames(gset))
gsetF <- gset[comCpGs, comSamps]

## Get covariables
pheno <- colData(gsetF)[, c("cohort", "e3_sex", "age_sample_years", "NK_6", "Bcell_6", 
                           "CD4T_6", "CD8T_6", "Gran_6", "Mono_6")]

pcs <- read.table("~/data/WS_HELIX/HELIX_analyses/expr_met_SM/data/snpPCA.eigenvec", 
                  row.names = 1)
colnames(pcs) <- c("Name", paste0("PC", 1:20))
pheno$Name <- rownames(pheno)
covars <- pheno %>%
  as_tibble() %>%
  left_join(pcs, by = "Name") %>%
  select(-Name)
covars <- covars[, !colnames(covars) %in% paste0("PC", 11:20)]
covarsmod <- model.matrix(~. - 1, covars)
rownames(covarsmod) <- rownames(covars)

# data objects for Matrix eQTL engine
snps <- SlicedData$new(t(as(geno, "numeric")))
gene <- SlicedData$new(getBeta(gsetF))
cvrt <- SlicedData$new(t(covarsmod))

snpspos <- plink$map %>%
  mutate(snp = snp.name,
         chr = chromosome, 
         pos = position) %>%
  select(snp, chr, pos)
genepos <- rowRanges(gsetF) %>%
  data.frame() %>%
  mutate(geneid = names(rowRanges(gsetF)),
         chr = gsub("chr", "", seqnames), 
         s1 = start,
         s2 = end) %>%
  select(geneid, chr, s1, s2)

# Slice data in blocks of 500 variables
snps$ResliceCombined(500)
gene$ResliceCombined(500)

# Output file name
output_file_name_cis = "results/eQTLanalysis/eQTM.cis.results.txt";
output_file_name_tra = "results/eQTLanalysis/eQTM.trans.results.txt";

# Only associations significant at this level will be saved
pvOutputThreshold_cis = 1e-7
pvOutputThreshold_tra = 1e-7

# Error covariance matrix
# Set to numeric() for identity.
errorCovariance = numeric();

# Distance for local gene-SNP pairs
cisDist <- 1e6;

# Linear model to use, modelANOVA, modelLINEAR, or modelLINEAR_CROSS
useModel <- modelLINEAR; # modelANOVA, modelLINEAR, or modelLINEAR_CROSS

## Run the analysis ####
me = Matrix_eQTL_main(
  snps = snps,
  gene = gene,
  cvrt = cvrt,
  output_file_name     = output_file_name_tra,
  pvOutputThreshold     = pvOutputThreshold_tra,
  useModel = useModel,
  errorCovariance = errorCovariance,
  verbose = TRUE,
  output_file_name.cis = output_file_name_cis,
  pvOutputThreshold.cis = pvOutputThreshold_cis,
  snpspos = snpspos,
  genepos = genepos,
  cisDist = cisDist,
  pvalue.hist = "qqplot",
  min.pv.by.genesnp = FALSE,
  noFDRsaveMemory = FALSE)

comCisQTL <- me$cis$eqtls %>%
  mutate(SNP = snps) %>%
  semi_join(mQTLs, by = c("SNP", "gene"))

comTransQTL <- me$trans$eqtls %>%
  mutate(SNP = snps) %>%
  semi_join(mQTLs, by = c("SNP", "gene"))
save(comCisQTL, comTransQTL, file = "results/eQTLanalysis/comQTLs.Rdata")

## Run analysis using ARIES scripts ####
mkdir results/ARIES

## Create files with selected probes
mQTLs <- read.table("data/ARIES_mQTLs.tab", header = TRUE, as.is = TRUE)

selSNPs <- unique(mQTLs$SNP)
selCpGs <- unique(mQTLs$gene)

write.table(selSNPs, file = "results/ARIES/selSNPs.tab", col.names = FALSE,
            row.names = FALSE, quote = FALSE)


## Create matrix with methylation
load("results/preprocessFiles/Methylation_GRSet.RData")
colnames(gset) <- gset$HelixID
M <- getBeta(gset)

M <- M[rownames(M) %in% selCpGs, ]
save(M, file = "results/ARIES/methy.Rdata")

## Create covars
covars <- colData(gset)[, c("cohort", "e3_sex", "age_sample_years", "NK_6", "Bcell_6", 
                            "CD4T_6", "CD8T_6", "Gran_6", "Mono_6")]
colnames(covars)[1:2] <- c("Batch", "Sex")
write.table(covars, file = "results/ARIES/covars.tab", quote = FALSE,
            sep = "\t")


ids <- gset$HelixID
write.table(cbind(ids, ids), file = "F7", col.names = FALSE,
            row.names = FALSE, quote = FALSE)


## Outside python
## Modify script to adhere folders
python ~/software/ariesmqtl/run.py --geno ~/data/WS_HELIX/HELIX_preproc/gwas/Final_data_HRCimp_QC2/HELIX.impQC.rs \
--meth results/ARIES/methy.Rdata --covars results/ARIES/covars.tab \
--fo results/ARIES/mqtls.txt --keep-snps results/ARIES/selSNPs.tab \
--tp F7

Rscript ./preprocessing.R --meth results/ARIES/methy.Rdata --covars results/ARIES/covars.tab

## Run association between expression and mQTLs ####

## Get list TCs and SNPs
load("results/MethComBatExpResidualsNoCellAdj/allres_simP_cpgs.Rdata")
load("results/eQTLanalysis/comQTLs.Rdata")
selSNPs <- unique(c(as.character(comCisQTL$SNP), as.character(comTransQTL$SNP)))
selTCs <- rbind(comCisQTL, comTransQTL) %>%
  mutate(CpG = gene) %>%
  left_join(df, by = "CpG") %>%
  filter(sigPair) %>%
  pull(TC) %>%
  unique()
  
  
## Load Genotypes
plinkSNPs <- read.table("~/data/WS_HELIX/HELIX_preproc/gwas/Final_data_HRCimp_QC2/HELIX.impQC.rs.bim")
comSNPs <- intersect(as.character(plinkSNPs$V2), selSNPs)
plink <- read.plink("/home/isglobal.lan/cruiz/data/WS_HELIX/HELIX_preproc/gwas/Final_data_HRCimp_QC2/HELIX.impQC.rs",
                    select.snps = comSNPs)

## Load Gene expression
load("results/preprocessFiles/Expression_SE_residuals.RData")
colnames(se) <- se$HelixID

## Select common samples
comSamps <- intersect(rownames(plink$geno), colnames(se))
geno <- plink$geno[comSamps, ]

## Select common samples and TCs
seF <- se[selTCs, comSamps]

## Get covariables
pheno <- colData(seF)[, c("cohort", "e3_sex", "age_sample_years", "NK_6", "Bcell_6", 
                            "CD4T_6", "CD8T_6", "Gran_6", "Mono_6")]

pcs <- read.table("~/data/WS_HELIX/HELIX_analyses/expr_met_SM/data/snpPCA.eigenvec", 
                  row.names = 1)
colnames(pcs) <- c("Name", paste0("PC", 1:20))
pheno$Name <- rownames(pheno)
covars <- pheno %>%
  as_tibble() %>%
  left_join(pcs, by = "Name") %>%
  dplyr::select(-Name)
covars <- covars[, !colnames(covars) %in% paste0("PC", 11:20)]
covarsmod <- model.matrix(~. - 1, covars)
rownames(covarsmod) <- rownames(covars)

# data objects for Matrix eQTL engine
snps <- SlicedData$new(t(as(geno, "numeric")))
gene <- SlicedData$new(assay(seF))
cvrt <- SlicedData$new(t(covarsmod))

snpspos <- plink$map %>%
  mutate(snp = snp.name,
         chr = chromosome, 
         pos = position) %>%
  select(snp, chr, pos)
genepos <- rowRanges(seF) %>%
  data.frame() %>%
  mutate(geneid = names(rowRanges(seF)),
         chr = gsub("chr", "", seqnames), 
         s1 = start,
         s2 = end) %>%
  select(geneid, chr, s1, s2)

# Slice data in blocks of 500 variables
snps$ResliceCombined(500)
gene$ResliceCombined(500)

# Output file name
output_file_name_cis = "results/eQTLanalysis/eQTL.cis.results.txt";
output_file_name_tra = "results/eQTLanalysis/eQTL.trans.results.txt";

# Only associations significant at this level will be saved
pvOutputThreshold_cis = 1e-7
pvOutputThreshold_tra = 1e-7

# Error covariance matrix
# Set to numeric() for identity.
errorCovariance = numeric();

# Distance for local gene-SNP pairs
cisDist <- 1e6;

# Linear model to use, modelANOVA, modelLINEAR, or modelLINEAR_CROSS
useModel <- modelLINEAR; # modelANOVA, modelLINEAR, or modelLINEAR_CROSS

## Run the analysis ####
gexpme = Matrix_eQTL_main(
  snps = snps,
  gene = gene,
  cvrt = cvrt,
  output_file_name = output_file_name_tra,
  pvOutputThreshold = pvOutputThreshold_tra,
  useModel = useModel,
  errorCovariance = errorCovariance,
  verbose = TRUE,
  output_file_name.cis = output_file_name_cis,
  pvOutputThreshold.cis = pvOutputThreshold_cis,
  snpspos = snpspos,
  genepos = genepos,
  cisDist = cisDist,
  pvalue.hist = "qqplot",
  min.pv.by.genesnp = FALSE,
  noFDRsaveMemory = FALSE)

save(gexpme, file = "results/eQTLanalysis/eQTLs.Rdata")
```
