## SupplementalCode for "Identification of autosomal cis expression quantitative trait methylation (cis eQTMs) in children’s blood": README

These scripts contains the code used to generate the catalogue files. Files 1-6 are bash script prepared to be run in a cluster with slurm. Paths are hardcoded inside the files and should be changed by the user.

Main model eQTMs (expression ~ DNA methylation + age + sex + cohort):

1.mainModel_splitDataByChr.sh: split data by chromosome
2.mainModel_runPermutations.sh: run permutations to adjust for multiple testing
3.mainModel_geteQTMs.sh: Compute eQTMs and generate table with significant eQTMs.

Additional model eQTMs (expression ~ DNA methylation + age + sex + cohort + cell proportions):

These scripts are equivalent to the main model, just using another linear model.

4.cellAdjModel_splitDataByChr.sh
5.cellAdjModel_runPermutations.sh
6.cellAdjModel_geteQTMs.sh

7.meQTL_analysis.R: R script to run the meQTL analysis for the CpGs in eQTMs in the main model. In order to speed up the computation, we only considered SNPs that were meQTLs in ARIES, considering a p-value < 1e-7. 
README: this file.
src: Folder with R script files used by the bash scripts.
