## SupplementalCode for "Identification of autosomal cis expression quantitative trait methylation (cis eQTMs) in children’s blood": createInputFiles.R

```
###############################################################################
#' Divide GenomicRatioSet (methylation) and SummarizedExperiment (Expression) to run
#' linear models
#' @param methy Path with the GenomicRatioSet
#' @param exprs Path with the SummarizedExperiment
#' @param out_fold Path with the folder to output the results
#' @param Operation Operations to be applied to data (autosomes: select probes in 
#' autosome chromosome; sex-stratify: divide the datasets in males and females)
#' @example 
#' Rscript createInputFiles.R '--args methy="gset_sm_res1.RData" exprs="summ_exp_sm_res1.RData" out_fold="./results/model1" autosomes
###############################################################################

## Load libraries ####
library('minfi', quietly = TRUE, verbose = FALSE)
library(SummarizedExperiment, quietly = TRUE, verbose = FALSE)

arg <- commandArgs(trailingOnly = T)

## Parse arguments
for(i in 1:3){
  eval(parse(text=arg[[i]]))
}


## Load methylation ####
load(methy) # gset

## Load Expression ####
load(exprs) # se

# Check data consistency ####
stopifnot(all(colnames(gset) == colnames(se)))

print("Data loaded")

## Remove probes in sexual chromosomes
if ("autosomes" %in% arg){
  gset <- gset[!seqnames(rowRanges(gset)) %in% c("chrX", "chrY"), ]
  se <- se[!seqnames(rowRanges(se)) %in% c("chrX", "chrY"), ]
}

## Stratify individuals by sex
if ("sex-stratify" %in% arg){
  gsetall <- gset
  gset <- gsetall[, gsetall$e3_sex == "male"]
  save(gset, file = paste0(out_fold, "/methyInputMale.Rdata"))

  gset <- gsetall[, gsetall$e3_sex == "female"]
  save(gset, file = paste0(out_fold, "/methyInputFemale.Rdata"))
  
  seall <- se
  se <- seall[, seall$e3_sex == "male"]  
  save(se, file = paste0(out_fold, "/gexpInputMale.Rdata"))

  se <- seall[, seall$e3_sex == "female"]
  save(se, file = paste0(out_fold, "/gexpInputFemale.Rdata"))
} else {
  save(gset, file = paste0(out_fold, "/methyInput.Rdata"))
  save(se, file = paste0(out_fold, "/gexpInput.Rdata"))
}
```
