## SupplementalCode for "Identification of autosomal cis expression quantitative trait methylation (cis eQTMs) in children’s blood": divide_data.R

```
###############################################################################
#' Divide GenomicRatioSet (methylation) and SummarizedExperiment (Expression) to run
#' linear models
#' @param methy Path with the GenomicRatioSet
#' @param exprs Path with the SummarizedExperiment
#' @param out_fold Path with the folder to output the results
#' @example 
#' Rscript divide_datasets.R '--args methy="gset_sm_res1.RData" exprs="summ_exp_sm_res1.RData" out_fold="model1"
###############################################################################

## Load libraries ####
library('minfi', quietly = TRUE, verbose = FALSE)
library(SummarizedExperiment, quietly = TRUE, verbose = FALSE)

arg <- commandArgs(trailingOnly = T)
## Parse arguments
for(i in 1:length(arg)){
  eval(parse(text=arg[[i]]))
}

## Load methylation ####
load(methy) # gset

## Load Expression ####
load(exprs) # se

# Check data consistency ####
stopifnot(colnames(gset) == colnames(se))
print("Data loaded")

## Get data matrices
ma <- assays(gset)$Beta # methylation assay
ea <- assays(se)$exprs # expression assay

## Get probes ranges
mr <- rowRanges(gset)[,0] # methylation ranges removing metadata
er <- rowRanges(se)[,0] # expression ranges removing metadata

# Automatic subsetting + saving
count <- 0
chrs <- unique(as.character(seqnames(rowRanges(gset))))
for (chr in chrs) {
  mrsub <- mr[seqnames(mr) == chr]
  ersub <- er[seqnames(er) == chr]
  fo <- findOverlaps(mrsub + 5e5, ersub)
  
  cpgnames <- names(mrsub)
  tcnames <- names(ersub)
  cpg <- cpgnames[from(fo)]
  tc <- tcnames[to(fo)]
  
  overlaps <- cbind(cpg, tc)
  masub <- ma[cpgnames,]
  easub <- ea[tcnames,]
  save(overlaps, file = paste0(out_fold, "/", 'overlaps', chr, '.RData'))
  save(masub, file = paste0(out_fold, "/", 'masub', chr, '.RData'))
  save(easub, file = paste0(out_fold, "/", 'easub', chr, '.RData'))
  print(paste(chr, nrow(overlaps), sep = ': '))
  count <- count + nrow(overlaps)
}
print(count)

## Save phenotypes
pheno <- colData(gset)
save(pheno, file = paste0(out_fold, "/pheno.RData"))

########################### Test with CHROMOSOME 22
# mrsub <- mr[seqnames(mr) == 'chr22'] # mr subset
# ersub <- er[seqnames(er) == 'chr22'] # er subset
# fo <- findOverlaps(mrsub + 5e5, ersub) # overlaps
# 
# cpgnames <- names(mrsub) # cpg names of the subset
# tcnames <- names(ersub) # tc names of the subset
# cpg <- cpgnames[from(fo)] # overlapped cpgs
# tc <- tcnames[to(fo)] # overlapped tcs
# 
# overlaps <- cbind(cpg, tc) # 308074
# masub <- ma[cpgnames,] # [1] 6831  832
# easub <- ea[tcnames,] # [1] 1382  832
###################################################
```
