## SupplementalCode for "Identification of autosomal cis expression quantitative trait methylation (cis eQTMs) in children’s blood": getBetaDistributionsCpGs.R

```
###############################################################################
# Reestimate pvalues using simulations
#' @param methy Path with the GenomicRatioSet
#' @param exprs Path with the SummarizedExperiment
#' @param out_fold Path with the folder to output the results
#' @example 
#' Rscript getBetaDistributionsCpGs.R '--args folder="results/MethComBatExpResidualsCellAdj" type="autosome"
###############################################################################
library(parallel)

arg <- commandArgs(trailingOnly = T)
## Parse arguments
for(i in 1:length(arg)){
  eval(parse(text=arg[[i]]))
}

load("results/preprocessFiles/allOverlaps.Rdata")
overDF$pair <- paste0(overDF$CpG, overDF$TC)

if (type == "autosome"){
  chr <- 1:22
} else if (type == "female") {
  chr <- c(1:22, "X")
} else if (type == "male"){
  chr <- c(1:22, "X", "Y")
}

## Load original results
getBetaDistributionsCpGs <- function(folder, chr = 1:22){
  ## Load list of folders
  simfolders <- dir(folder, pattern = "sim[0-9]", full.names = TRUE)
  
  res <- lapply(chr, function(chrom){
    
    print(chrom)
    resList <- lapply(simfolders, function(simFold) {
      print(paste0(simFold, "/outputchr", chrom, ".txt.gz"))
      df <- read.table(gzfile(paste0(simFold, "/outputchr", chrom, ".txt.gz")),
                 as.is = TRUE)
      ### Remove bad overlaps (solve bug)
      df[paste0(df$V1, df$V2) %in% overDF$pair, ]
    })
    CpGs <- unique(resList[[1]]$V1)
    
    distrs <- mclapply(CpGs, function(cpg){
      
      ## Get pvalues per CpG
      resCpG <- lapply(resList, function(x) {
        df <- x[x$V1 == cpg, ]
        pvals <- df$V5
      })
      
      perms <- Reduce(cbind, resCpG)
      pmin <-  matrixStats::colMins(perms, na.rm=TRUE)
      
      llhd2 <- function(x,p) {
        ans <- -sum(log(dbeta(x,p[1],p[2])))
        ans
      }
      
      pIni <- c(1,100)
      
      param <- nlm(llhd2, x=pmin, p=pIni)$estimate
      
      param
    }, mc.cores = 15)
    names(distrs) <- CpGs
    distrs
    
  })
}
CpGsDistrs <- getBetaDistributionsCpGs(folder, chr)
save(CpGsDistrs, file = paste0(folder, "/CpGsDistr.Rdata"))
```
