## SupplementalCode for "Identification of autosomal cis expression quantitative trait methylation (cis eQTMs) in children’s blood": getSignificantPairs.R

```
###############################################################################
# Reestimate pvalues using simulations
#' @param resFolder Folder with the analyses results
#' @param distribution File with the beta distributions
#' @param base Significance based on cpgs or genes?
#' @example 
#' Rscript getSignificantPairs.R '--args resFolder="results/MethComBatExpResidualsCellAdj" distribution="results/MethComBatExpResidualsCellAdj/CpGsDistr.Rdata" base="cpgs"
###############################################################################

library(pbapply)
pboptions(type="timer")


arg <- commandArgs(trailingOnly = T)
## Parse arguments
for(i in 1:length(arg)){
  eval(parse(text=arg[[i]]))
}

load(distribution)
if (base == "cpgs"){
  distr <- unlist(CpGsDistrs, recursive = FALSE)
} else if (base == "genes"){
  distr <- unlist(GeneDistrs, recursive = FALSE)
} else {
  stop("Base should be cpgs or genes")
}

load("results/preprocessFiles/allOverlaps.Rdata")
overDF$pair <- paste0(overDF$CpG, overDF$TC)

## Load results
files <- dir(resFolder, pattern = "outputchr", full.names = TRUE)
resList <- lapply(files, function(x) read.table(gzfile(x), as.is = TRUE))
df <- Reduce(rbind, resList)
colnames(df) <- c("CpG", "TC", "FC", "SD", "p.value", "CI0.05", "CI0.95")
df <- df[paste0(df$CpG, df$TC) %in% overDF$pair, ]

## Compute p-value per feature
if (base == "cpgs"){
  feats <- unique(df$CpG)
} else if (base == "genes"){
  feats <- unique(df$TC)
}
feats <- intersect(feats, names(distr))

featPvals <- function(feat, df, distr, base){
  
  d <- distr[[feat]]
  col <- ifelse(base == "cpgs", "CpG", "TC")
  df <- df[df[, col, drop = TRUE] == feat, , drop = FALSE]
  
  pbeta(min(df$p.value), d[1], d[2], lower.tail = TRUE)
}

featsPvals <- pbsapply(feats, featPvals, df = df, distr = distr, base = base, 
                     cl = 25)
featStatsDF <- data.frame(feat = feats, p.val = featsPvals, 
                          p.val.adj = p.adjust(featsPvals, "BH"),
                          stringsAsFactors = FALSE)
sigFeats <- subset(featStatsDF, p.val.adj < 0.05)$feat
thresP <- max(featStatsDF[sigFeats, "p.val"])

isSig <- function(row, df, sigFeats, distr, thres, base){
  

col <- ifelse(base == "cpgs", "CpG", "TC")
  feat <- df[row, col]
  d <- distr[[feat]]
  
  if (!feat %in% sigFeats){
    return(FALSE)
  }
  pval <- df[row, "p.value"]
  pbeta(pval, d[1], d[2], lower.tail = TRUE) <= thres
}

df$sigPair <- pbsapply(seq_len(nrow(df)), isSig, df = df, distr = distr, base = base, 
                sigFeats = sigFeats, thres = thresP, cl = 25)

save(df, featStatsDF, sigFeats, file = paste0(resFolder, "/allres_simP_", base, ".Rdata"))
```
