## SupplementalCode for "Identification of autosomal cis expression quantitative trait methylation (cis eQTMs) in children’s blood": runLinearModelSubset.R

```
###############################################################################
#' Run linear model between methylation and expression in HELIX
#' @param data_fold Path with the methylation, expression and overlaps data
#' @param chr Chromosome used to compute the associations
#' @param model Model used in the linear model (see Define linear models section)
#' @param out_fold Path with the folder to output the results
#' @param sim Number of simulation
#' @example 
#' Rscript runLinearModelSubset '--args data_fold="model1" chr="1" out_fold="results/model1" model="cell"' 
###############################################################################
library("parallel", verbose = FALSE)
library("S4Vectors", verbose = FALSE)

arg <- commandArgs(trailingOnly = T)

print(arg)

## Parse arguments
for(i in 1:4){
  eval(parse(text=arg[[i]]))
}

# Load data ####
a <- paste0(data_fold, "/", 'easub', chr, '.RData') # object: easub
b <- paste0(data_fold, "/", 'masub', chr, '.RData') # object: masub
c <- paste0(data_fold, "/", 'overlaps', chr, '.RData', sep = '') # object: overlaps
load(a)
load(b)
load(c)
load(paste0(data_fold, "/", "pheno.RData"))

# Check data consistency ####
stopifnot(colnames(masub) == colnames(easub), 
          rownames(pheno) == colnames(masub))
          
# Define Linear models ####
models <- c(cell = easub[tc,] ~ masub[cpg,] + pheno$cohort + pheno$e3_sex +
              pheno$age_sample_years  + pheno$NK_6 + pheno$Bcell_6 +
              pheno$CD4T_6 + pheno$CD8T_6 + pheno$Gran_6 + pheno$Mono_6, 
            nocell = easub[tc,] ~ masub[cpg,] + pheno$cohort + pheno$e3_sex +
              pheno$age_sample_years,
            cellStrat = easub[tc,] ~ masub[cpg,] + pheno$cohort + 
              pheno$age_sample_years  + pheno$NK_6 + pheno$Bcell_6 +
              pheno$CD4T_6 + pheno$CD8T_6 + pheno$Gran_6 + pheno$Mono_6, 
            nocellStrat = easub[tc,] ~ masub[cpg,] + pheno$cohort + 
              pheno$age_sample_years)
stopifnot(model %in% names(models))
model <- models[[model]]

# Sample Expression data ####
## Only in simulations
if (length(arg) == 5){
  sim <- as.numeric(eval(parse(text = arg[[5]])))
  set.seed(sim)
  easub <- easub[, sample(colnames(easub))]
  
  ### Add sim name to out_fold
  out_fold <- paste0(out_fold, "/sim", sim)
}

print(out_fold)

resCorr <- function(x) {
  cpg <- overlaps[x, 1]
  tc <- overlaps[x, 2]
  fit <- lm(model, data = environment())
  return(c(cpg,
           tc,
           summary(fit)$coef[2,1],
           summary(fit)$coef[2,2],
           summary(fit)$coef[2,4],
           confint(fit)[2,1],
           confint(fit)[2,2]))
}

len <- 1:nrow(overlaps)
results <- mclapply(len, resCorr, mc.cores = 16)
output <- matrix(unlist(results), ncol = 7, byrow = TRUE)
filename <- paste(out_fold, '/output', chr, '.txt', sep = '')
write.table(output, file = filename, col.names = F, row.names = F,
            quote = F)
```
