## SupplementalCode for "Identification of autosomal cis expression quantitative trait methylation (cis eQTMs) in children’s blood": createWebFiles.R

```
#'##############################################################################
#' Create files for HELIX website
#' - Main model
#'   - All pairs
#'   - eQTMs
#'   - SNPs + CpGs + TCs (eQTMs)
#' - Cell adjusted model
#'   - All pairs
#'   - eQTMs
#'##############################################################################

## Load libraries ####
library(dplyr)
library(tidyr)
library(BiocGenerics)

## Load datasets ####
load("results/preprocessFiles/allOverlaps.Rdata")
load("results/preprocessFiles/methyAnnotation.Rdata")
load("results/preprocessFiles/gexpAnnotation.Rdata")
load("results/eQTLanalysis/eQTLs.Rdata")
load("results/eQTLanalysis/comQTLs.Rdata")
mQTLs <- read.table("data/ARIES_mQTLs.tab", header = TRUE, as.is = TRUE)
mQTLsH2 <- read.table("results/ARIES/mqtls.txt", header = TRUE, as.is = TRUE)

## Change name when loading results
load("results/MethComBatExpResidualsNoCellAdj/allres_simP_cpgs.Rdata")
modU <- as_tibble(df)
featsU <- featStatsDF

load("results/MethComBatExpResidualsCellAdj/allres_simP_cpgs.Rdata")
modC <- as_tibble(df)
featsC <- featStatsDF


## Select pairs detected in previous analysis from ARIES pipeline
## Merge with ARIES original data
ARIESannot <- read.table("data/ariesmqtlsnps.bim", as.is = TRUE)
colnames(ARIESannot) <- c("chr", "SNP", "cm", "pos", "Ref", "Alt")

HELIXannot <- read.table("~/data/WS_HELIX/HELIX_preproc/gwas/Final_data_HRCimp_QC2/HELIX.impQC.rs.bim", as.is = TRUE)
colnames(HELIXannot) <- c("chr", "SNP", "cm", "pos", "Ref", "Alt")

### Modify methylation annotation
methyAnnotGood <- methyAnnot %>%
  as_tibble() %>%
  mutate(CpG = Name,
         CpG_chr = chr,
         CpG_pos = pos,
         CpG_gene = sapply(UCSC_RefGene_Name, function(x) paste(unique(x), collapse = "/")),
         CpG_range = meth_range) %>%
  select(starts_with("CpG"), -CpG_maf)


### Modify Expression annotation
gexpAnnotGood <- expAnnot %>%
  as_tibble() %>%
  mutate(TC_gene_start = start,
         TC_gene_end = stop,
         TC_gene_TSS = TSS_Affy,
         TC_gene = sapply(GeneSymbol_Affy, function(x) paste(unique(x), collapse = "/")),
         TC_call_rate = CallRate) %>%
  select(starts_with("TC"), -TC_size)


## Main model tables ####
### Prepare CpGs p-vals
mainFeats <- featsU %>%
  mutate(CpG = feat, 
         CpG_pVal = p.val) %>% 
  select(starts_with("CpG"))


### Add required columns
mainAll <- modU %>%
  select(-starts_with("CI")) %>%
  mutate(log2FC = FC/10, SE = SD) %>%
  select(CpG, TC, log2FC, SE, p.value, sigPair) %>%
  left_join(methyAnnotGood, by = "CpG") %>%
  left_join(mainFeats, by = "CpG") %>%
  left_join(gexpAnnotGood, by = "TC")

write.table(mainAll, file = "HELIX_blood_eQTM_WebCat/eQTM_autosome_unadj.cells.txt", 
            col.names = TRUE, sep = "\t",
            quote = FALSE, row.names = FALSE)

maineQTM <- subset(mainAll, sigPair == TRUE)
write.table(maineQTM, file = "HELIX_blood_eQTM_WebCat/eQTM_autosome_unadj.cells_SIG.txt", 
            col.names = TRUE,  sep = "\t",
            quote = FALSE, row.names = FALSE)


## Cell adjusted model tables ####
### Prepare CpGs p-vals
cellFeats <- featsC %>%
  mutate(CpG = feat, 
         CpG_pVal = p.val) %>% 
  select(starts_with("CpG"))


### Add required columns
cellAll <- modC %>%
  select(-starts_with("CI")) %>%
  mutate(log2FC = FC/10, SE = SD) %>%
  select(CpG, TC, log2FC, SE, p.value, sigPair) %>%
  left_join(methyAnnotGood, by = "CpG") %>%
  left_join(cellFeats, by = "CpG") %>%
  left_join(gexpAnnotGood, by = "TC")

write.table(cellAll, file = "HELIX_blood_eQTM_WebCat/eQTM_autosome_adj.cells.txt", 
            col.names = TRUE, sep = "\t",
            quote = FALSE, row.names = FALSE)

celleQTM <- subset(cellAll, sigPair == TRUE)
write.table(celleQTM, file = "HELIX_blood_eQTM_WebCat/eQTM_autosome_adj.cells_SIG.txt", 
            col.names = TRUE, sep = "\t",
            quote = FALSE, row.names = FALSE)


## SNP-CpG-TC table
comMQTLs <- mQTLsH2 %>%
  mutate(gene = CpG) %>%
  dplyr::select(SNP, gene, A1, A2, freq, b, se, p, N, r2) %>%
  semi_join(rbind(comCisQTL, comTransQTL), by = c("SNP", "gene")) %>%
  inner_join(left_join(mQTLs, dplyr::select(ARIESannot, SNP, Ref, Alt), by = "SNP"), 
             by = c("SNP", "gene")) %>%
  as_tibble()

comMQTLs.f <- comMQTLs %>%
  filter(!is.na(Ref)) %>%
  filter(!(sign(b) != sign(beta) & A1 == Ref)) %>%
  filter(!(sign(b) == sign(beta) & A1 == Alt))

eQTL <- rbind(gexpme$cis$eqtls, gexpme$trans$eqtls) %>%
  mutate(TC = gene) %>%
  dplyr::select(-gene)

sigDf <- filter(modU, sigPair)


mergedDf <- rbind(comCisQTL, comTransQTL) %>%
  mutate(CpG = gene) %>%
  left_join(select(comMQTLs.f, SNP, gene, A1, A2, freq), by = c("SNP", "gene")) %>%
  inner_join(sigDf, by = "CpG") %>%
  inner_join(eQTL, by = c("snps", "TC")) %>%
  dplyr::select(-sigPair) %>%
  filter(sign(beta.x)*sign(FC) == sign(beta.y))

snpTab <- mergedDf %>%
  mutate(SNP_CpG_beta = beta.x/10,
         SNP_CpG_p.value = pvalue.x,
         SNP_TC_log2FC = beta.y,
         SNP_TC_p.value = pvalue.y,
         CpG_TC_log2FC = FC/10,
         CpG_TC_p.value = p.value,
         EA = A1,
         NEA = A2,
         EF = freq) %>%
  select(SNP, CpG, TC, EA, NEA, EF, SNP_CpG_beta, SNP_CpG_p.value,
         SNP_TC_log2FC, SNP_TC_p.value, CpG_TC_log2FC,
         CpG_TC_p.value)
write.table(snpTab, file = "HELIX_blood_eQTM_WebCat/eQTM_SNPs_autosome_unadj.cells.txt", 
            col.names = TRUE, sep = "\t",
            quote = FALSE, row.names = FALSE)
```
