## SupplementalCode for "Identification of autosomal cis expression quantitative trait methylation (cis eQTMs) in children’s blood": eQTM_catalogue.R

```
#'##############################################################################
#' Generate eQTM catalogue figures and tables
#' This script contains the code for
#'  - Volcano plot
#'  - QC checks
#'  - Get number significants pairs and make subgroups
#'  - Make distance plots
#'  - Compare Illumina annotation vs eQTMs
#'  - Compare Non-cell adjusted  (reference) vs cell adjusted models
#'  - Compare with other eQTMs studies
#'##############################################################################

## Load libraries ####
library(dplyr)
library(ggplot2)
library(cowplot)
library(tidyr)
library(S4Vectors)
library(MultiDataSet)
library(ggforce)
library(GenomicRanges)
library(BSgenome.Hsapiens.UCSC.hg19)
library(limma)
library(eulerr)

## Load datasets ####
load("results/preprocessFiles/allOverlaps.Rdata")
load("results/preprocessFiles/methyAnnotation.Rdata")
load("results/preprocessFiles/gexpAnnotation.Rdata")

## Change name when loading results
load("results/MethComBatExpResidualsCellAdj/allres_simP_cpgs.Rdata")
modC <- df
featsC <- featStatsDF

## Create useful vars
### data.frame only with significant pairs
sigDf <- modC %>%
  as_tibble() %>%
  filter(sigPair)

codingTCs <- subset(expAnnot, Coding == "coding")$transcript_cluster_id
sigTCs <- unique(sigDf$TC)

### data.frame with results and overlaps info
modC_comp <- as_tibble(inner_join(modC, overDF, by = c("CpG", "TC")))

### Modify methylation annotation
methyAnnot <- methyAnnot %>%
  as_tibble() %>%
  mutate(CpG = Name)

# General Overview ####
## Statistics ####
### Significant pairs
nrow(sigDf)

### Significant TC
length(unique(sigDf$TC))

### Significant coding TC
sum(unique(sigDf$TC) %in% codingTCs)

### Significant CpGs
length(unique(sigDf$CpG))

### Proportion negative pairs
sum(sigDf$FC < 0)

mean(sigDf$FC < 0)*100

## Statistics with FDR
### Significant pairs
sigDF_FDR <- modC %>%
  as_tibble() %>%
  mutate(FDR = p.adjust(p.value, method = "BH")) %>%
  filter(FDR < 0.05)
nrow(sigDF_FDR)

### Significant TC
length(unique(sigDF_FDR$TC))

### Significant CpGs
length(unique(sigDF_FDR$CpG))


## Distribution CpGs/TC
CpG_plot <- sigDf %>%
  group_by(CpG) %>%
  summarize(n = n()) %>%
  ggplot(aes(x = n)) + geom_histogram(binwidth = 1) +
  scale_x_continuous("eGenes per CpG", limits = c(0, 50)) +
  scale_y_continuous("Number of CpGs")  +
  geom_vline(aes(xintercept = median(n))) +
  theme_bw() +
  theme(plot.title = element_text(hjust = 0.5)) +
  ggtitle("CpGs pairing distribution in cis eQTMs")

TC_plot <- sigDf %>%
  group_by(TC) %>%
  summarize(n = n()) %>%
  ggplot(aes(x = n)) + geom_histogram(binwidth = 1) +
  scale_x_continuous("CpGs per eGene", limits = c(0, 50)) +
  scale_y_continuous("Number of eGenes")  +
  geom_vline(aes(xintercept = median(n))) +
  theme_bw() +
  theme(plot.title = element_text(hjust = 0.5)) +
  ggtitle("eGenes pairing distribution in cis eQTMs")


png("paper/eQTMs_CpGs_TC_distr.png", width = 2000, height = 1500, res = 300)
plot_grid(CpG_plot, TC_plot, labels = "AUTO", nrow = 2)
dev.off()

sigDf %>%
  group_by(TC) %>%
  summarize(n = n()) %>%
  `$`("n") %>%
  summary()

sigDf %>%
  group_by(CpG) %>%
  summarize(n = n()) %>%
  `$`("n") %>%
  summary()


## QCs ####
### CpGs variability
featsC_var <- methyAnnot %>%
  as_tibble() %>%
  mutate(feat = Row.names) %>%
  dplyr::select(feat, meth_range, variability) %>%
  right_join(featsC, by = "feat") %>%
  mutate(sig = ifelse(p.val.adj < 0.05, "significant", "random"))
table(featsC_var$variability, featsC_var$sig)
chisq.test(table(featsC_var$variability, featsC_var$sig))

t <- table(featsC_var$variability, featsC_var$sig)
t[1]/t[2]/t[3]*t[4]

png("paper/CpGVar_eQTMs.png", width = 2000, height = 1000, res = 300)
featsC_var %>%
  mutate(sigVar = ifelse(sig == "random", "non eCpGs", "eCpGs")) %>%
  ggplot(aes(x = sigVar, y = meth_range, fill = sigVar)) + 
  geom_boxplot() +
  theme_bw() +
  scale_x_discrete(name = "CpG Type") +
  theme(legend.position = "none") +
  scale_y_continuous(name = "Methylation range") +
  scale_fill_manual(values = c("#999999", "#FFFFFF"))
dev.off()

### CpGs Reliability
featsC_rel <- methyAnnot %>%
  as_tibble() %>%
  mutate(feat = Row.names) %>%
  dplyr::select(feat, Reliability) %>%
  mutate(rel = ifelse(Reliability > 0.4, "Reliable", "Unreliable")) %>%
  right_join(featsC, by = "feat") %>%
  mutate(sig = ifelse(p.val.adj < 0.05, "significant", "random"))
t <- table(featsC_rel$rel, featsC_rel$sig)
t
chisq.test(t)
t[2]/t[1]/t[4]*t[3]
wilcox.test(Reliability ~ sig, featsC_rel)

png("paper/CpGReliability_eQTMs.png", width = 2000, height = 1000, res = 300)
featsC_rel %>%
  mutate(sigVar = ifelse(sig == "random", "non eCpGs", "eCpGs")) %>%
  ggplot(aes(x = sigVar, y = Reliability, fill = sigVar)) + 
  geom_boxplot() +
  theme_bw() +
  scale_x_discrete(name = "CpG Type") +
  theme(legend.position = "none") +
  scale_y_continuous(name = "Probe reliability") +
  scale_fill_manual(values = c("#999999", "#FFFFFF"))
dev.off()


### TC call rate
int_TC <- expAnnot %>%
  dplyr::select(probeset_id, Expressed, CallRate) %>%
  mutate(sig = ifelse(probeset_id %in% sigTCs, "eGenes", "non eGenes"))
table(int_TC$CallRate < 90, int_TC$sig)

chisq.test(table(int_TC$CallRate > 90, int_TC$sig))
# X-squared = 1149, df = 1, p-value < 2.2e-16
t <- table(int_TC$CallRate < 90, int_TC$sig)
t[1]/t[2]/t[3]*t[4]

png("paper/CallRate_eQTMs.png", width = 2000, height = 1000, res = 300)
int_TC %>%
  ggplot(aes(x = sig, y = CallRate, fill = sig)) + 
  geom_boxplot() +
  theme_bw() +
  scale_x_discrete(name = "Gene type") +
  theme(legend.position = "none") +
  scale_y_continuous(name = "Gene call rate") +
  scale_fill_manual(values = c("#999999", "#FFFFFF"))
dev.off()

### Classify CpG in groups ####
CpGsSum <- modC %>%
  group_by(CpG) %>%
  summarise(Type = ifelse(sum(sigPair) == 0, "Non-significant",
                          ifelse(sum(sigPair) == 1, "Mono", "Multi")),
            Direction = ifelse(sum(sigPair) == 0, "Non-significant",
                               ifelse(all(FC[sigPair] > 0), "Positive", 
                                      ifelse(all(FC[sigPair] < 0), "Inverse", "Both")))) %>%
  mutate(Combined = ifelse(Type == "Non-significant", 
                           "Non-significant", 
                           paste(Type, Direction, sep = "_")))
t <- table(CpGsSum$Type, CpGsSum$Direction)
t <- t[c("Mono", "Multi"), c("Inverse", "Positive", "Both")]

chisq.test(t[, -3])
# X-squared = 8.2039, df = 1, p-value = 0.00418
t[2]/t[1]/t[4]*t[3]
addmargins(prop.table(t))

sink("paper/CpGs_type.txt")
addmargins(t)
sink()

# Distance plots ####
# Distance distribution
## Signif vs no-signif

ks.test(filter(modC_comp, sigPair) %>% pull(., Distance),
        filter(modC_comp, !sigPair) %>% pull(., Distance))
# Two-sample Kolmogorov-Smirnov test
# 
# data:  filter(modU_comp, sigPair) %>% pull(., Distance) and filter(modU_comp, !sigPair) %>% pull(., Distance)
# D = 0.15302, p-value < 2.2e-16
# alternative hypothesis: two-sided

wilcox.test(filter(modC_comp, sigPair) %>% pull(., Distance))
# Wilcoxon rank sum test with continuity correction
# 
# data:  filter(modU_comp, sigPair) %>% pull(., Distance) and filter(modU_comp, !sigPair) %>% pull(., Distance)
# V = 932277653, p-value < 2.2e-16
# alternative hypothesis: true location shift is not equal to 0


modC_comp %>% 
  group_by(sigPair) %>%
  summarize(m = median(Distance), 
            l = quantile(Distance, 0.25), 
            h = quantile(Distance, 0.75))

## Distance per direction
modC_comp %>% 
  filter(sigPair) %>%
  mutate(Direction = ifelse(FC > 0, "Positive", "Inverse")) %>%
  group_by(Direction) %>%
  summarize(m = median(Distance), 
            l = quantile(Distance, 0.25), 
            h = quantile(Distance, 0.75))


## Distance per direction + eQTM 
dist_plot <- modC_comp %>%
  mutate(Direction = ifelse(!sigPair, "Non eQTMs", ifelse(FC > 0, "Positive", "Inverse")),
         Direction = factor(Direction, levels = c("Inverse", "Positive", "Non eQTMs"))) %>%
  ggplot(aes(x = Distance, color = Direction)) + geom_density() + 
  theme_bw() + 
  scale_color_manual(name = "eQTM type", values = c("#E69F00", "#009E73", "#000000")) +
  scale_x_continuous(name = "CpG-Gene's TSS distance", 
                     breaks = c(-5e5, -2e5, 0, 2e5, 5e5), 
                     labels = c("-500Kb", "-250Kb", "0", "250Kb", "500Kb")) +
  scale_y_continuous(name = "Density") + 
  theme(plot.title = element_text(hjust = 0.5))
png("paper/distance_distr.png", width = 2000, height = 1300, res = 300)
dist_plot 
dev.off()


# Effect size ####
summary(abs(subset(modC_comp, sigPair)$FC/10))
mean(subset(modC_comp, sigPair)$FC/10 < 0.5)

ks.test(abs(subset(sigDf, FC > 0)$FC)/10,
        abs(subset(sigDf, FC < 0)$FC)/10)

modC_comp %>%
  filter(sigPair) %>%
  mutate(Dir = ifelse(FC > 0, "Positive", "Inverse"),
         FC = FC/10) %>%
  group_by(Dir) %>%
  summarize(m = median(FC),
            l = quantile(FC, 0.25),
            h = quantile(FC, 0.75))


# Distance vs Effect size
dist_eff <- modC_comp %>%
  filter(sigPair) %>%
  mutate(Direction = ifelse(FC > 0, "Positive", "Inverse"),
         FCwind = ifelse(abs(FC) > quantile(abs(FC), 0.99), quantile(abs(FC), 0.99)*sign(FC), FC)) %>%
  ggplot(aes(x = Distance, y = FCwind/10, color = Direction)) + 
  geom_point(alpha = 0.15) + 
  theme_bw() + 
  scale_color_manual(name = "eQTM type", 
                     breaks = c("Inverse", "Positive"),
                     values = c("#E69F00", "#009E73")) +
  scale_x_continuous(name = "eCpG-eGene's TSS distance",
                     breaks = c(-5e5, -2e5, 0, 2e5, 5e5), 
                     labels = c("-500Kb", "-250Kb", "0", "250Kb", "500Kb")) +
  scale_y_continuous(name = "log2 FC/0.1 Methylation") + 
  theme(plot.title = element_text(hjust = 0.5))

png("paper/distance_effect.png", width = 2000, height = 1300, res = 300)
dist_eff
dev.off()

png("paper/distance_effect_panel.png", width = 2000, height = 1500, res = 300)
plot_grid(dist_plot, dist_eff, labels = "AUTO", nrow = 2)
dev.off()

png("paper/distance_effect_reviewer.png", width = 2000, height = 1300, res = 300)
modC_comp %>%
  filter(sigPair) %>%
  mutate(Direction = ifelse(FC > 0, "Positive", "Inverse"),
         Distance.dir = ifelse(Distance >= 0, +1, -1),
         Distance = log10(abs(Distance) + 0.1)*Distance.dir,
         FCabs = ifelse(abs(FC) > quantile(abs(FC), 0.99), quantile(abs(FC), 0.99), abs(FC))) %>%
  ggplot(aes(x = Distance, y = FCabs/10, color = Direction)) + 
  geom_point(alpha = 0.1) + 
  theme_bw() + 
  scale_color_manual(name = "eQTM type", 
                     breaks = c("Inverse", "Positive"),
                     values = c("#E69F00", "#009E73")) +
  scale_x_continuous(name = "CpG-TC TSS distance",
                     breaks = c(-5, -4, -3, 0, 3, 4, 5), 
                     labels = c("-100Kb", "-10Kb", "-1Kb", "0", "1Kb", "10Kb", "100Kb")) +
  scale_y_continuous(name = "log2 FC/0.1 Methylation") + 
  theme(plot.title = element_text(hjust = 0.5))
dev.off()

png("paper/distance_effect_reviewer2.png", width = 2000, height = 1300, res = 300)
modC_comp %>%
  filter(sigPair) %>%
  mutate(DistanceQuant = cut(abs(Distance) + 0.1, c(0, 10, 100, 1e3, 1e4, 1e5, 5e5)),
         FCabs = ifelse(abs(FC) > quantile(abs(FC), 0.95), quantile(abs(FC), 0.95), abs(FC))) %>%
  ggplot(aes(x = DistanceQuant, y = abs(FCabs/10))) + 
  geom_boxplot() + 
  theme_bw() + 
  scale_y_continuous(name = "log2 FC/0.1 Methylation") + 
  theme(plot.title = element_text(hjust = 0.5))
dev.off()

png("paper/distance_pvalue_reviewer.png", width = 2000, height = 1300, res = 300)
modC_comp %>%
  filter(sigPair) %>%
  mutate(Direction = ifelse(FC > 0, "Positive", "Inverse")) %>%
  ggplot(aes(x = Distance, y = -log10(p.value), color = Direction)) + 
  geom_point(alpha = 0.1) + 
  theme_bw() + 
  scale_color_manual(name = "eQTM type", 
                     breaks = c("Inverse", "Positive"),
                     values = c("#E69F00", "#009E73")) +
  scale_x_continuous(breaks = c(-5e5, -2e5, 0, 2e5, 5e5), 
                     labels = c("-500Kb", "-250Kb", "0", "250Kb", "500Kb")) +
  scale_y_continuous(name = "-log10 p-value") + 
  theme(plot.title = element_text(hjust = 0.5))
dev.off()


modC_comp %>%
  filter(sigPair) %>%
  mutate(Direction = ifelse(FC > 0, "Positive", "Inverse")) %>%
  cor.test(x = pull(., Distance), 
           y = pull(., FC),
           cot = .)

## Correlation FC vs Distance
modC_comp %>%
  filter(sigPair) %>%
  lm(abs(FC) ~ abs(Distance), .) %>%
  summary()
modC_comp %>%
  filter(sigPair) %>%
  lm(abs(FC/10) ~ log10(abs(Distance)+0.1), .) %>%
  summary()
modC_comp %>%
  filter(sigPair & Distance > 0) %>%
  lm(abs(FC) ~ log10(abs(Distance)+0.1), .) %>%
  summary()
modC_comp %>%
  filter(sigPair & Distance < 0) %>%
  lm(abs(FC) ~ log10(abs(Distance)+0.1), .) %>%
  summary()
modC_comp %>%
  filter(sigPair) %>%
  lm(abs(FC) ~ sign(Distance), .) %>%
  summary()
modC_comp %>%
  filter(sigPair) %>%
  filter(abs(Distance) < 10e3) %>%
  summarize(n = sum(abs(FC) > 20))


modC_comp %>%
  filter(sigPair) %>%
  mutate(type = ifelse(abs(Distance) < 10, "Close", 
                       ifelse(abs(Distance) < 1e3, "Cis", "Distant")),
         distQuant = cut(log10(abs(Distance) + 1), 5)) %>%
  group_by(distQuant) %>%
  summarize(m = median(abs(FC)),
            n = n())


## Illumina Annotation ####
modC_Annot <- modC %>%
  as_tibble() %>%
  dplyr::select(CpG, TC, FC, p.value, sigPair) %>%
  left_join(CpGsSum, by = "CpG") %>%
  left_join(dplyr::select(methyAnnot, CpG, UCSC_RefGene_Group, UCSC_RefGene_Name), by = "CpG") %>%
  left_join(dplyr::select(expAnnot, TC, GeneSymbol_Affy), by = "TC") %>%
  mutate(GeneAffy = strsplit(GeneSymbol_Affy, ";"))

## Create list with genes in common in Affy and Illumina Annotation
methGenes <- unique(unlist(modC_Annot$UCSC_RefGene_Name))
expGenes <- unique(unlist(modC_Annot$GeneAffy))
comGenes <- intersect(methGenes, expGenes)

## Create list with genes in common in Affy and Illumina Annotation that are
## part of an eQTM
modC_Annot_f <- modC_Annot %>%
  filter(sigPair)

methGenesf <- unique(unlist(modC_Annot_f$UCSC_RefGene_Name))
expGenesf <- unique(unlist(modC_Annot_f$GeneAffy))
comGenesf <- intersect(methGenesf, expGenesf)

modC_Annot_f %>%
  dplyr::select(CpG, UCSC_RefGene_Name) %>%
  distinct() %>%
  summarize(n = n(),
            n_g = sum(UCSC_RefGene_Name != ""),
            n_GE = sum(sapply(UCSC_RefGene_Name, function(x) any(x %in% comGenes))))


modC_Annot_f2 <- modC_Annot_f %>%
  filter(sapply(UCSC_RefGene_Name, function(x) any(x %in% comGenes))) %>%
  mutate(match = sapply(seq_len(n()), function(x) any(GeneAffy[[x]] %in% UCSC_RefGene_Name[[x]])))


table(modC_Annot_f2$match)
modC_Annot_f2 %>%
  mutate(Dir = ifelse(FC > 0, "Positive", "Inverse")) %>%
  group_by(Dir) %>%
  summarize(N = sum(match),
            P = mean(match))


annotated <- modC_Annot_f2 %>%
  group_by(CpG) %>%
  summarize(found = any(match)) %>%
  left_join(CpGsSum, by = "CpG")
mean(annotated$found)
table(annotated$found)


annotated %>%
  group_by(Combined) %>%
  summarize(N = sum(found),
            P = mean(found))


# Enrichment of pairs with matching annotation ####
modC_Annot <- modC_Annot %>%
  mutate(gene_match = sapply(seq_len(n()), function(x) any(GeneAffy[[x]] %in% UCSC_RefGene_Name[[x]])))

t <- table(modC_Annot$sigPair, modC_Annot$gene_match)
t[1]/t[2]/t[3]*t[4]
chisq.test(t)

# Enrichment by gene position based on annotation ####
## Expand CpGs and TCs to make all pairs between CpG Gene and TC
## Restrict pairs where CpG gene and TC gene match
modC_gene_pairs <- modC_Annot %>%
  filter(gene_match) %>%
  mutate(Gene_Group = strsplit(UCSC_RefGene_Group, ";")) %>%
  unnest(c(Gene_Group, UCSC_RefGene_Name)) %>%
  mutate(GeneAffy = strsplit(GeneSymbol_Affy, ";")) %>%
  unnest(GeneAffy) %>%
  dplyr::select(CpG, TC, FC, sigPair, p.value, UCSC_RefGene_Name, Gene_Group, GeneAffy) %>%
  filter(UCSC_RefGene_Name == GeneAffy) %>%
  distinct()
  

gpos <- c("TSS1500", "TSS200", "5'UTR", "1stExon", "Body", "3'UTR")
names(gpos) <- gpos
#gpos <- c(as.list(gpos), list(Comb = c("TSS1500", "5'UTR", "Body")))

ORs <- lapply(gpos, function(x) {
  t <- table(modC_gene_pairs$Gene_Group %in% x, modC_gene_pairs$sigPair)
  p.val = chisq.test(t)$p.value
  or <-  t[1]/t[2]/t[3]*t[4]
  ORl <- log(or)
  SEl <- sqrt(1/t[1, 1] + 1/t[1, 2] + 1/t[2, 1] + 1/t[2, 2])
  c(OR = or, p.val = p.val, ORm = exp(ORl - 1.96*SEl), 
    ORM = exp(ORl + 1.96*SEl))
  
  }) %>%
  Reduce(f = cbind)
colnames(ORs) <- gpos  


ORs2 <- lapply(gpos, function(x) {
  cot <- filter(modC_gene_pairs, sigPair)
  t <- table(cot$Gene_Group %in% x, sign(cot$FC))
  list(p.value = chisq.test(t)$p.value, OR = t[1]/t[2]/t[3]*t[4])
}) %>%
  Reduce(f = cbind)
colnames(ORs2) <- gpos  


ORsInv <- lapply(gpos, function(x) {
  cot <- filter(modC_gene_pairs, modC_gene_pairs$sigPair == FALSE | FC < 0)
  t <- table(cot$Gene_Group %in% x, cot$sigPair)
  p.val = chisq.test(t)$p.value
  or <-  t[1]/t[2]/t[3]*t[4]
  ORl <- log(or)
  SEl <- sqrt(1/t[1, 1] + 1/t[1, 2] + 1/t[2, 1] + 1/t[2, 2])
  c(OR = or, p.val = p.val, ORm = exp(ORl - 1.96*SEl), 
    ORM = exp(ORl + 1.96*SEl))
}) %>%
  Reduce(f = cbind)
colnames(ORsInv) <- gpos  

ORsPos <- lapply(gpos, function(x) {
  cot <- filter(modC_gene_pairs, modC_gene_pairs$sigPair == FALSE | FC > 0)
  t <- table(cot$Gene_Group %in% x, cot$sigPair)
  p.val = chisq.test(t)$p.value
  or <-  t[1]/t[2]/t[3]*t[4]
  ORl <- log(or)
  SEl <- sqrt(1/t[1, 1] + 1/t[1, 2] + 1/t[2, 1] + 1/t[2, 2])
  c(OR = or, p.val = p.val, ORm = exp(ORl - 1.96*SEl), 
    ORM = exp(ORl + 1.96*SEl))
}) %>%
  Reduce(f = cbind)
colnames(ORsPos) <- gpos  

convertORmat <- function(OR){
  OR %>%
    data.frame() %>%
    mutate(vars = c("OR", "p-value", "ORm", "ORM")) %>%
    gather(Region, value, 1:6) %>%
    mutate(Region = gsub("X", "", Region),
           Region = gsub(".", "'", Region, fixed = TRUE))
  
}

png("paper/CpGEnrichGenePosition.png", width = 3000, height = 2000, res = 300)
rbind(convertORmat(ORs) %>% mutate(type = "All"),
      convertORmat(ORsInv) %>% mutate(type = "Inverse"),
      convertORmat(ORsPos) %>% mutate(type = "Positive")) %>%
  spread(vars, value) %>%
  mutate(Region = factor(Region, levels = gpos)) %>%
  ggplot(aes(x = type, y = OR, fill = Region)) + 
  geom_bar(stat = "identity", position=position_dodge()) + 
  geom_errorbar(position=position_dodge(.9), width=.25, aes(ymin = ORm, ymax = ORM)) +
  scale_y_continuous(trans = "log2", 
                     breaks = scales::trans_breaks("log2", function(x) round(2^x, 2))) +
  geom_hline(yintercept = 1) +
  scale_x_discrete(name = "eQTM type") +
  scale_fill_discrete(name = "Gene relative position") +
  theme_bw() 
dev.off()

## Age variability (MEDALL - Xu et al) ####
fsum <- function(x){
  if ( is.factor(x))
    "eQTM"
  else {
    sum(x)
  }
}
g2 <- function(x, gpos){
  sapply(gpos, function(y) getOR2(y, df = x))
}
getOR2 <- function(col, df){
  t <- data.matrix(df[, c(col, colnames(df)[ncol(df)])])
  t[, 2] <- t[, 2] - t[, 1]
  or <- t[1, 1]/(t[1, 2])/(t[2, 1])*t[2, 2]
  p.val <- chisq.test(t)$p.value
  ORl <- log(or)
  SEl <- sqrt(1/t[1, 1] + 1/t[1, 2] + 1/t[2, 1] + 1/t[2, 2])
  c(OR = or, p.val = p.val, ORm = exp(ORl - 1.96*SEl), 
    ORM = exp(ORl + 1.96*SEl))
}


types <- c("Inverse", "Positive", "eQTM")

agedf <- read.csv2("data/DiffMethyAgeCpGs.csv", as.is = TRUE)
agedf <- agedf %>%
  as_tibble() %>%
  mutate(Dir = ifelse(as.numeric(beta8.avg) > as.numeric(beta0.avg), "Increased", "Decreased"), 
         CpG = ILMNID)

ageSum <- CpGsSum %>%
  dplyr::select(CpG, Direction) %>%
  mutate(Direction = factor(Direction, levels = c("Inverse", "Positive", "Both", "Non-significant"))) %>%
  left_join(dplyr::select(agedf, Dir, CpG), by = "CpG") %>%
  mutate(Dir = ifelse(is.na(Dir), "Constant", Dir)) %>%
  group_by(Direction, Dir) %>%
  summarize(n = n()) %>%
  spread(Dir, n) %>%
  ungroup() %>%
  rbind(filter(., Direction != "Non-significant") %>% 
          summarize_all(fsum)) %>%
  mutate(Direction = as.character(Direction),
         Direction = ifelse(is.na(Direction), "eQTM", Direction),
         Variable = Decreased + Increased,
         tot = Variable + Constant)

ageG <- c("Variable", "Decreased", "Increased")


typesAge <- lapply(types, function(t){
  rbind(filter(ageSum, Direction == t),
        filter(ageSum, Direction == "Non-significant")) %>%
    g2(ageG) %>%
    as_tibble() %>%
    mutate(par = c("OR", "p.val", "ORlow", "ORhigh")) %>%
    gather("Region", "Value", 1:3) %>%
    mutate(Type = t)
})
combAge <- Reduce(rbind, typesAge)
combAge$dataset <- "MeDALL"


ageSum_rel <- CpGsSum %>%
  dplyr::select(CpG, Direction) %>%
  mutate(Direction = factor(Direction, levels = c("Inverse", "Positive", "Both", "Non-significant"))) %>%
  left_join(dplyr::select(agedf, Dir, CpG), by = "CpG") %>%
  left_join(dplyr::select(methyAnnot, CpG, Reliability), by = "CpG") %>%
  filter(Reliability > 0.4) %>%
  mutate(Dir = ifelse(is.na(Dir), "Constant", Dir)) %>%
  group_by(Direction, Dir) %>%
  summarize(n = n()) %>%
  spread(Dir, n) %>%
  ungroup() %>%
  rbind(filter(., Direction != "Non-significant") %>% 
          summarize_all(fsum)) %>%
  mutate(Direction = as.character(Direction),
         Direction = ifelse(is.na(Direction), "eQTM", Direction),
         Variable = Decreased + Increased,
         tot = Variable + Constant)

typesAge_rel <- lapply(types, function(t){
  rbind(filter(ageSum_rel, Direction == t),
        filter(ageSum_rel, Direction == "Non-significant")) %>%
    g2(ageG) %>%
    as_tibble() %>%
    mutate(par = c("OR", "p.val", "ORlow", "ORhigh")) %>%
    gather("Region", "Value", 1:3) %>%
    mutate(Type = t)
})
combAge_rel <- Reduce(rbind, typesAge_rel)
combAge_rel$dataset <- "MeDALL"


## Age variability (Mulder) ####
epideltadf <- read.table("data/epidelta_2020-07-17.txt", as.is = TRUE)
epideltadf$CpG <- rownames(epideltadf)
epideltadf <- epideltadf %>%
  as_tibble() %>%
  mutate(Dir = ifelse(M1.change.p > 1e-7, "Constant",
                      ifelse(M1.change.estimate > 0, "Increased", "Decreased")))

age_cont_Sum <- CpGsSum %>%
  dplyr::select(CpG, Direction) %>%
  mutate(Direction = factor(Direction, levels = c("Inverse", "Positive", "Both", "Non-significant"))) %>%
  left_join(dplyr::select(epideltadf, Dir, M1.change.estimate, CpG), by = "CpG") %>%
  group_by(Direction, Dir) %>%
  summarize(n = n()) %>%
  spread(Dir, n) %>%
  ungroup() %>%
  rbind(filter(., Direction != "Non-significant") %>% 
          summarize_all(fsum)) %>%
  mutate(Direction = as.character(Direction),
         Direction = ifelse(is.na(Direction), "eQTM", Direction),
         Variable = Decreased + Increased,
         tot = Variable + Constant)

ageG <- c("Variable", "Decreased", "Increased")


types_cont_Age <- lapply(types, function(t){
  rbind(filter(age_cont_Sum, Direction == t),
        filter(age_cont_Sum, Direction == "Non-significant")) %>%
    g2(ageG) %>%
    as_tibble() %>%
    mutate(par = c("OR", "p.val", "ORlow", "ORhigh")) %>%
    gather("Region", "Value", 1:3) %>%
    mutate(Type = t)
})
comb_cont_Age <- Reduce(rbind, types_cont_Age)
comb_cont_Age$dataset <- "Epidelta"

age_cont_Sum_rel <- CpGsSum %>%
  dplyr::select(CpG, Direction) %>%
  mutate(Direction = factor(Direction, levels = c("Inverse", "Positive", "Both", "Non-significant"))) %>%
  left_join(dplyr::select(epideltadf, Dir, M1.change.estimate, CpG), by = "CpG") %>%
  left_join(dplyr::select(methyAnnot, CpG, Reliability), by = "CpG") %>%
  filter(Reliability > 0.4) %>%
  group_by(Direction, Dir) %>%
  summarize(n = n()) %>%
  spread(Dir, n) %>%
  ungroup() %>%
  rbind(filter(., Direction != "Non-significant") %>% 
          summarize_all(fsum)) %>%
  mutate(Direction = as.character(Direction),
         Direction = ifelse(is.na(Direction), "eQTM", Direction),
         Variable = Decreased + Increased,
         tot = Variable + Constant)

types_cont_Age_reliability <- lapply(types, function(t){
  rbind(filter(age_cont_Sum_rel, Direction == t),
        filter(age_cont_Sum_rel, Direction == "Non-significant")) %>%
    g2(ageG) %>%
    as_tibble() %>%
    mutate(par = c("OR", "p.val", "ORlow", "ORhigh")) %>%
    gather("Region", "Value", 1:3) %>%
    mutate(Type = t)
})
comb_cont_Age_reliability <- Reduce(rbind, types_cont_Age_reliability)
comb_cont_Age_reliability$dataset <- "Epidelta"


## Make combined plot
age_var_comb <- rbind(comb_cont_Age, combAge) %>%
  mutate(dataset = factor(dataset, levels = c("MeDALL", "Epidelta"))) %>%
  spread(par, Value) %>%
  mutate(Type = ifelse(Type == "eQTM", "All", Type),
         Type = factor(Type, levels = c("All", "Inverse", "Positive"))) %>%
  ggplot(aes(x = Region, y = OR, fill = Type)) + 
  geom_bar(stat = "identity", position=position_dodge(), color = "black") + 
  geom_errorbar(position=position_dodge(.9), width=.25, aes(ymin = ORlow, ymax = ORhigh)) +
  scale_y_continuous(trans = "log2", 
                     breaks = scales::trans_breaks("log2", function(x) round(2^x, 2))) +
  geom_hline(yintercept = 1) +
  scale_x_discrete(name = "Change in methylation with age", limits = ageG) +
  scale_fill_manual(name = "eCpGs type", values = c("#999999", "#E69F00", "#009E73"),
                    labels = c("All", "Inverse", "Positive")) +
  theme_bw() +
  facet_grid(~ dataset)


## Make combined plot
age_var_comb_rel <- rbind(comb_cont_Age_reliability, combAge_rel) %>%
  mutate(dataset = factor(dataset, levels = c("MeDALL", "Epidelta"))) %>%
  spread(par, Value) %>%
  mutate(Type = ifelse(Type == "eQTM", "All", Type),
         Type = factor(Type, levels = c("All", "Inverse", "Positive"))) %>%
  ggplot(aes(x = Region, y = OR, fill = Type)) + 
  geom_bar(stat = "identity", position=position_dodge(), color = "black") + 
  geom_errorbar(position=position_dodge(.9), width=.25, aes(ymin = ORlow, ymax = ORhigh)) +
  scale_y_continuous(trans = "log2", 
                     breaks = scales::trans_breaks("log2", function(x) round(2^x, 2))) +
  geom_hline(yintercept = 1) +
  scale_x_discrete(name = "Change in methylation with age", limits = ageG) +
  scale_fill_manual(name = "eCpGs type", values = c("#999999", "#E69F00", "#009E73"),
                    labels = c("All", "Inverse", "Positive")) +
  theme_bw() +
  facet_grid(~ dataset)

png("paper/CpGEnrich_ageVar_reliability.png", width = 3000, height = 1500, res = 300)
age_var_comb_rel
dev.off()

## Overlap age variability datasets
merged_age <- left_join(select(agedf, CpG, Dir), 
                        select(epideltadf, CpG, Dir, M1.change.p), 
                        by = "CpG")
mean(merged_age$Dir.x == merged_age$Dir.y)     


left_join(select(agedf, CpG, Dir), 
          select(epideltadf, CpG, Dir, M1.change.p), by = "CpG") %>%
  left_join(dplyr::select(methyAnnot, CpG, Reliability), by = "CpG") %>%
  filter(Reliability > 0.4) %>%
  mutate(concordant = Dir.x == Dir.y) %>%
  summarize(m = mean(concordant))


png("paper/CpGEnrich_ageVar_reliability_cont.png", width = 3000, height = 1500, res = 300)
as_tibble(methyAnnot) %>%
  mutate(CpG = Name) %>%
  select(CpG, Reliability) %>%
  left_join(select(CpGsSum, CpG, Type)) %>%
  mutate(Significant = ifelse(Type == "Non-significant", "non-eQTMs", "eQTMs")) %>%
  left_join(dplyr::select(agedf, Dir, CpG), by = "CpG") %>%
  mutate(Xu = ifelse(is.na(Dir), "Constant", Dir))  %>%
  select(-Dir) %>%
  left_join(dplyr::select(epideltadf, Dir, CpG), by = "CpG") %>%
  mutate(Epidelta = Dir) %>%
  filter(!is.na(Significant) & !is.na(Epidelta)) %>%
  select(Xu, Epidelta, CpG, Reliability, Significant) %>%
  gather(Database, Status, 1:2) %>%
  mutate(Database = ifelse(Database == "Xu", "MeDALL", "Epidelta"),
         Database = factor(Database, levels = c("MeDALL", "Epidelta"))) %>%
  ggplot(aes(x = Status, y = Reliability, fill = Significant)) +
  geom_violin() +
  scale_x_discrete(name = "Change in methylation") +
  scale_y_continuous(name = "Probe reliability") +
  scale_fill_manual(name = "CpG type", values = c("#999999", "#FFFFFF")) +
  facet_grid(~ Database) +
  theme_bw()
dev.off()


## Compare with other eQTM studies (Kennedy) PMID:  29914364 ####
eQTM_blood <- read.delim("data/Kennedy_eQTM_catalogue_wholeBlood.txt", as.is = TRUE)
eQTM_monocytes <- read.delim("data/Kennedy_eQTM_catalogue_Monocytes.txt", as.is = TRUE)

## Remove CpGs and Genes not present in HELIX
eQTM_blood.f <- subset(eQTM_blood, CpG.probe %in% unique(modC$CpG) & 
                         annot.gene %in% expGenes)
eQTM_monocytes.f <- subset(eQTM_monocytes, CpG.probe %in% unique(modC$CpG) & 
                         annot.gene %in% expGenes)

## Compare with main model
blood.merge <- eQTM_blood.f %>%
  mutate(CpG = CpG.probe, GeneSymbol_Affy = annot.gene) %>%
  inner_join(modC_Annot, by = c("CpG", "GeneSymbol_Affy")) %>%
  left_join(select(expAnnot, c("TC", "start", "stop")), by = "TC") %>%
  mutate(sigType = ifelse(sigPair, ifelse(p.val < 1e-11, "Both", "Main"), 
                          ifelse(p.val < 1e-11, "WholeBlood", "None")),
         sigType = factor(sigType, levels = c("Both", "None", "Main", "WholeBlood")),
         sigType2 = ifelse(sigPair, "Shared", "WholeBlood"))

monocytes.merge <- eQTM_monocytes.f %>%
  mutate(CpG = CpG.probe, GeneSymbol_Affy = annot.gene) %>%
  inner_join(modC_Annot, by = c("CpG", "GeneSymbol_Affy")) %>%
  mutate(sigType = ifelse(sigPair, ifelse(p.val < 1e-11, "Both", "Main"), 
                          ifelse(p.val < 1e-11, "Monocytes", "None")),
         sigType = factor(sigType, levels = c("Both", "None", "Main", "Monocytes")),
         sigType2 = ifelse(sigPair, "Shared", "Monocytes"))


## Map one pair of HELIX (lowest p-value) to each pair in GTP and MESA
blood.merge.uniq <- blood.merge %>% 
  group_by(exp.Probe, CpG.probe) %>%
  top_n(-1, p.value)

monocytes.merge.uniq <- monocytes.merge %>% 
  group_by(exp.Probe, CpG.probe) %>%
  top_n(-1, p.value)


blood.merge.uniq %>% group_by(sigType) %>% summarize(rFC = cor(FC, beta), 
                                                pFC = cor.test(FC, beta)$p.value,
                                                signConc = mean(sign(FC) == sign(beta)))
blood.merge.uniq %>% group_by(sigType2) %>% summarize(rFC = cor(FC, beta), 
                                                 pFC = cor.test(FC, beta)$p.value,
                                                 signConc = mean(sign(FC) == sign(beta)))


monocytes.merge.uniq %>% group_by(sigType) %>% summarize(rFC = cor(FC, beta), 
                                                    pFC = cor.test(FC, beta)$p.value,
                                                    signConc = mean(sign(FC) == sign(beta)))
monocytes.merge.uniq %>% group_by(sigType2) %>% summarize(rFC = cor(FC, beta), 
                                                     pFC = cor.test(FC, beta)$p.value,
                                                     signConc = mean(sign(FC) == sign(beta)))
table(blood.merge.uniq$sigType2)
table(monocytes.merge.uniq$sigType2)


## Comparison catalogue pairs in Kennedy ####
monocytes.merge$pair <- paste(monocytes.merge$CpG, monocytes.merge$TC)
blood.merge$pair <- paste(blood.merge$CpG, blood.merge$TC)
modC_Annot$MESA <- ifelse(paste(modC_Annot$CpG, modC_Annot$TC) %in% monocytes.merge$pair, 
                          "MESA", "not in MESA")
modC_Annot$GTP <- ifelse(paste(modC_Annot$CpG, modC_Annot$TC) %in% blood.merge$pair, 
                          "GTP", "not in GTP")
modC_Annot$adults <- ifelse(modC_Annot$sigPair, 
                           ifelse(modC_Annot$MESA == "MESA" | modC_Annot$GTP == "GTP", "Shared", "Children"), 
                           "Other")

modC_Annot %>% 
  filter(adults != "Other") %>%
  group_by(adults) %>%
  summarize(nCpG = length(unique(CpG)),
            nTC = length(unique(TC)))


modC_Annot %>% filter(sigPair == TRUE) %>% group_by(GTP) %>% 
  summarize(m = median(-log10(p.value)))
modC_Annot %>% filter(sigPair == TRUE) %>% group_by(MESA) %>% 
  summarize(m = median(-log10(p.value)))
modC_Annot %>% filter(sigPair == TRUE) %>% group_by(adults) %>% 
  summarize(m = median(-log10(p.value)))

modC_Annot %>% filter(sigPair == TRUE) %>% group_by(MESA) %>% 
  summarize(m = median(p.value))
modC_Annot %>% filter(sigPair == TRUE) %>% group_by(GTP) %>% 
  summarize(m = median(p.value))
modC_Annot %>% filter(sigPair == TRUE) %>% group_by(adults) %>% 
  summarize(m = median(p.value))


modC_comp$MESA <- ifelse(paste(modC_comp$CpG, modC_comp$TC) %in% monocytes.merge$pair, 
                         "p-value < 1e-5", "p-value > 1e-5")
modC_comp$GTP <- ifelse(paste(modC_comp$CpG, modC_comp$TC) %in% blood.merge$pair, 
                         "p-value < 1e-5", "p-value > 1e-5")
modC_comp$adult <- ifelse(modC_comp$sigPair, 
                           ifelse(modC_comp$MESA == "p-value < 1e-5" | modC_comp$GTP == "p-value < 1e-5", "Shared", "Children"), 
                           "Other")


wilcox.test(abs(subset(modC_comp, sigPair & adult == "Shared")$FC)/10,
        abs(subset(modC_comp, sigPair & adult == "Children")$FC)/10)

modC_Annot %>% filter(sigPair == TRUE) %>% group_by(adult) %>% 
  summarize(m = median(abs(FC)/10))

modC_comp %>% filter(sigPair == TRUE) %>% group_by(adult) %>% 
  summarize(m = median(SD))


modC_comp %>% filter(sigPair == TRUE) %>% ggplot(aes(x = abs(FC), color = adult)) +
  geom_density() + scale_x_continuous(limits = c(0, 6))
modC_comp %>% filter(sigPair == TRUE) %>% ggplot(aes(x = SD, color = adult)) +
  geom_density() + scale_x_continuous(limits = c(0, 3))
wilcox.test(abs(subset(modC_comp, sigPair & adult == "Shared")$SD),
        abs(subset(modC_comp, sigPair & adult == "Children")$SD))


wilcox.test(-log10(subset(modC_comp, sigPair & adult == "Shared")$p.value),
            -log10(subset(modC_comp, sigPair & adult == "Children")$p.value))


plot_all <- modC_comp %>% filter(sigPair == TRUE) %>% 
  mutate(Type = ifelse(adult == "Children", "Child-specific", "Age-shared")) %>%
  ggplot(aes(x = Distance, color = Type)) +
  geom_density() +
  theme_bw() + 
  scale_x_continuous(name = "eCpG-eGene's TSS distance",
                     breaks = c(-5e5, -2e5, 0, 2e5, 5e5), 
                     labels = c("-500Kb", "-250Kb", "0", "250Kb", "500Kb")) +
  scale_y_continuous(name = "Density") + 
  scale_color_discrete(name = "eQTM type") +
  theme(plot.title = element_text(hjust = 0.5))
png("paper/distance_distr_Kennedy.png", width = 2000, height = 1000, res = 300)
plot_all
dev.off()

modC_comp %>% filter(sigPair == TRUE) %>% 
  mutate(Type = ifelse(adult == "Children", "Children-specific", "Children and adult shared")) %>%
  group_by(Type) %>%
  summarize(m = median(Distance),
            q25 = quantile(Distance, 0.25),
            q75 = quantile(Distance, 0.75))
ks.test(subset(modC_comp, sigPair == TRUE & adult == "Children")$Distance,
        subset(modC_comp, sigPair == TRUE & adult != "Children")$Distance) 

### Plot overlap between datasets
adult_db <- full_join(select(blood.merge.uniq, exp.Probe, CpG.probe, sigType2),
                       select(monocytes.merge.uniq, exp.Probe, CpG.probe, sigType2),
                       by = c("exp.Probe", "CpG.probe")) %>%
  mutate(name = paste(exp.Probe, CpG.probe),
         sigType2.x = ifelse(is.na(sigType2.x), "None", sigType2.x),
         sigType2.y = ifelse(is.na(sigType2.y), "None", sigType2.y),
         GTP = sigType2.x != "None",
         MESA = sigType2.y != "None",
         HELIX = sigType2.x == "Shared" | sigType2.y == "Shared")
adult_over <- plot(euler(adult_db[, c("GTP", "MESA", "HELIX")], shape = "ellipse"), 
           fills = list(fill = c("cyan", "darksalmon", "darkseagreen"), alpha = 0.5),
     quantities = list(fontsize = 8), 
     main = "Adult eQTMs in HELIX")


helix_over <- modC_Annot %>% 
  filter(sigPair == TRUE) %>%
  mutate(HELIX = TRUE,
         GTP = GTP == "GTP",
         MESA = MESA == "MESA") %>%
  select(GTP, MESA, HELIX) %>%
  euler(shape = "ellipse") %>%
  plot(fills = list(fill = c("cyan", "darksalmon", "darkseagreen"), alpha = 0.5),
       quantities = list(fontsize = 8), 
       main = "HELIX eQTMs in adult cohorts")

bottom <- plot_grid(adult_over, helix_over, labels = c("B", "C"))

png("paper/AgeEffectPlot.png", width = 3000, height = 2000, res = 300)
plot_grid(age_var_comb, bottom, ncol = 1, labels = c("A", ""))
dev.off()

#### Prepare data in previous sections   ####
childCpGs <- unique(modC_Annot[modC_Annot$adults == "Children", ]$CpG)
sharedCpGs <- unique(modC_Annot[modC_Annot$adults == "Shared", ]$CpG)

length(intersect(childCpGs, sharedCpGs))
length(setdiff(sharedCpGs, childCpGs))
length(setdiff(childCpGs, sharedCpGs))

childCpGs.f <- setdiff(childCpGs, sharedCpGs)
sharedCpGs.f <- setdiff(sharedCpGs, childCpGs)


## Select genes
childTCs <- unique(modC_Annot[modC_Annot$adults == "Children", ]$TC)
childGenes <- filter(expAnnot, probeset_id %in% childTCs)$GeneSymbol_Affy
childGenes <- unique(unlist(strsplit(childGenes, ";")))

sharedTCs <- unique(modC_Annot[modC_Annot$adults == "Shared", ]$TC)
sharedGenes <- filter(expAnnot, probeset_id %in% sharedTCs)$GeneSymbol_Affy
sharedGenes <- unique(unlist(strsplit(sharedGenes, ";")))

length(intersect(sharedTCs, childTCs))
length(setdiff(sharedTCs, childTCs))
length(setdiff(childTCs, sharedTCs))

## Remove genes regulated by different CpGs in both models
childGenes.f <- setdiff(childGenes, sharedGenes)
sharedGenes.f <- setdiff(sharedGenes, childGenes)

## Compare with reliability ####
png("paper/kenney_comparison_reliability.png", width = 2000, height = 1000, res = 300)
data.frame(CpG = c(childCpGs, sharedCpGs), 
           Type = rep(c("Child-specific eCpGs", "Age-shared eCpGs"), 
                      c(length(childCpGs), length(sharedCpGs)))) %>%
  left_join(dplyr::select(methyAnnot, CpG, Reliability), by = "CpG") %>%
  ggplot(aes(x = Type, y = Reliability, fill = Type)) +
  geom_boxplot() +
  scale_x_discrete(name = "eCpG type") +
  scale_y_continuous(name = "Probe reliability") +
  scale_fill_manual(name = "", values = c("#F8766D", "#00BFC4")) +
  theme_bw() +
  theme(legend.position = "none")
dev.off()

## Compare with meQTLs ####
load("results/eQTLanalysis/comQTLs.Rdata") 
mQTLs <- read.table("data/ARIES_mQTLs.tab", header = TRUE, as.is = TRUE)
mQTLsH2 <- read.table("results/ARIES/mqtls.txt", header = TRUE, as.is = TRUE)

ARIESannot <- read.table("data/ariesmqtlsnps.bim", as.is = TRUE)
colnames(ARIESannot) <- c("chr", "SNP", "cm", "pos", "Ref", "Alt")

HELIXannot <- read.table("~/data/WS_HELIX/HELIX_preproc/gwas/Final_data_HRCimp_QC2/HELIX.impQC.rs.bim", as.is = TRUE)
colnames(HELIXannot) <- c("chr", "SNP", "cm", "pos", "Ref", "Alt")

comMQTLs <- mQTLsH2 %>%
  mutate(gene = CpG) %>%
  dplyr::select(SNP, gene, A1, A2, freq, b, se, p, N, r2) %>%
  semi_join(rbind(comCisQTL, comTransQTL), by = c("SNP", "gene")) %>%
  inner_join(left_join(mQTLs, dplyr::select(ARIESannot, SNP, Ref, Alt), by = "SNP"), 
             by = c("SNP", "gene")) %>%
  as_tibble()

comMQTLs.f <- comMQTLs %>%
  filter(!is.na(Ref)) %>%
  filter(!(sign(b) != sign(beta) & A1 == Ref)) %>%
  filter(!(sign(b) == sign(beta) & A1 == Alt))

meqtlCpGs <- unique(comMQTLs.f$gene)

meQTL_Sum <- data.frame(CpG = c(childCpGs, sharedCpGs), 
                        Type = rep(c("Children-specific", "Children and adult shared"), 
                                   c(length(childCpGs), length(sharedCpGs)))) %>%
  rbind(data.frame(CpG = filter(methyAnnot, !Name %in% .$CpG)$Name, 
                   Type = "None")) %>%
  mutate(meQTL = CpG %in% meqtlCpGs) %>%
  group_by(Type) %>%
  summarize(meQTLin = sum(meQTL), no_meQTL = sum(!meQTL))

 getOR(2:3, meQTL_Sum[-2, ]) ## Shared
 getOR(2:3, meQTL_Sum[-1, ]) ## children

## Compare ROADMAP chromatin states ####
 chromStates <- c("TssA", "TssAFlnk", "TxFlnk", "TxWk", "Tx", "EnhG", "Enh",
                  "ZNF.Rpts", "Het", "TssBiv", "BivFlnk", "EnhBiv", "ReprPC",
                  "ReprPCWk", "Quies")
 sum2 <- function(x) sum(!x, na.rm = TRUE)
 
chromSt_adult <-  data.frame(CpG = c(childCpGs, sharedCpGs),
                             eQTM = rep(c("Child-specific eCpGs", "Age-shared eCpGs"),
                                        c(length(childCpGs), length(sharedCpGs)))) %>%
  rbind(data.frame(CpG = filter(methyAnnot, !Name %in% .$CpG)$Name,
                   eQTM = "None")) %>%
  left_join(mutate(methyAnnot, CpG = Name) %>%  dplyr::select(CpG, eval(chromStates))) %>%
  filter(!is.na(Quies)) %>%
  group_by(eQTM) %>%
  summarize_at(chromStates, list(sum = sum, sum2 = sum2, prop = mean))


chromSt_enrich_adult <- lapply(c("Child-specific eCpGs", "Age-shared eCpGs"), function(t){
  rbind(filter(chromSt_adult, eQTM == t),
        filter(chromSt_adult, eQTM == "None")) %>%
    select(ends_with("sum"), ends_with("sum2")) %>%
    g(chromStates, cols = c("_sum", "_sum2")) %>%
    as_tibble() %>%
    mutate(par = c("OR", "p.val", "ORlow", "ORhigh")) %>%
    gather("Region", "Value", 1:15) %>%
    mutate(eQTM = t)
})
chromSt_enrich_adult <- Reduce(rbind, chromSt_enrich_adult)

chrom_adult <- chromSt_enrich_adult %>%
  spread(par, Value) %>%
  mutate(Group = factor(ifelse(Region %in% c("TssA", "TssAFlnk"), "TssProxProm",
                               ifelse(Region %in% c("Tx", "TxWk"), "ActTrans", 
                                      ifelse(Region %in% c("Enh", "EnhG"), "Enhancer", 
                                             ifelse(Region %in% c("TssBiv", "BivFlnk", "EnhBiv"), "BivReg", 
                                                    ifelse(Region %in% c("ReprPC", "ReprPCWk"), "ReprPoly", Region)
                                             )
                                      )
                               )
  ), 
  levels = c("TssProxProm", "TxFlnk", "ActTrans", "Enhancer", "ZNF.Rpts", "BivFlnk", "BivReg", "Het", "ReprPoly", "Quies")
  ),
  ) %>%
  ggplot(aes(x = Region, y = OR, fill = eQTM)) + 
  geom_bar(stat = "identity", position=position_dodge(), color= "black") + 
  geom_errorbar(position=position_dodge(.9), width=.25, aes(ymin = ORlow, ymax = ORhigh)) +
  scale_y_continuous(trans = "log2", 
                     breaks = scales::trans_breaks("log2", function(x) round(2^x, 2))) +
  scale_fill_discrete(name = "eCpG type") +
  geom_hline(yintercept = 1) +
  scale_x_discrete(name = "ROADMAP chromatin states") +
  facet_wrap(~ Group, scales = "free_x") +
  theme_bw() +
  theme(plot.title = element_text(hjust = 0.5))

png("paper/enrich_chromStates_adult.png", width = 2500, height = 1500, res = 300)
chrom_adult
dev.off()

chromSt_adult_prop <- chromSt_adult %>%
  select(eQTM, ends_with("prop")) %>%
  gather(categories, proportion, 2:(2+length(chromStates) - 1)) %>%
  ungroup() %>%
  mutate(Type = as.character(eQTM),
         Type = ifelse(Type == "None", "non-eQTMs", Type),
         Type = factor(Type, levels = c("non-eQTMs", "Age-shared eQTMs", "Child-specific eQTMs")),
         categories = gsub("_prop", "", categories),
         Group = factor(ifelse(categories %in% c("TssA", "TssAFlnk"), "TssProxProm",
                               ifelse(categories %in% c("Tx", "TxWk"), "ActTrans", 
                                      ifelse(categories %in% c("Enh", "EnhG"), "Enhancer", 
                                             ifelse(categories %in% c("TssBiv", "BivFlnk", "EnhBiv"), "BivReg", 
                                                    ifelse(categories %in% c("ReprPC", "ReprPCWk"), "ReprPoly", categories)
                                             )
                                      )
                               )
         ), 
         levels = c("TssProxProm", "TxFlnk", "ActTrans", "Enhancer", "ZNF.Rpts", "BivFlnk", "BivReg", "Het", "ReprPoly", "Quies")
         ),
         categories = factor(categories, levels = chromStates)) %>%
  ggplot(aes(x = categories, y = proportion*100, fill = Type)) +
  geom_bar(stat = "identity", position = position_dodge(), color = "black") +
  scale_x_discrete(name = "ROADMAP chromatin states") +
  facet_wrap(~ Group, scales = "free_x") +  
  scale_y_continuous(name = "Proportion of CpGs (%)") +
  scale_fill_manual(name = "CpG type", values = c("#FFFFFF", "#F8766D", "#00BFC4")) +
  theme_bw()

png("paper/chromStatesProp_age.png", width = 2500, height = 2000, res = 300)
chromSt_adult_prop
dev.off()
```
