## SupplementalCode for "Identification of autosomal cis expression quantitative trait methylation (cis eQTMs) in children’s blood": eQTM_genetics.R

```
#'##############################################################################
#' Genetic variants and eQTMs
#' This script contains the code for
#'  - Heritability
#'  - Association with genetics
#'##############################################################################

# Load data and libraries ####
library(ggplot2)
library(dplyr)
library(cowplot)
library(tidyr)
library(GenomicRanges)
library(snpStats)
library(MASS)
library(topGO)

load("results/MethComBatExpResidualsCellAdj/allres_simP_cpgs.Rdata")
load("results/preprocessFiles/allOverlaps.Rdata")
load("results/preprocessFiles/methyAnnotation.Rdata")
load("results/preprocessFiles/gexpAnnotation.Rdata")

# Get useful variables ####
methyAnnot <- methyAnnot %>%
  as_tibble() %>%
  mutate(CpG = Name)

CpGsNum <- df %>%
  group_by(CpG) %>%
  summarise(nTC = sum(sigPair)) %>%
  mutate(nCat = ifelse(nTC > 3, "4+", nTC)) %>%
  left_join(dplyr::select(methyAnnot, CpG, Reliability), by = "CpG")
sigDf <- df %>%
  as_tibble() %>%
  filter(sigPair)

codingTCs <- subset(expAnnot, Coding == "coding")$transcript_cluster_id


CpGsSum <- df %>%
  group_by(CpG) %>%
  summarise(Type = ifelse(sum(sigPair) == 0, "Non-significant",
                          ifelse(sum(sigPair) == 1, "Mono", "Multi")),
            Direction = ifelse(sum(sigPair) == 0, "Non-significant",
                               ifelse(all(FC[sigPair] > 0), "Positive", 
                                      ifelse(all(FC[sigPair] < 0), "Inverse", "Both")))) %>%
  mutate(Combined = ifelse(Type == "Non-significant", 
                           "Non-significant", 
                           paste(Type, Direction, sep = "_")))


## Heritability ####
## Create summary tibble
h2df <- read.csv("data/HeritabilityCpGs", as.is = TRUE)
herm <- h2df %>%
  as_tibble() %>%
  ## Remove CpGs with NA in h2
  filter(!is.na(h2_total)) %>%
  mutate(CpG = cgid) %>%
  inner_join(CpGsNum, h2df, by = "CpG") 

h2tot <- herm %>%
  mutate(type = ifelse(nCat == 0, "non eQTMs", "eQTMs")) %>%
  ggplot(aes(x = nCat, y = h2_total, fill = type)) + 
  geom_boxplot() +
  scale_fill_manual(values=c("grey", "white")) +
  geom_hline(yintercept = c(0.2, 0.5), linetype="dashed", colour = "blue") + 
  scale_x_discrete(name = "Number of Genes associated with a CpG") +
  scale_y_continuous(name = "Total heritability", limits = c(0, 1)) +
  theme_bw() +
  theme(plot.title = element_text(hjust = 0.5),
        legend.position = "none")

h2SNP <- herm %>%
  mutate(type = ifelse(nCat == 0, "non eQTMs", "eQTMs")) %>%
  ggplot(aes(x = nCat, y = h2_SNPs, fill = type)) + 
  geom_boxplot() +
  scale_fill_manual(values=c("grey", "white")) +
  geom_hline(yintercept = c(0.2, 0.5), linetype="dashed", colour = "blue") + 
  scale_x_discrete(name = "Number of Genes associated with a CpG") +
  scale_y_continuous(name = "SNP heritability", limits = c(0, 1)) +
  theme_bw() +
  theme(plot.title = element_text(hjust = 0.5),
        legend.position = "none")

herm %>%
  mutate(sig = ifelse(nTC == 0, "Non-significant", "Significant"),
         sig = factor(sig, levels = c("Non-significant", "Significant"))) %>%
  group_by(sig) %>%
  summarize(m = median(h2_total))

wilcox.test(subset(herm, nTC == 0)$h2_total,
            subset(herm, nTC != 0)$h2_total, conf.int = TRUE)


herm %>%
  filter(nTC != 0) %>%
  lm(formula = h2_total ~  nTC) %>%
  summary()

herm %>%
  mutate(sig = ifelse(nTC == 0, "Non-significant", "Significant"),
         sig = factor(sig, levels = c("Non-significant", "Significant"))) %>%
  group_by(sig) %>%
  summarize(m = median(h2_SNPs))

wilcox.test(subset(herm, nTC == 0)$h2_SNPs,
            subset(herm, nTC != 0)$h2_SNPs, conf.int = TRUE)
 

herm %>%
  filter(nTC != 0) %>%
  lm(formula = h2_SNPs ~  nTC) %>%
  summary()

h2tot.rel <- herm %>%
  filter(!is.na(Reliability) & Reliability >= 0.4) %>%
  mutate(type = ifelse(nCat == 0, "non eQTMs", "eQTMs")) %>%
  ggplot(aes(x = nCat, y = h2_total, fill = type)) + 
  geom_boxplot() +
  scale_fill_manual(values=c("grey", "white")) +
  geom_hline(yintercept = c(0.2, 0.5), linetype="dashed", colour = "blue") + 
  scale_x_discrete(name = "Number of Genes associated with a CpG") +
  scale_y_continuous(name = "Total heritability", limits = c(0, 1)) +
  theme_bw() +
  theme(plot.title = element_text(hjust = 0.5),
        legend.position = "none")

h2SNP.rel <- herm %>%
  filter(!is.na(Reliability) & Reliability >= 0.4) %>%
  mutate(type = ifelse(nCat == 0, "non eQTMs", "eQTMs")) %>%
  ggplot(aes(x = nCat, y = h2_SNPs, fill = type)) + 
  geom_boxplot() +
  scale_fill_manual(values=c("grey", "white")) +
  geom_hline(yintercept = c(0.2, 0.5), linetype="dashed", colour = "blue") + 
  scale_x_discrete(name = "Number of Genes associated with a CpG") +
  scale_y_continuous(name = "SNP heritability", limits = c(0, 1)) +
  theme_bw() +
  theme(plot.title = element_text(hjust = 0.5),
        legend.position = "none")

png("paper/eQTMs_heritability_rel.png", width = 2500, height = 750, res = 300)
plot_grid(h2tot.rel, h2SNP.rel, ncol = 2, labels = "AUTO")
dev.off()

wilcox.test(subset(herm, nTC == 0 & Reliability >= 0.4)$h2_total,
            subset(herm, nTC != 0 & Reliability >= 0.4)$h2_total, conf.int = TRUE)


wilcox.test(subset(herm, nTC == 0 & Reliability >= 0.4)$h2_SNPs,
            subset(herm, nTC != 0 & Reliability >= 0.4)$h2_SNPs, conf.int = TRUE)


## Common mQTLs between ARIES and HELIX ####
load("results/eQTLanalysis/comQTLs.Rdata")
mQTLs <- read.table("data/ARIES_mQTLs.tab", header = TRUE, as.is = TRUE)
mQTLsH2 <- read.table("results/ARIES/mqtls.txt", header = TRUE, as.is = TRUE)

## Select pairs detected in previous analysis from ARIES pipeline
## Merge with ARIES original data
ARIESannot <- read.table("data/ariesmqtlsnps.bim", as.is = TRUE)
colnames(ARIESannot) <- c("chr", "SNP", "cm", "pos", "Ref", "Alt")

HELIXannot <- read.table("~/data/WS_HELIX/HELIX_preproc/gwas/Final_data_HRCimp_QC2/HELIX.impQC.rs.bim", as.is = TRUE)
colnames(HELIXannot) <- c("chr", "SNP", "cm", "pos", "Ref", "Alt")

comMQTLs <- mQTLsH2 %>%
  mutate(gene = CpG) %>%
  dplyr::select(SNP, gene, A1, A2, freq, b, se, p, N, r2) %>%
  semi_join(rbind(comCisQTL, comTransQTL), by = c("SNP", "gene")) %>%
  inner_join(left_join(mQTLs, dplyr::select(ARIESannot, SNP, Ref, Alt), by = "SNP"), 
             by = c("SNP", "gene")) %>%
  as_tibble()

## Remove pairs where effect direction do not match
comMQTLs.f <- comMQTLs %>%
  filter(!is.na(Ref)) %>%
  filter(!(sign(b) != sign(beta) & A1 == Ref)) %>%
  filter(!(sign(b) == sign(beta) & A1 == Alt))

comCpGs <- unique(comMQTLs.f$gene)

meQTLTab <- CpGsNum %>%
  mutate(mQTL = CpG %in% comCpGs,
         cisQTL = CpG %in% comCisQTL$gene,
         transQTL = CpG %in% comTransQTL$gene) %>%
  group_by(nCat) %>%
  summarize_if(is.logical, list(sum, mean))

sum(sigDf$CpG %in% comCpGs)
sum(unique(sigDf$CpG) %in% comCpGs)

meQTLTab2 <- CpGsNum %>%
  mutate(mQTLin = CpG %in% comCpGs,
         mQTLOut = !CpG %in% comCpGs) %>%
  group_by(nCat) %>%
  summarize_if(is.logical, sum)

ORs <- sapply(c("1", "2", "3", "4+"), function(x){
  x2 <- data.matrix(subset(meQTLTab2, nCat %in% c("0", x))[, 3:2])
  OR <- x2[1]/x2[2]/x2[3]*x2[4]
  p <- chisq.test(x2)$p.value
  c(OR = OR, p = p)
  
})
tmp <- meQTLTab2 %>%
  mutate(cat = ifelse(nCat %in% c("0", "1"), nCat, "2+")) %>%
  filter(cat != "1") %>%
  group_by(cat) %>%
  summarize(mQTLin = sum(mQTLin),
            mQTLOut = sum(mQTLOut))
x2 <- data.matrix(tmp[, 3:2])
OR <- x2[1]/x2[2]/x2[3]*x2[4]
p <- chisq.test(x2)$p.value


x2 <- meQTLTab2 %>% 
  mutate(cat = ifelse(nCat == 0, "noeQTM", "eQTM")) %>%
  group_by(cat) %>%
  summarize(mQTLin = sum(mQTLin), mQTLOut = sum(mQTLOut)) %>%
  data.matrix()
x2 <- x2[, -1]
x2[1]/x2[2]/x2[3]*x2[4] 

         
comMQTLs.f %>%
  dplyr::select(SNP, gene) %>%
  distinct() %>%
  group_by(gene) %>%
  summarize(CpGs = n()) %>%
  pull(., CpGs) %>%
  summary()

comMQTLs.f %>%
  dplyr::select(SNP, gene) %>%
  distinct() %>%
  group_by(gene) %>%
  summarize(CpGs = n()) %>%
  mutate(eQTM = ifelse(gene %in% unique(sigDf$CpG), "eQTM", "non-eQTM")) %>%
  group_by(eQTM) %>%
  summarize(m = median(CpGs),
            l = quantile(CpGs, 0.25),
            h = quantile(CpGs, 0.75))

meQTL_p <- CpGsNum %>%
  mutate(mQTL = CpG %in% comCpGs) %>%
  group_by(nCat) %>%
  summarize(prop = mean(mQTL)*100) %>%
  mutate(type = ifelse(nCat == 0, "non-eQTMs", "eQTMs")) %>%
  ggplot(aes(x = nCat, y = prop,  fill = type)) +
  geom_bar(position = "dodge", stat = "identity", color = "black") +
  scale_fill_manual(values=c("grey", "white")) +
  scale_x_discrete(name = "Number of Genes associated with a CpG") +
  scale_y_continuous(name = "CpGs with meQTLs (%)") +
  theme_bw() +
  theme(legend.position = "none")
  

png("paper/eQTMsGenetics.png", width = 3500, height = 2000, res = 300)
plot_grid(plot_grid(h2tot, h2SNP, nrow = 2, labels = "AUTO"), meQTL_p, ncol = 2, labels = c("", "C"))
dev.off()

meQTLTab2 <- CpGsNum %>%
  mutate(mQTLin = CpG %in% comCpGs,
         mQTLOut = !CpG %in% comCpGs) %>%
  group_by(nCat) %>%
  summarize_if(is.logical, sum)

tab.rel <- CpGsNum %>%
  filter(!is.na(Reliability) & Reliability >= 0.4) %>%
  mutate(mQTLin = CpG %in% comCpGs,
         mQTLOut = !CpG %in% comCpGs) %>%
  group_by(nCat) %>%
  summarize_if(is.logical, sum) %>% 
  mutate(cat = ifelse(nCat == 0, "noeQTM", "eQTM")) %>%
  group_by(cat) %>%
  summarize(mQTLin = sum(mQTLin), mQTLOut = sum(mQTLOut)) %>%
  data.matrix()
tab.rel <- tab.rel[, -1]
tab.rel[1]/tab.rel[2]/tab.rel[3]*tab.rel[4] 


meQTL_p_rel <- CpGsNum %>%
  filter(!is.na(Reliability) & Reliability >= 0.4) %>%
  mutate(mQTL = CpG %in% comCpGs) %>%
  group_by(nCat) %>%
  summarize(prop = mean(mQTL)*100) %>%
  mutate(type = ifelse(nCat == 0, "non eQTMs", "eQTMs")) %>%
  ggplot(aes(x = nCat, y = prop,  fill = type)) +
  geom_bar(position = "dodge", stat = "identity", color = "black") +
  scale_fill_manual(values=c("grey", "white")) +
  scale_x_discrete(name = "Number of Genes associated with a CpG") +
  scale_y_continuous(name = "CpGs with meQTLs (%)") +
  theme_bw() +
  theme(legend.position = "none")

png("paper/eQTMs_meqtlProp_rel.png", width = 2000, height = 1000, res = 300)
meQTL_p_rel
dev.off()

## meQTLs vs reliability
png("paper/eQTMs_genetics_reliability.png", width = 2000, height = 1000, res = 300)

CpGsNum %>%
  filter(nTC > 0) %>%
  mutate(mQTL = ifelse(CpG %in% comCpGs, "With meQTLs", "Without meQTLs")) %>%
  ggplot(aes(x = mQTL, y = Reliability)) +
  geom_boxplot() +
  scale_x_discrete(name = "eCpG type") +
  scale_y_continuous(name = "Probe reliability") +
  theme_bw() +
  theme(legend.position = "none")
dev.off()


# Integrate with gene expression ####
load("results/eQTLanalysis/eQTLs.Rdata")
eQTL <- rbind(gexpme$cis$eqtls, gexpme$trans$eqtls) %>%
  mutate(TC = gene) %>%
  dplyr::select(-gene)

## Merge all associations 
## Remove non-coherent associations
mergedDf <- rbind(comCisQTL, comTransQTL) %>%
  mutate(CpG = gene) %>%
  semi_join(comMQTLs.f, by = c("SNP", "gene")) %>%
  inner_join(sigDf, by = "CpG") %>%
  inner_join(eQTL, by = c("snps", "TC")) %>%
  dplyr::select(-sigPair) %>%
  filter(sign(beta.x)*sign(FC) != sign(beta.y)) %>%
  tibble()

length(unique(paste(mergedDf$CpG, mergedDf$TC)))
length(unique(paste(mergedDf$CpG, mergedDf$TC)))/nrow(sigDf)
length(unique(mergedDf$CpG))
length(unique(mergedDf$CpG))/length(unique(sigDf$CpG))
length(unique(mergedDf$TC))
length(unique(mergedDf$TC))/length(unique(sigDf$TC))
sum(unique(mergedDf$TC) %in% codingTCs)
sum(unique(mergedDf$TC) %in% codingTCs)/sum(unique(sigDf$TC) %in% codingTCs)


mergedDf <- rbind(comCisQTL, comTransQTL) %>%
  mutate(CpG = gene) %>%
  dplyr::select(-gene, -SNP) %>%
  inner_join(sigDf, by = "CpG") %>%
  inner_join(eQTL, by = c("snps", "TC")) %>%
  dplyr::select(-sigPair)

mergedDf %>%
  dplyr::select(CpG, TC) %>%
  distinct() %>%
  group_by(TC) %>%
  summarize(CpGs = n()) %>%
  pull(., CpGs) %>%
  summary()

mergedDf %>%
  dplyr::select(snps, TC) %>%
  distinct() %>%
  group_by(TC) %>%
  summarize(CpGs = n()) %>%
  pull(., CpGs) %>%
  summary()


mergedDf %>%
  dplyr::select(snps, CpG) %>%
  distinct() %>%
  group_by(CpG) %>%
  summarize(CpGs = n()) %>%
  pull(., CpGs) %>%
  summary()

mergedDf %>%
  dplyr::select(TC, CpG) %>%
  distinct() %>%
  group_by(CpG) %>%
  summarize(CpGs = n()) %>%
  pull(., CpGs) %>%
  summary()
# Min. 1st Qu.  Median    Mean 3rd Qu.    Max.
# 1.00    1.00    1.00    1.87    2.00   21.00


### Example
# SNP: rs11585123
# CpG: cg15580684
# Gene: AJAP1 (TC01000080.hg.1)

plink <- read.plink("/home/isglobal.lan/cruiz/data/WS_HELIX/HELIX_preproc/gwas/Final_data_HRCimp_QC2/HELIX.impQC.rs")
load("results/preprocessFiles/Methylation_GRSet.RData")
colnames(gset) <- gset$HelixID

load("results/preprocessFiles/Expression_SE_residuals.RData")
colnames(se) <- se$HelixID

## Common samples
comSamps <- intersect(intersect(rownames(plink$geno), colnames(gset)), colnames(se))

alleles <- plink$map["rs11585123", c("allele.1", "allele.2")]
alleles <- c(paste0(alleles[1], alleles[1]),
             paste0(alleles[1], alleles[2]),
             paste0(alleles[2], alleles[2]))

dat <- data.frame(geno = as.numeric(plink$geno[comSamps, "rs11585123"]),
                  methy = as.numeric(getBeta(gset["cg15580684", comSamps])),
                  gexp = as.numeric(assay(se["TC01000080.hg.1", comSamps])))
dat$geno <- alleles[dat$geno]

sm <- ggplot(dat, aes(x = factor(geno), y = methy)) + 
  geom_boxplot() +
  scale_x_discrete(name = "rs11585123") +
  scale_y_continuous(name = "cg15580684") +
  theme_bw()

sg <- ggplot(dat, aes(x = factor(geno), y = gexp)) + 
  geom_boxplot() +
  scale_x_discrete(name = "rs11585123") +
  scale_y_continuous(name = "TC01000080.hg.1") +
  theme_bw()

  
me <- ggplot(dat, aes(x = methy, y = gexp)) + 
  geom_point() +
  scale_x_continuous(name = "cg15580684") +
  scale_y_continuous(name = "TC01000080.hg.1") +
  geom_smooth(method = "lm") +
  theme_bw()
png("paper/eQTMstrio.png", width = 3500, height = 2000, res = 300)
plot_grid(plot_grid(sm, sg, nrow = 1), me, nrow = 2)
dev.off()

## Run enrichment analysis genes in trios ####
## Copy functions from eQTM_interpretation.R
snpGenes <- df %>%
  dplyr::select(TC) %>%
  distinct() %>%
  mutate(sig = factor(ifelse(TC %in% unique(mergedDf$TC), 1, 0)))
snpGos <- computeGOs(snpGenes)
## Genes used in GOs
snpGos$go@geneData["Significant"]

save(snpGos, file = "paper/snpGOs.Rdata")
```
