## SupplementalCode for "Identification of autosomal cis expression quantitative trait methylation (cis eQTMs) in children’s blood": eQTM_interpretation.R

```
#'##############################################################################
#' Generate eQTM interpretation figures and tables
#' This script contains the code for
#'  - Gene enrichment
#'  - CpG enrichment 
#'  - eQTMs and genes and CpGs changing during childhood
#'  - Overlap with EWAS atlas
#'##############################################################################

# Load data and libraries ####
library(topGO)
library(ggplot2)
library(cowplot)
library(tidyr)
library(hta20transcriptcluster.db)
library(openxlsx)
library(GenomicRanges)
library(dplyr)

load("results/MethComBatExpResidualsCellAdj/allres_simP_cpgs.Rdata")
load("results/preprocessFiles/allOverlaps.Rdata")
load("results/preprocessFiles/methyAnnotation.Rdata")
load("results/preprocessFiles/gexpAnnotation.Rdata")

# Get useful variables ####
methyAnnot <- methyAnnot %>%
  as_tibble() %>%
  mutate(CpG = Name)

CpGsSum <- df %>%
  group_by(CpG) %>%
  summarise(Type = ifelse(sum(sigPair) == 0, "Non-significant",
                          ifelse(sum(sigPair) == 1, "Mono", "Multi")),
            Direction = ifelse(sum(sigPair) == 0, "Non-significant",
                               ifelse(all(FC[sigPair] > 0), "Positive", 
                                      ifelse(all(FC[sigPair] < 0), "Inverse", "Both")))) %>%
  mutate(Combined = ifelse(Type == "Non-significant", 
                           "Non-significant", 
                           paste(Type, Direction, sep = "_"))) %>%
  left_join(dplyr::select(methyAnnot, CpG, Reliability), by = "CpG")


## Gene enrichment ####
### Define functions
#' Compute GOs compute GO using two tests: 
#'  - classic: get p-value for each GO independent of the other
#'  - weight01: takes into account the GO hierarchy
## We selected only those terms with a p-value < 0.01 in both methods.

getGOdata <- function(df){
  genesv <- df$sig
  names(genesv) <- df$TC
  
  Data <- new("topGOdata", 
              description = "GO analysis of TCs",
              ontology = "BP",
              allGenes = genesv,
              annot = annFUN.db,
              nodeSize = 10,
              affyLib = "hta20transcriptcluster.db")
}

computeGOs <- function(df){
  
  Data <- getGOdata(df)
  mixed <- runTest(Data, algorithm = "weight01", statistic = "fisher")
  
  
  finTab <- GenTable(Data, 
                     w0 = mixed,
                     orderBy = "w0",
                     topNodes = length(score(mixed)))
  list(go = mixed, table = finTab)
}

runGO <- function(annot, df, filter){
  selCpGs <- filter(annot, Combined == filter)$CpG
  selGenes <- df %>%
    group_by(TC) %>%
    summarize(sig = factor(
      ifelse(any(CpG %in% selCpGs & sigPair), 1, 0)))
  go <- computeGOs(selGenes)
  go
}


## All genes
allGenes <- df %>%
  group_by(TC) %>%
  summarize(sig = factor(ifelse(any(sigPair), 1, 0)))
allGos <- computeGOs(allGenes)
allGOData <- getGOdata(allGenes)

## Save as Rdata data and process in local
save(allGos, file = "paper/GOobjects.Rdata")

## Genes in eQTMs from reliable probes
reliableGenes <- methyAnnot %>%
  select(Reliability, CpG) %>%
  right_join(df, by = "CpG") %>%
  group_by(TC) %>%
  summarize(sig = factor(ifelse(any(sigPair & Reliability > 0.4), 1, 0)))
reliableGos <- computeGOs(reliableGenes)
save(reliableGos, file = "paper/reliableGOobjects.Rdata")


## Genes used in GOs
allGos$go@geneData["Significant"]

## GOs changed
sum(allGos$table$w0 < 0.001)


## It does not work on server. Run locally.
library(GOfuncR)
library(topGO)
library(dplyr)
server <- "/run/user/1000/gvfs/sftp:host=isgws06.isglobal.lan,user=cruiz/home/isglobal.lan/cruiz/data/WS_HELIX/HELIX_analyses/expr_met_SM/paper/"

load(paste0(server, "GOobjects.Rdata"))
load(paste0(server, "reliableGOobjects.Rdata"))
load(paste0(server, "snpGOs.Rdata"))

## Define categories for GO terms
## Use http://amigo.geneontology.org/amigo/dd_browse and only select GOs > 10 genes in H. Sapiens.
## Exclude terms with regulation
##' Immune: 
##' - immune system process
##' - cell killing
##' Use level one/two parent
##' 
##' Adaptive immunity:
##' - adaptive immune response
##' - B cell selection
##' - T cell selection
##' - antigen processing and presentation
##' - B killer cell activation involved in immune response
##' - T cell activation involved in immune response
##' - complement activation
##' - complement-dependent cytotoxicity
##' - T cell mediated cytotoxicity
##' - B cell mediated immunity
##' - T cell mediated immunity
##' - opsonization
##' - B cell activation involved in immune response
##' - T cell activation involved in immune response
##' - B cell activation
##' - T cell activation
##' - B cell activation involved in immune response
##' - T cell activation involved in immune response
##' - B cell differentiation
##' - T cell differentiation
##' - B cell proliferation
##' - T cell proliferation
##' - lymphocyte homeostasis
##' - T cell extravasation
##' - lymphocyte migration
##' - lymphocyte coestimulation
##' - immunoglobulin production
##' - T cell cytokine production
##' - humoral immune response mediated by circulating immunoglobulin
##' - antigen receptor-mediated signaling pathway
##' 
##' 
##' Innate immunity:
##' - innate immune response
##' - astrocyte activation involved in immune response
##' - endothelial cell activation involved in immune response
##' - myeloid cell activation involved in immune response
##' - natural killer cell activation involved in immune response
##' - natural killer cell mediated cytotoxicity
##' - neutrophil mediated cytotoxicity
##' - immune response-regulating cell surface receptor signaling pathway involved in phagocytosis
##' - leukocyte degranulation
##' - dendritic cell cytokine production
##' - myeloid leukocyte mediated immunity
##' - natural killer cell mediated immunity
##' - myeloid cell activation involved in immune response
##' - natural killer cell activation involved in immune response
##' - leukocyte activation involved in inflammatory response
##' - natural killer cell activation involved in immune response
##' - natural killer cell differentiation
##' - natural killer cell proliferation
##' - natural killer cell activation
##' - myeloid leukocyte activation
##' - neutrophil homeostasis
##' - nuetrophil extravasation
##' - dendritic cell migration
##' - mononuclear cell migration
##' - myeloid leukocyte migration
##' - myeloid cell homeostasis
##' - dendritic cell cytokine production
##' - myeloid leukocyte cytokine production
##' - antimicrobial humoral response
##' - inflammatory response to antigenic stimulus

immune <- c("immune system process", "cell killing")
adaptive <- c("adaptive immune response",
              "antigen receptor-mediated signaling pathway",
              "B cell selection",
              "T cell selection",
              "antigen processing and presentation",
              "B killer cell activation involved in immune response",
              "T cell activation involved in immune response",
              "complement activation",
              "complement-dependent cytotoxicity",
              "T cell mediated cytotoxicity",
              "B cell mediated immunity",
              "T cell mediated immunity",
              "opsonization",
              "B cell activation involved in immune response",
              "T cell activation involved in immune response",
              "B cell activation",
              "T cell activation",
              "B cell activation involved in immune response",
              "T cell activation involved in immune response",
              "B cell differentiation",
              "T cell differentiation",
              "B cell proliferation",
              "T cell proliferation",
              "lymphocyte homeostasis",
              "T cell extravasation",
              "lymphocyte migration",
              "lymphocyte coestimulation",
              "immunoglobulin production",
              "T cell cytokine production",
              "humoral immune response mediated by circulating immunoglobulin")
innate <- c("innate immune response",
            "astrocyte activation involved in immune response",
            "endothelial cell activation involved in immune response",
            "myeloid cell activation involved in immune response",
            "natural killer cell activation involved in immune response",
            "natural killer cell mediated cytotoxicity",
            "neutrophil mediated cytotoxicity",
            "immune response-regulating cell surface receptor signaling pathway involved in phagocytosis",
            "leukocyte degranulation",
            "dendritic cell cytokine production",
            "myeloid leukocyte mediated immunity",
            "natural killer cell mediated immunity",
            "myeloid cell activation involved in immune response",
            "natural killer cell activation involved in immune response",
            "leukocyte activation involved in inflammatory response",
            "natural killer cell activation involved in immune response",
            "natural killer cell differentiation",
            "natural killer cell proliferation",
            "natural killer cell activation",
            "myeloid leukocyte activation",
            "neutrophil homeostasis",
            "nuetrophil extravasation",
            "dendritic cell migration",
            "mononuclear cell migration",
            "myeloid leukocyte migration",
            "myeloid cell homeostasis",
            "dendritic cell cytokine production",
            "myeloid leukocyte cytokine production",
            "antimicrobial humoral response",
            "inflammatory response to antigenic stimulus")

top_GOs <- subset(get_child_nodes("GO:0008150"), distance == 1)$child_name
celProc_GOs <- subset(get_child_nodes("GO:0009987"), distance == 1)$child_name

sumF <- function(x, topGOs){
  selGOs <- topGOs[topGOs %in% x]
  paste(selGOs, collapse = ";")
}


addImmunityInfo <- function(tab){
  get_parent_nodes(tab$GO.ID) %>%
    group_by(child_go_id) %>%
    summarize(parent = sumF(parent_name, topGOs = top_GOs), 
              immune = ifelse(any(adaptive %in% parent_name), 
                              "adaptive",
                              ifelse(any(innate %in% parent_name), 
                                     "innate",
                                     "undetermined")),
              GO_term = head(parent_name, 1)) %>%
    mutate(immune = ifelse(immune == "undetermined" & parent != "immune",
                           NA, immune),
           GO.ID = child_go_id) %>%
    right_join(tab)  %>%
    select(GO.ID, GO_term, w0, parent, immune)
}

## Select GOs with p-value < 0.001
allMod <- addImmunityInfo(subset(allGos$table, as.numeric(w0) < 0.001))
snpMod <- addImmunityInfo(subset(snpGos$table, as.numeric(w0) < 0.001))
relMod <- addImmunityInfo(subset(reliableGos$table, as.numeric(w0) < 0.001))

write.table(allMod[, c("GO.ID", "GO_term", "w0", "parent", "immune")], 
            file = paste0(server, "/GOsAllGenes.txt"), 
            quote = FALSE, col.names = TRUE, sep = "\t", row.names = FALSE)
write.table(snpMod[, c("GO.ID", "GO_term", "w0", "parent", "immune")], 
            file = paste0(server, "/GOsGenesSNPs.txt"), 
            quote = FALSE, col.names = TRUE, sep = "\t", row.names = FALSE)
write.table(relMod[, c("GO.ID", "GO_term", "w0", "parent", "immune")], 
            file = paste0(server, "/GOsGenesReliable.txt"), 
            quote = FALSE, col.names = TRUE, sep = "\t", row.names = FALSE)


# CpG Enrichment ####
types <- c("eQTM", "Positive", "Inverse")

## Define vars and function
getOR <- function(cols, df){
  t <- data.matrix(df[, cols])
  or <- t[1, 1]/(t[1, 2])/(t[2, 1])*t[2, 2]
  p.val <- chisq.test(t)$p.value
  ORl <- log(or)
  SEl <- sqrt(1/t[1, 1] + 1/t[1, 2] + 1/t[2, 1] + 1/t[2, 2])
  c(OR = or, p.val = p.val, ORm = exp(ORl - 1.96*SEl), 
    ORM = exp(ORl + 1.96*SEl))
}

g <- function(x, gpos, cols = c("in", "ou")){
  sapply(gpos, function(y) getOR(paste0(y, cols), df = x))
}


## Methylation levels ####
fsum <- function(x){
  if ( is.factor(x))
    "eQTM"
  else {
    sum(x)
  }
}

methMethLevels <- as_tibble(methyAnnot) %>%
  mutate(CpG = Name) %>%
  select(CpG, median_cat) %>%
  right_join(CpGsSum, by = "CpG") %>%
  mutate(Direction = factor(Direction, levels = c("Inverse", "Positive", "Both", "Non-significant"))) %>%
  group_by(Direction, median_cat) %>%
  summarize(n = n()) %>%
  spread(median_cat, n) %>%
  ungroup() %>%
  rbind(filter(., Direction != "Non-significant") %>% 
            summarize_all(fsum)) %>%
  mutate(Direction = as.character(Direction),
         Direction = ifelse(is.na(Direction), "eQTM", Direction),
         tot = low + medium + high)

cats <- c("low", "medium", "high")

## Define vars and function
getOR2 <- function(col, df){
  t <- data.matrix(df[, c(col, colnames(df)[ncol(df)])])
  t[, 2] <- t[, 2] - t[, 1]
  or <- t[1, 1]/(t[1, 2])/(t[2, 1])*t[2, 2]
  p.val <- chisq.test(t)$p.value
  ORl <- log(or)
  SEl <- sqrt(1/t[1, 1] + 1/t[1, 2] + 1/t[2, 1] + 1/t[2, 2])
  c(OR = or, p.val = p.val, ORm = exp(ORl - 1.96*SEl), 
    ORM = exp(ORl + 1.96*SEl))
}

g2 <- function(x, gpos){
  sapply(gpos, function(y) getOR2(y, df = x))
}

typesMethLevs <- lapply(types, function(t){
  rbind(filter(methMethLevels, Direction == t),
        filter(methMethLevels, Direction == "Non-significant")) %>%
    g2(cats) %>%
    as_tibble() %>%
    mutate(par = c("OR", "p.val", "ORlow", "ORhigh")) %>%
    gather("Region", "Value", 1:3) %>%
    mutate(Type = t)
})
combMethLevs <- Reduce(rbind, typesMethLevs)

methLevsPlot <- combMethLevs %>%
  spread(par, Value) %>%
  mutate(Type = ifelse(Type == "eQTM", "All", Type),
         Type = factor(Type, levels = c("All", "Inverse", "Positive"))) %>%
  ggplot(aes(x = Region, y = OR, fill = Type)) + 
  geom_bar(stat = "identity", position=position_dodge(), color = "black") + 
  geom_errorbar(position=position_dodge(.9), width=.25, aes(ymin = ORlow, ymax = ORhigh)) +
  scale_y_continuous(trans = "log2", 
                     breaks = scales::trans_breaks("log2", function(x) round(2^x, 2))) +
  geom_hline(yintercept = 1) +
  scale_x_discrete(name = "Categories of methylation levels", limits = cats, labels = tools::toTitleCase(cats)) +
  scale_fill_manual(name = "eCpG type", values = c("#999999", "#E69F00", "#009E73"),
                    labels = c("All", "Inverse", "Positive")) +
  theme_bw() 

png("paper/CpGEnrich_methLevels.png", width = 3000, height = 2000, res = 300)
methLevsPlot
dev.off()


methMethLevels.rel <- as_tibble(methyAnnot) %>%
  mutate(CpG = Name) %>%
  select(CpG, median_cat) %>%
  right_join(CpGsSum, by = "CpG") %>%
  filter(!is.na(Reliability) & Reliability >= 0.4) %>%
  mutate(Direction = factor(Direction, levels = c("Inverse", "Positive", "Both", "Non-significant"))) %>%
  group_by(Direction, median_cat) %>%
  summarize(n = n()) %>%
  spread(median_cat, n) %>%
  ungroup() %>%
  rbind(filter(., Direction != "Non-significant") %>% 
          summarize_all(fsum)) %>%
  mutate(Direction = as.character(Direction),
         Direction = ifelse(is.na(Direction), "eQTM", Direction),
         tot = low + medium + high)

typesMethLevs.rel <- lapply(types, function(t){
  rbind(filter(methMethLevels.rel, Direction == t),
        filter(methMethLevels.rel, Direction == "Non-significant")) %>%
    g2(cats) %>%
    as_tibble() %>%
    mutate(par = c("OR", "p.val", "ORlow", "ORhigh")) %>%
    gather("Region", "Value", 1:3) %>%
    mutate(Type = t)
})
methLevsPlot.rel <- Reduce(rbind, typesMethLevs.rel) %>%
  spread(par, Value) %>%
  mutate(Type = ifelse(Type == "eQTM", "All", Type),
         Type = factor(Type, levels = c("All", "Inverse", "Positive"))) %>%
  ggplot(aes(x = Region, y = OR, fill = Type)) + 
  geom_bar(stat = "identity", position=position_dodge(), color = "black") + 
  geom_errorbar(position=position_dodge(.9), width=.25, aes(ymin = ORlow, ymax = ORhigh)) +
  scale_y_continuous(trans = "log2", 
                     breaks = scales::trans_breaks("log2", function(x) round(2^x, 2))) +
  geom_hline(yintercept = 1) +
  scale_x_discrete(name = "Categories of methylation levels", limits = cats, labels = tools::toTitleCase(cats)) +
  scale_fill_manual(name = "eCpG type", values = c("#999999", "#E69F00", "#009E73"),
                    labels = c("All", "Inverse", "Positive")) +
  theme_bw() 

png("paper/CpGEnrich_methLevels_rel.png", width = 3000, height = 2000, res = 300)
methLevsPlot.rel
dev.off()

# CpG Islands ####
methIsland <- as_tibble(methyAnnot) %>%
  mutate(CpG = Name) %>%
  select(CpG, Relation_to_Island) %>%
  right_join(CpGsSum, by = "CpG") %>%
  mutate(Direction = factor(Direction, levels = c("Inverse", "Positive", "Both", "Non-significant"))) %>%
  group_by(Direction, Relation_to_Island) %>%
  summarize(n = n()) %>%
  spread(Relation_to_Island, n) %>%
  ungroup() %>%
  rbind(filter(., Direction != "Non-significant") %>% 
          summarize_all(fsum)) %>%
  mutate(Direction = as.character(Direction),
         Direction = ifelse(is.na(Direction), "eQTM", Direction),
         tot = Island + N_Shelf + N_Shore + OpenSea + S_Shelf + S_Shore)

  
islandStates <- c("N_Shelf", "N_Shore", "Island", "S_Shore", "S_Shelf", "OpenSea")


typesIsland <- lapply(types, function(t){
    rbind(filter(methIsland, Direction == t),
          filter(methIsland, Direction == "Non-significant")) %>%
    g2(islandStates) %>%
    as_tibble() %>%
    mutate(par = c("OR", "p.val", "ORlow", "ORhigh")) %>%
    gather("Region", "Value", 1:length(islandStates)) %>%
    mutate(Type = t)
})
combIsland <- Reduce(rbind, typesIsland)

cpgPosPlot <- combIsland %>%
  spread(par, Value) %>%
  mutate(Type = ifelse(Type == "eQTM", "All", Type),
         Type = factor(Type, levels = c("All", "Inverse", "Positive"))) %>%
  ggplot(aes(x = Region, y = OR, fill = Type)) + 
  geom_bar(stat = "identity", position=position_dodge(), color = "black") + 
  geom_errorbar(position=position_dodge(.9), width=.25, aes(ymin = ORlow, ymax = ORhigh)) +
  scale_y_continuous(trans = "log2", 
                     breaks = scales::trans_breaks("log2", function(x) round(2^x, 2))) +
  geom_hline(yintercept = 1) +
  scale_x_discrete(name = "CpG island relative position", limits = islandStates) +
  scale_fill_manual(name = "eCpG type", values = c("#999999", "#E69F00", "#009E73"),
                    labels = c("All", "Inverse", "Positive")) +
  theme_bw() +
  theme(legend.position = "none")

png("paper/CpGEnrichIsland.png", width = 3000, height = 2000, res = 300)
cpgPosPlot
dev.off()

methIsland.rel <- as_tibble(methyAnnot) %>%
  mutate(CpG = Name) %>%
  select(CpG, Relation_to_Island) %>%
  right_join(CpGsSum, by = "CpG") %>%
  filter(!is.na(Reliability) & Reliability >= 0.4) %>%
  mutate(Direction = factor(Direction, levels = c("Inverse", "Positive", "Both", "Non-significant"))) %>%
  group_by(Direction, Relation_to_Island) %>%
  summarize(n = n()) %>%
  spread(Relation_to_Island, n) %>%
  ungroup() %>%
  rbind(filter(., Direction != "Non-significant") %>% 
          summarize_all(fsum)) %>%
  mutate(Direction = as.character(Direction),
         Direction = ifelse(is.na(Direction), "eQTM", Direction),
         tot = Island + N_Shelf + N_Shore + OpenSea + S_Shelf + S_Shore)


typesIsland.rel <- lapply(types, function(t){
  rbind(filter(methIsland.rel, Direction == t),
        filter(methIsland.rel, Direction == "Non-significant")) %>%
    g2(islandStates) %>%
    as_tibble() %>%
    mutate(par = c("OR", "p.val", "ORlow", "ORhigh")) %>%
    gather("Region", "Value", 1:length(islandStates)) %>%
    mutate(Type = t)
})
cpgPosPlot.rel <- Reduce(rbind, typesIsland.rel) %>%
  spread(par, Value) %>%
  mutate(Type = ifelse(Type == "eQTM", "All", Type),
         Type = factor(Type, levels = c("All", "Inverse", "Positive"))) %>%
  ggplot(aes(x = Region, y = OR, fill = Type)) + 
  geom_bar(stat = "identity", position=position_dodge(), color = "black") + 
  geom_errorbar(position=position_dodge(.9), width=.25, aes(ymin = ORlow, ymax = ORhigh)) +
  scale_y_continuous(trans = "log2", 
                     breaks = scales::trans_breaks("log2", function(x) round(2^x, 2))) +
  geom_hline(yintercept = 1) +
  scale_x_discrete(name = "CpG island relative position", limits = islandStates) +
  scale_fill_manual(name = "eCpG type", values = c("#999999", "#E69F00", "#009E73"),
                    labels = c("All", "Inverse", "Positive")) +
  theme_bw() 

png("paper/CpGEnrichIsland_rel.png", width = 3000, height = 2000, res = 300)
cpgPosPlot.rel
dev.off()


# Chromatin states ####
chromStates <- c("TssA", "TssAFlnk", "TxFlnk", "TxWk", "Tx", "EnhG", "Enh",
                 "ZNF.Rpts", "Het", "TssBiv", "BivFlnk", "EnhBiv", "ReprPC",
                 "ReprPCWk", "Quies")
sum2 <- function(x) sum(!x)


methChromStIni <- as_tibble(methyAnnot) %>%
  mutate(CpG = Name) %>%
  select(CpG, eval(chromStates)) %>%
  right_join(CpGsSum, by = "CpG") %>%
  mutate(Direction = ifelse(Type == "Non-significant", "non-eQTM", "eQTM")) %>%
  group_by(Direction) %>%
  filter(!is.na(TssA)) %>%
  summarize_at(chromStates, list(sum = sum, sum2 = sum2, prop = mean))

methChromSt <- as_tibble(methyAnnot) %>%
  mutate(CpG = Name) %>%
  select(CpG, eval(chromStates)) %>%
  right_join(CpGsSum, by = "CpG") %>%
  mutate(Direction = factor(Direction, levels = c("Inverse", "Positive", "Both", "Non-significant"))) %>%
  group_by(Direction) %>%
  filter(!is.na(TssA)) %>%
  summarize_at(chromStates, list(sum = sum, sum2 = sum2, prop = mean))  %>%
  filter(Direction != "Non-significant") %>%
  rbind(methChromStIni)


typesChromSt <- lapply(types, function(t){
  rbind(filter(methChromSt, Direction == t),
        filter(methChromSt, Direction == "non-eQTM")) %>%
    select(ends_with("sum"), ends_with("sum2")) %>%
    g(chromStates, cols = c("_sum", "_sum2")) %>%
    as_tibble() %>%
    mutate(par = c("OR", "p.val", "ORlow", "ORhigh")) %>%
    gather("Region", "Value", 1:15) %>%
    mutate(Type = t)
})
combChromSt <- Reduce(rbind, typesChromSt)

chromStatesPlot <- combChromSt %>%
  spread(par, Value) %>%
  mutate(Type = ifelse(Type == "eQTM", "All", Type),
         Type = factor(Type, levels = c("All", "Inverse", "Positive")),
         Group = factor(ifelse(Region %in% c("TssA", "TssAFlnk"), "TssProxProm",
                               ifelse(Region %in% c("Tx", "TxWk"), "ActTrans", 
                                      ifelse(Region %in% c("Enh", "EnhG"), "Enhancer", 
                                             ifelse(Region %in% c("TssBiv", "BivFlnk", "EnhBiv"), "BivReg", 
                                                    ifelse(Region %in% c("ReprPC", "ReprPCWk"), "ReprPoly", Region)
                                             )
                                      )
                               )
         ), 
         levels = c("TssProxProm", "TxFlnk", "ActTrans", "Enhancer", "ZNF.Rpts", "BivFlnk", "BivReg", "Het", "ReprPoly", "Quies")
         )
  ) %>%
  ggplot(aes(x = Region, y = OR, fill = Type)) + 
  geom_bar(stat = "identity", position=position_dodge(), color = "black") + 
  geom_errorbar(position=position_dodge(.9), width=.25, aes(ymin = ORlow, ymax = ORhigh)) +
  scale_y_continuous(trans = "log2", 
                     breaks = scales::trans_breaks("log2", function(x) round(2^x, 2))) +
  geom_hline(yintercept = 1) +
  scale_x_discrete(name = "ROADMAP chromatin states") +
  scale_fill_manual(name = "eCpG type", values = c("#999999", "#E69F00", "#009E73"),
                    labels = c("All", "Inverse", "Positive")) +
  facet_wrap(~ Group, scales = "free_x") +
  theme_bw() +
  theme(legend.position = "none")

png("paper/CpGEnrichChromStates.png", width = 3000, height = 2000, res = 300)
chromStatesPlot
dev.off()


methChromStIni.rel <- as_tibble(methyAnnot) %>%
  mutate(CpG = Name) %>%
  select(CpG, eval(chromStates)) %>%
  right_join(CpGsSum, by = "CpG") %>%
  filter(!is.na(Reliability) & Reliability >= 0.4) %>%
  mutate(Direction = ifelse(Type == "Non-significant", "non-eQTM", "eQTM")) %>%
  group_by(Direction) %>%
  filter(!is.na(TssA)) %>%
  summarize_at(chromStates, list(sum = sum, sum2 = sum2, prop = mean))

methChromSt.rel <- as_tibble(methyAnnot) %>%
  mutate(CpG = Name) %>%
  select(CpG, eval(chromStates)) %>%
  right_join(CpGsSum, by = "CpG") %>%
  filter(!is.na(Reliability) & Reliability >= 0.4) %>%
  mutate(Direction = factor(Direction, levels = c("Inverse", "Positive", "Both", "Non-significant"))) %>%
  group_by(Direction) %>%
  filter(!is.na(TssA)) %>%
  summarize_at(chromStates, list(sum = sum, sum2 = sum2, prop = mean))  %>%
  filter(Direction != "Non-significant") %>%
  rbind(methChromStIni.rel)


typesChromSt.rel <- lapply(types, function(t){
  rbind(filter(methChromSt.rel, Direction == t),
        filter(methChromSt.rel, Direction == "non-eQTM")) %>%
    select(ends_with("sum"), ends_with("sum2")) %>%
    g(chromStates, cols = c("_sum", "_sum2")) %>%
    as_tibble() %>%
    mutate(par = c("OR", "p.val", "ORlow", "ORhigh")) %>%
    gather("Region", "Value", 1:15) %>%
    mutate(Type = t)
})
chromStatesPlot.rel <- Reduce(rbind, typesChromSt.rel) %>%
  spread(par, Value) %>%
  mutate(Type = ifelse(Type == "eQTM", "All", Type),
         Type = factor(Type, levels = c("All", "Inverse", "Positive")),
         Group = factor(ifelse(Region %in% c("TssA", "TssAFlnk"), "TssProxProm",
                               ifelse(Region %in% c("Tx", "TxWk"), "ActTrans", 
                                      ifelse(Region %in% c("Enh", "EnhG"), "Enhancer", 
                                             ifelse(Region %in% c("TssBiv", "BivFlnk", "EnhBiv"), "BivReg", 
                                                    ifelse(Region %in% c("ReprPC", "ReprPCWk"), "ReprPoly", Region)
                                             )
                                      )
                               )
         ), 
         levels = c("TssProxProm", "TxFlnk", "ActTrans", "Enhancer", "ZNF.Rpts", "BivFlnk", "BivReg", "Het", "ReprPoly", "Quies")
         )
  ) %>%
  ggplot(aes(x = Region, y = OR, fill = Type)) + 
  geom_bar(stat = "identity", position=position_dodge(), color = "black") + 
  geom_errorbar(position=position_dodge(.9), width=.25, aes(ymin = ORlow, ymax = ORhigh)) +
  scale_y_continuous(trans = "log2", 
                     breaks = scales::trans_breaks("log2", function(x) round(2^x, 2))) +
  geom_hline(yintercept = 1) +
  scale_x_discrete(name = "ROADMAP chromatin states") +
  scale_fill_manual(name = "eCpG type", values = c("#999999", "#E69F00", "#009E73"),
                    labels = c("All", "Inverse", "Positive")) +
  facet_wrap(~ Group, scales = "free_x") +
  theme_bw() 

png("paper/CpGEnrichChromStates_rel.png", width = 3000, height = 2000, res = 300)
chromStatesPlot.rel
dev.off()

### Combined plot
png("paper/enrich_combined_reliable.png", width = 3500, height = 3000, res = 300)
plot_grid(
  plot_grid(cpgPosPlot.rel, methLevsPlot.rel, labels = c("A", "C"), nrow = 1),
  chromStatesPlot.rel, 
  labels = c("", "B"), ncol = 1, rel_heights = c(1, 2)
)
dev.off()

title <- ggdraw() + 
  draw_label(
    "Enrichment for regulatory elements",
    fontface = 'bold')
png("paper/enrich_combined.png", width = 2200, height = 1800, res = 300)
plot_grid(
  title,
  plot_grid(cpgPosPlot, methLevsPlot, labels = c("A", "C"), nrow = 1),
  chromStatesPlot, 
  labels = c("", "", "B"), ncol = 1, rel_heights = c(0.1, 1, 2)
)
dev.off()

# Overlap with literature ####
#### EWAS Catalog ####
ewasdf <- read.delim(gzfile("data/EWAS_Catalog_03-07-2019.txt.gz"))

## Restrict to results in blood and European
ewasdfF <- ewasdf %>%
  filter(grepl("Whole|Peripheral", Tissue)) %>%
  filter(!N_EUR == "-" & (N_EAS != "-" | N_SAS != "-" | N_AFR != "-" | N_AMR != "-" | N_OTH == "-"))

ewasCatSum <- ewasdfF %>%
  dplyr::select(CpG) %>%
  mutate(Present = "Present") %>%
  distinct() %>%
  right_join(CpGsSum, by = "CpG") %>%
  mutate(Present = ifelse(is.na(Present), "Absent", Present)) %>%
  as_tibble() 


catalCpGs <- unique(ewasdfF$CpG)
CpGsSum %>%
  mutate(mQTL = CpG %in% catalCpGs,
         Type = ifelse(Combined != "Non-significant", "Significant", "Non-significant")) %>%
  group_by(Type) %>%
  summarize(n = sum(mQTL),
            tot = n(),
            prop = mean(mQTL))

ewasCatSum2 <-  ewasCatSum %>% 
  group_by(CpG) %>% 
  group_by(Present, Direction) %>%
  summarize(n = n()) %>%
  spread(Present, n) %>%
  mutate(Direction = factor(Direction)) %>%
  rbind(filter(., Direction != "Non-significant") %>% 
          summarize_all(fsum))  %>%
  mutate(Direction = as.character(Direction)) %>%
  select(Direction, Present, Absent)

typesCatal <- lapply(types, function(t){
  rbind(filter(ewasCatSum2, Direction == t),
        filter(ewasCatSum2, Direction == "Non-significant")) %>%
    getOR(cols = 2:3) %>%
    as_tibble() %>%
    mutate(par = c("OR", "p.val", "ORlow", "ORhigh")) %>%
    mutate(Type = t, 
           dataset = "EWAS Catalog" )
})
combCatal <- Reduce(rbind, typesCatal)


catal_meth <- as_tibble(methyAnnot) %>%
  mutate(CpG = Name) %>%
  select(CpG, median) %>%
  right_join(ewasCatSum, by = "CpG") %>%
  mutate(Significant = ifelse(Type == "Non-significant", "Non-eQTM", "eQTM")) %>%
  ggplot(aes(x = Present, y = median, fill = Significant)) +
  geom_violin() +
  scale_x_discrete(name = "Presence in EWAS catalogue") +
  scale_y_continuous(name = "Median methylation") +
  scale_fill_manual(name = "", values = c("#999999", "#FFFFFF")) +
  theme_bw()

ewasCatSum2.rel <-  ewasCatSum %>% 
  filter(!is.na(Reliability) & Reliability >= 0.4) %>%
  group_by(CpG) %>% 
  group_by(Present, Direction) %>%
  summarize(n = n()) %>%
  spread(Present, n) %>%
  mutate(Direction = factor(Direction)) %>%
  rbind(filter(., Direction != "Non-significant") %>% 
          summarize_all(fsum))  %>%
  mutate(Direction = as.character(Direction)) %>%
  select(Direction, Present, Absent)

typesCatal.rel <- lapply(types, function(t){
  rbind(filter(ewasCatSum2.rel, Direction == t),
        filter(ewasCatSum2.rel, Direction == "Non-significant")) %>%
    getOR(cols = 2:3) %>%
    as_tibble() %>%
    mutate(par = c("OR", "p.val", "ORlow", "ORhigh")) %>%
    mutate(Type = t, 
           dataset = "EWAS Catalog" )
})
combCatal.rel <- Reduce(rbind, typesCatal.rel)


## EWAS Atlas #### 
## Download time: 27/11/2019
atAssoc <- read.delim("data/EWAS_Atlas_associations.tsv")
atStudy <- read.delim("data/EWAS_Atlas_studies.tsv")
atCohort <- read.delim("data/EWAS_Atlas_cohorts.tsv")

## Combined traits (web page does not allow to download all files)
# traitsL <- lapply(1:5, function(x) read.csv(paste0("data/tableExport", x, ".csv")))
# traits <- Reduce(rbind, traitsL)
traits <- read.csv("data/traitsTable.csv")

colnames(atStudy)[1] <- "Study.id"
colnames(atCohort)[c(2, 4, 11, 15)] <- c("Study.id", "Array", "Source", "Ancestry")
atlasDF <- atStudy %>%
  select(Study.id) %>%
  full_join(select(atCohort, Study.id, Array, Source, Ancestry), by = "Study.id") %>% 
  left_join(atAssoc, ., by ="Study.id") %>%
  as_tibble() %>%
  distinct()

## Select Europeans and blood and add trait type
atlasDFsel <- atlasDF %>%
  filter(Ancestry %in% c("Not reported", "European") &
           Source %in% c("whole blood", "peripheral blood")) %>%
  left_join(select(traits, Trait, Type), by = "Trait")


atlasSum <- CpGsSum %>%
  mutate(Changed = CpG %in% atlasDFsel$Probe.id) %>%
  distinct() 
atlasSum %>%
  summarize(all = sum(Changed),
            eQTMs = sum(Changed & Type != "Non-significant")) %>%
  mutate(eQTMs_cat = eQTMs/all)

atlasSum2 <-  atlasSum %>% 
  mutate(Present = ifelse(Changed, "Present", "Absent")) %>%
  group_by(Present, Direction) %>%
  summarize(n = n()) %>%
  spread(Present, n) %>%
  mutate(Direction = factor(Direction)) %>%
  rbind(filter(., Direction != "Non-significant") %>% 
          summarize_all(fsum))  %>%
  mutate(Direction = as.character(Direction)) %>%
  select(Direction, Present, Absent)

  
typesAtlas <- lapply(types, function(t){
  rbind(filter(atlasSum2, Direction == t),
        filter(atlasSum2, Direction == "Non-significant")) %>%
    getOR(cols = 2:3) %>%
    as_tibble() %>%
    mutate(par = c("OR", "p.val", "ORlow", "ORhigh")) %>%
    mutate(Type = t, 
           dataset = "EWAS Atlas")
})
combAtlas <- Reduce(rbind, typesAtlas)


ewas_db <- combCatal %>%
  rbind(combAtlas) %>%
  spread(par, value) %>%
  mutate(Type = ifelse(Type == "eQTM", "All", Type),
         Type = factor(Type, levels = c("All", "Inverse", "Positive"))) %>%
  ggplot(aes(x = Type, y = OR, fill = Type)) + 
  geom_bar(stat = "identity", position=position_dodge(), color = "black") + 
  geom_errorbar(position=position_dodge(.9), width=.25, aes(ymin = ORlow, ymax = ORhigh)) +
  scale_y_continuous(trans = "log2", 
                     breaks = scales::trans_breaks("log2", function(x) round(2^x, 2))) +
  geom_hline(yintercept = 1) +
  scale_x_discrete(name = "eCpG type", labels = c("All", "Inverse", "Positive")) +
  scale_fill_manual(name = "eCpG type", values = c("#999999", "#E69F00", "#009E73")) +
  theme_bw() +
  theme(legend.position = "none",
        plot.title = element_text(hjust = 0.5)) +
  facet_grid(~  dataset) +
  ggtitle("All CpG probes")

png("paper/CpGEnrichEWASdbs.png", width = 3000, height = 1000, res = 300)
ewas_db
dev.off()


atlasSum2.rel <- atlasSum %>% 
  filter(!is.na(Reliability) & Reliability >= 0.4) %>%
  mutate(Present = ifelse(Changed, "Present", "Absent")) %>%
  group_by(Present, Direction) %>%
  summarize(n = n()) %>%
  spread(Present, n) %>%
  mutate(Direction = factor(Direction)) %>%
  rbind(filter(., Direction != "Non-significant") %>% 
          summarize_all(fsum))  %>%
  mutate(Direction = as.character(Direction)) %>%
  select(Direction, Present, Absent)


typesAtlas.rel <- lapply(types, function(t){
  rbind(filter(atlasSum2.rel, Direction == t),
        filter(atlasSum2.rel, Direction == "Non-significant")) %>%
    getOR(cols = 2:3) %>%
    as_tibble() %>%
    mutate(par = c("OR", "p.val", "ORlow", "ORhigh")) %>%
    mutate(Type = t, 
           dataset = "EWAS Atlas")
})
combAtlas.rel <- Reduce(rbind, typesAtlas.rel)


ewas_db.rel <- combCatal.rel %>%
  rbind(combAtlas.rel) %>%
  spread(par, value) %>%
  mutate(Type = ifelse(Type == "eQTM", "All", Type),
         Type = factor(Type, levels = c("All", "Inverse", "Positive"))) %>%
  ggplot(aes(x = Type, y = OR, fill = Type)) + 
  geom_bar(stat = "identity", position=position_dodge(), color = "black") + 
  geom_errorbar(position=position_dodge(.9), width=.25, aes(ymin = ORlow, ymax = ORhigh)) +
  scale_y_continuous(trans = "log2", 
                     breaks = scales::trans_breaks("log2", function(x) round(2^x, 2)),
                     limits = c(0.9, 3.5)) +
  geom_hline(yintercept = 1) +
  scale_x_discrete(name = "eCpG type", labels = c("All", "Inverse", "Positive")) +
  scale_fill_manual(name = "eCpG type", values = c("#999999", "#E69F00", "#009E73")) +
  theme_bw() +
  theme(legend.position = "none",
        plot.title = element_text(hjust = 0.5)) +
  facet_grid(~  dataset) +
  ggtitle("Reliable CpG probes")


png("paper/CpGEnrichEWASdbs_rel.png", width = 3000, height = 1000, res = 300)
ewas_db.rel
dev.off()

png("paper/CpGEnrichEWASdbs_panel.png", width = 3000, height = 2000, res = 300)
plot_grid(ewas_db, ewas_db.rel, labels = c("A", "B"), nrow = 2)
dev.off()
```
