## SupplementalCode for "Identification of autosomal cis expression quantitative trait methylation (cis eQTMs) in children’s blood": featuresDescriptives.R

```
###############################################################################
#' Create table with features descriptives
###############################################################################

library(dplyr)
library(S4Vectors)
library(minfi)
library(ggplot2)
library(cowplot)
library(matrixStats)

## Load data  ####
load("results/preprocessFiles/gexpAnnotation.Rdata")
load("results/preprocessFiles/methyAnnotation.Rdata")
load("results/preprocessFiles/allOverlaps.Rdata")
load("results/preprocessFiles/Methylation_GRSet.RData")
load("results/preprocessFiles/Expression_SE_residuals.RData")

## Total number CpGs
methyAnnot_aut <- methyAnnot[!methyAnnot$chr %in% c("chrX", "chrY"), ]
nrow(methyAnnot_aut)

## Total number TCs
sum(!expAnnot$seqname %in% c("chrX", "chrY"))

## Total number coding TCs
sum(expAnnot$Coding == "coding" & !expAnnot$seqname %in% c("chrX", "chrY"))

## Total number pairs
overDF_aut <- overDF[!grepl("X|Y", overDF$TC),]
nrow(overDF_aut)

## CpGs in pairs
length(unique(overDF_aut$CpG))
## CpGs not in pairs
sum(!methyAnnot$chr %in% c("chrX", "chrY")) - length(unique(overDF_aut$CpG))

## TCs in pairs
length(unique(overDF_aut$TC))

## TCs not in pairs
expAnnot_aut <- expAnnot[!expAnnot$seqname %in% c("chrX", "chrY"), ]
nrow(expAnnot_aut[!expAnnot_aut$probeset_id %in% overDF_aut$TC, ])
           

## Coding TCs not in pairs
subset(expAnnot_aut[!expAnnot_aut$probeset_id %in% overDF_aut$TC, ], Coding == "coding")

## Distribution CpGs per methylation level
prop.table(table(methyAnnot_aut$median_cat))*100

## Distribution CpGs per methylation variability
prop.table(table(methyAnnot_aut$variability))*100


## Proportion TCs with call rate == 100%
mean(expAnnot$CallRate > 90)
  # invariant   variant
# 44.0306   55.9694
# 


## Methylation distribution
betas <- as.vector(getBeta(gset))
png("paper/MethylationDistribution.png", width = 2000, height = 1500, res = 300)
betas %>%
  tibble::enframe(name = "a") %>%
  ggplot(aes(x = value)) + geom_density() + 
  theme_bw() + 
  scale_x_continuous("Methylation (betas)") +
  scale_y_continuous("", breaks = NULL) + 
  geom_vline(xintercept = c(0.3, 0.7), linetype = 2)
dev.off()

## CpGs/TC
CpG_plot <- overDF_aut %>%
  group_by(CpG) %>%
  summarize(n = n()) %>%
  ggplot(aes(x = n)) + geom_histogram(binwidth = 10) +
  scale_x_continuous("Genes per CpG", limits = c(0, 1250)) +
  scale_y_continuous("Number of CpGs")  +
  geom_vline(aes(xintercept = median(n))) +
  ggtitle("CpGs pairing distribution") +
  theme_bw() +
  theme(plot.title = element_text(hjust = 0.5))


TC_plot <- overDF_aut %>%
  group_by(TC) %>%
  summarize(n = n()) %>%
  ggplot(aes(x = n)) + geom_histogram(binwidth = 10) +
  scale_x_continuous("CpGs per Gene", limits = c(0, 1250)) +
  scale_y_continuous("Number of Genes")  +
  geom_vline(aes(xintercept = median(n))) +
  ggtitle("Genes pairing distribution") +
  theme_bw() +
  theme(plot.title = element_text(hjust = 0.5))


png("paper/CpGs_TC_distr.png", width = 2000, height = 1500, res = 300)
plot_grid(CpG_plot, TC_plot, labels = "AUTO", nrow = 2)
dev.off()

## CpGs per TC
overDF_aut %>%
  group_by(CpG) %>%
  summarize(n = n()) %>%
  summary()

## TCs per CpG
overDF_aut %>%
  group_by(TC) %>%
  summarize(n = n()) %>%
  summary()

## Correlation variability vs median methylation ####
png("paper/methy_Var_medianMeth.png", width = 2000, height = 1500, res = 300)
methyAnnot_aut %>% 
  data.frame() %>%
  ggplot(aes(x = median_cat, y = meth_range)) + 
  geom_boxplot() +
  scale_x_discrete(name = "Categories of methylation levels", labels = c("Low", "Medium", "High")) +
  scale_y_continuous(name = "Methylation range") +
  theme_bw()
dev.off()

summary(lm(meth_range ~ median_cat, methyAnnot_aut))

summary(lm(meth_range ~ factor(median_cat == "medium"), methyAnnot_aut))

## Expression distribution
png("paper/ExpressionDistribution.png", width = 2000, height = 1500, res = 300)
se %>%
  assay() %>%
  rowMedians() %>%
  tibble::enframe(name = "a") %>%
  ggplot(aes(x = value)) + geom_density() + 
  theme_bw() + 
  scale_x_continuous("Median Expression per TC") +
  scale_y_continuous("", breaks = NULL) 
dev.off()


## Expression Call Rate distribution
png("paper/ExpressionCRDistribution.png", width = 2000, height = 1500, res = 300)
expAnnot %>%
  ggplot(aes(x = CallRate)) + 
  geom_histogram() + 
  theme_bw() + 
  scale_x_continuous("TCs Call Rate") +
  scale_y_continuous("", breaks = NULL) 
dev.off()

## Correlation median Expression and Call Rate
medExp <- rowMedians(assay(se))
names(medExp) <- rownames(se)

if (identical(expAnnot$probeset_id, names(medExp))) {
  png("paper/ExpressionMedianCRDistribution.png", width = 2000, height = 1500, res = 300)
  data.frame(median = medExp, CallRate = expAnnot$CallRate) %>%
    mutate(CallRateCat = ifelse(CallRate > 90, ">90%", "<90%")) %>%
    ggplot(aes(x = CallRateCat, y = median)) +
    geom_boxplot() +
    scale_x_discrete(name = "TC Call Rate") +
    scale_y_continuous(name = "Median Expression") +
    theme_bw()
  dev.off()
  
  data.frame(median = medExp, CallRate = expAnnot$CallRate) %>%
    mutate(CallRateCat = ifelse(CallRate > 90, ">90%", "<90%")) %>%
    lm(median ~ CallRateCat, .) %>%
    summary()
}
# Call:
#   lm(formula = median ~ CallRateCat, data = .)
# 
# Residuals:
#   Min      1Q  Median      3Q     Max
# -3.5863 -0.6579 -0.2111  0.3318 15.8389
# 
# Coefficients:
#   Estimate Std. Error t value Pr(>|t|)
# (Intercept)      3.63635    0.01003   362.5   <2e-16 ***
#   CallRateCat>90%  2.35810    0.01479   159.4   <2e-16 ***
#   ---
#   Signif. codes:  0 ‘***’ 0.001 ‘**’ 0.01 ‘*’ 0.05 ‘.’ 0.1 ‘ ’ 1
# 
# Residual standard error: 1.816 on 60690 degrees of freedom
# Multiple R-squared:  0.2951,    Adjusted R-squared:  0.2951
# F-statistic: 2.541e+04 on 1 and 60690 DF,  p-value: < 2.2e-16
```
