## SupplementalCode for "Identification of autosomal cis expression quantitative trait methylation (cis eQTMs) in children’s blood": populationDescriptives.R

```
###############################################################################
#' Create table with population descriptives
###############################################################################

library(dplyr)
library(S4Vectors)
library(tidyr)

## Load data  ####
load("results/MethComBatExpResidualsCellAdj/pheno.RData")

getSum <- function(vec, type = "continuous"){
  if (type == "continuous"){
    c(median = median(vec, na.rm = TRUE), 
      range = quantile(vec, probs = c(0.25, 0.75), na.rm = TRUE))
  } else if (type == "categorical"){
    t <- table(vec)
    data.frame(names = names(t),  tab = as.vector(t), 
               props = as.vector(prop.table(t)))
  }
}

## Categorical variables ####
### Describe Cohort ("cohort") and Sex ("e3_sex")
catVars <- c("cohort", "e3_sex")
catVecs <- lapply(catVars, function(x) pheno[, x])
catSums <- lapply(catVecs, getSum, type = "categorical")

## Create table
catTab <- Reduce(rbind, catSums)
## Convert proportions to percentages
catTab$props <- paste0(sprintf("%.2f", catTab$props*100), "%")
write.table(catTab, file = "paper/PopDescrip_cat.txt", col.names = TRUE, 
            quote = FALSE, row.names = FALSE)

## Categorical variables ####
### Describe Age ("age_sample_years"), zBMI ("hs_zbmi_theano") and 
### Cell types ("NK", "Bcell", "CD4T", "CD8T", "Mono", "Neu")
contVars <- c("age_sample_years", "NK_6", "Bcell_6", "CD4T_6", "CD8T_6", "Mono_6", "Gran_6")
contVecs <- lapply(contVars, function(x) pheno[, x])
contSums <- lapply(contVecs, getSum, type = "continuous")

## Create table
contTab <- data.frame(Reduce(rbind, contSums))
## Convert proportions to percentages
contVals <- sprintf("%.3f (%.3f-%.3f)", contTab$median, contTab$range.25., contTab$range.75.)
contTab <- data.frame(names = contVars, vals = contVals)
write.table(contTab, file = "paper/PopDescrip_cont.txt", col.names = TRUE, 
            sep = "\t", quote = FALSE, row.names = FALSE)


## Statistics stratified by cohort
getSumCohort <- function(tab, varname, type = "continuous"){
  tab$var <- tab[, varname]
  df <- tab %>%
    data.frame() %>%
    group_by(cohort)
  if (type == "continuous"){
    df %>% 
      summarize(median = median(var, na.rm = TRUE), 
                range.25 = quantile(var, probs = 0.25, na.rm = TRUE),
                range.75 = quantile(var, probs = 0.75, na.rm = TRUE)) %>%
      mutate(val = sprintf("%.2f (%.2f-%.2f)", median, range.25, range.75)) %>%
      select(cohort, val)
    
  } else if (type == "categorical"){
    df %>%
      group_by(cohort, var) %>%
      summarize(N = n()) %>%
      group_by(cohort) %>%
      mutate(val = paste0(N, " (", round(N/sum(N)*100, 1), "%)")) %>%
      select(cohort, var, val) %>%
      spread(var, val)
  }
}
cohortCat <- getSumCohort(pheno, "e3_sex", type = "categorical")
cohortCont <- lapply(contVars, getSumCohort, tab = pheno) %>%
  Reduce(f = function(x, y) left_join(x, y, by = "cohort"), x = .) %>%
  left_join(cohortCat, ., by = "cohort")
colnames(cohortCont)[-c(1:3)] <- contVars
write.table(cohortCont, file = "paper/PopDescrip_cohort.txt", col.names = TRUE, 
            sep = "\t", quote = FALSE, row.names = FALSE)
```
