## SupplementalCode for "Identification of autosomal cis expression quantitative trait methylation (cis eQTMs) in children’s blood": README

This folder contains the scripts needed to reproduce the plots and tables of the manuscript.

createWebFiles.R: generates the files found in the web catalogue.
eQTM_catalogue.R: generates figures and tables describing main features of eQTMs
eQTM_genetics.R: generates figures and tables of the effect of genetics on eQTMs
eQTM_interpretation.R: generates figures and tables of the molecular enrichment of eQTMs.
featuresDescriptives.R: generates figures and tables of descriptives of DNA methylation and gene expression probes.
populationDescriptives.R: generates figures and tables of the populations descriptives.
README: this file
