## SupplementalCode for "Identification of autosomal cis expression quantitative trait methylation (cis eQTMs) in children’s blood": README

1.preprocessData: Scripts to prepare data for eQTM analysis.
2.generateCatalogue: Scripts to generate the eQTM catalogue.
3.DownstreamAnalyses: Scripts to reproduce the tables and figures of the manuscript.
README: this file.
